# Hilbert-Schmidt and Sobol sensitivity indices for static and time series Wnt signaling measurements in colorectal cancer - Part A

**DOI:** 10.1101/035519

**Authors:** shriprakash sinha

## Abstract

Ever since the accidental discovery of Wingless [Sharma R.P., Drosophila information service, 1973, 50, p 134], research in the field of Wnt signaling pathway has taken significant strides in wet lab experiments and various cancer clinical trials, augmented by recent developments in advanced computational modeling of the pathway. Information rich gene expression profiles reveal various aspects of the signaling pathway and help in studying different issues simultaneously. Hitherto, not many computational studies exist which incorporate the simultaneous study of these issues. This manuscript • explores the strength of contributing factors in the signaling pathway, • analyzes the existing causal relations among the inter/extracellular factors effecting the pathway based on prior biological knowledge and • investigates the deviations in fold changes in the recently found prevalence of psychophysical laws working in the pathway. To achieve this goal, local and global sensitivity analysis is conducted on the (non)linear responses between the factors obtained from static and time series expression profiles using the density (Hilbert-Schmidt Information Criterion) and variance (Sobol) based sensitivity indices. The results show the advantage of using density based indices over variance based indices mainly due to the former’s employment of distance measures & the kernel trick via Reproducing kernel Hilbert space (RKHS) that capture nonlinear relations among various intra/extracellular factors of the pathway in a higher dimensional space. In time series data, using these indices it is now possible to observe where in time, which factors get influenced & contribute to the pathway, as changes in concentration of the other factors are made. This synergy of prior biological knowledge, sensitivity analysis & representations in higher dimensional spaces can facilitate in time based administration of target therapeutic drugs & reveal hidden biological information within colorectal cancer samples. Code has been made available at Google drive on https://drive.google.com/folderview?id=0B7Kkv8wlhPU-Q2NBZGt1ZERrSVE&usp=sharing

## 1 Introduction

### 1.1 A short review

Sharma^1^’s accidental discovery of the Wingless played a pioneering role in the emergence of a widely expanding research field of the Wnt signaling pathway. A majority of the work has focused on issues related to • the discovery of genetic and epigenetic factors affecting the pathway (Thorstensen *et al.*^2^ & Baron and Kneissel^3^), • implications of mutations in the pathway and its dominant role on cancer and other diseases (Clevers^4^), • investigation into the pathway’s contribution towards embryo development (Sokol^5^), homeostasis (Pinto *et al.*^6^, Zhong *et al.*^7^) and apoptosis (Pećina-Šlaus^8^) and • safety and feasibility of drug design for the Wnt pathway (Kahn^9^, Garber^10^, Voronkov and Krauss^11^, Blagodatski *et al.*^12^ & Curtin and Lorenzi^13^). Approximately forty years after the discovery, important strides have been made in the research work involving several wet lab experiments and cancer clinical trials (Kahn^9^, Curtin and Lorenzi^13^) which have been augmented by the recent developments in the various advanced computational modeling techniques of the pathway. More recent informative reviews have touched on various issues related to the different types of the Wnt signaling pathway and have stressed not only the activation of the Wnt signaling pathway via the Wnt proteins (Rao and Kühl^14^) but also the on the secretion mechanism that plays a major role in the initiation of the Wnt activity as a prelude (Yu and Virshup^15^).

The work in this paper investigates some of the current aspects of research regarding the pathway via sensitivity analysis while using static (Jiang *et al.*^16^) and time series (Gujral and MacBeath^17^) gene expression data retrieved from colorectal cancer samples.

### 1.2 Canonical Wnt signaling pathway

Before delving into the problem statement, a brief introduction to the Wnt pathway is given here. From the recent work of Sinha^19^, the canonical Wnt signaling pathway is a transduction mechanism that contributes to embryo development and controls homeostatic self renewal in several tissues (Clevers^4^). Somatic mutations in the pathway are known to be associated with cancer in different parts of the human body. Prominent among them is the colorectal cancer case (Gregorieff and Clevers^20^). In a succinct overview, the Wnt signaling pathway works when the Wnt ligand gets attached to the Frizzled(*FZD*)/*LRP* coreceptor complex. *FZD* may interact with the Dishevelled (*DVL*) causing phosphorylation. It is also thought that Wnts cause phosphorylation of the *LRP* via casein kinase 1 (*CK* 1) and kinase *GSK*3. These developments further lead to attraction of Axin which causes inhibition of the formation of the degradation complex. The degradation complex constitutes of *AXIN*, the *β-catenin* transportation complex *APC, CK*1 and *GSK*3. When the pathway is active the dissolution of the degradation complex leads to stabilization in the concentration of *β-catenin* in the cytoplasm. As *β-catenin* enters into the nucleus it displaces the *GROUCHO* and binds with transcription cell factor *TCF* thus instigating transcription of Wnt target genes. *GROUCHO* acts as lock on *TCF* and prevents the transcription of target genes which may induce cancer. In cases when the Wnt ligands are not captured by the coreceptor at the cell membrane, *AXIN* helps in formation of the degradation complex. The degradation complex phosphorylates *β-catenin* which is then recognized by *FBOX/WD* repeat protein *β-TRCP*. *β-TRCP* is a component of ubiquitin ligase complex that helps in ubiquitination of *β-catenin* thus marking it for degradation via the proteasome. Cartoons depicting the phenomena of Wnt being inactive and active are shown in figures 1(A) and 1(B), respectively.

**Fig. 1.**
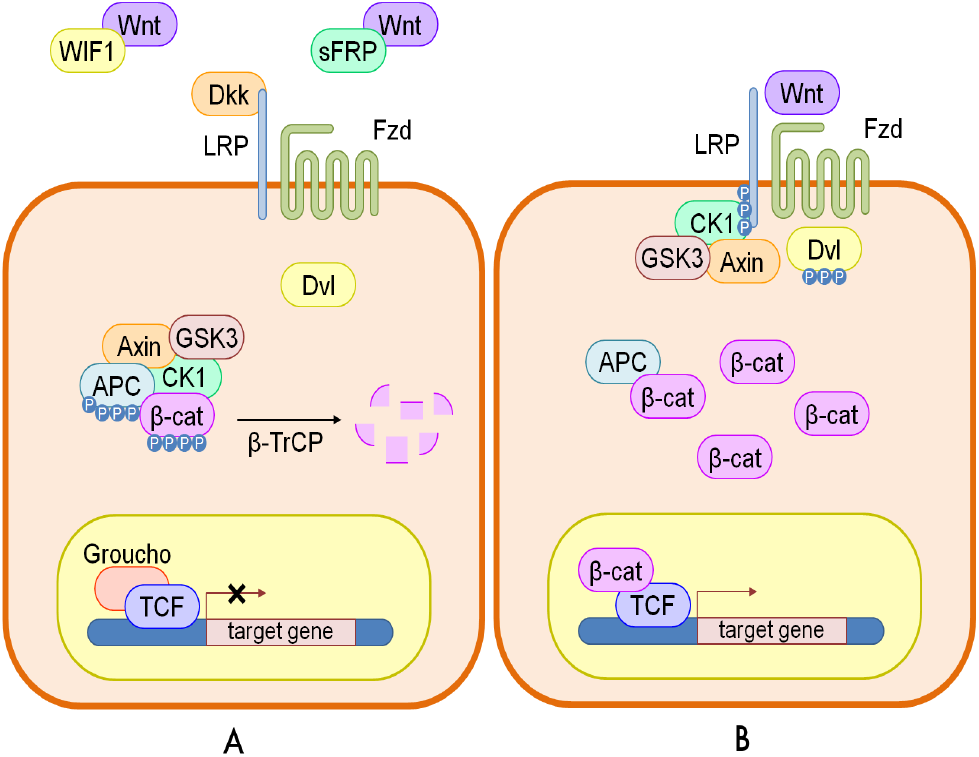
A cartoon of Wnt signaling pathway contributed by Verhaegh *et al.*^18^. Part (A) represents the destruction of *β-catenin* leading to the inactivation of the Wnt target gene. Part (B) represents activation of Wnt target gene.

## 2 Problem statement & sensitivity analysis

Succinctly the endeavour is to address the following issues - • explore the strength of contributing factors in the signaling pathway, • analyse the existing causal relations among the inter/extracellular factors effecting the pathway based on prior biological knowledge and • investigate the significance of deviations in fold changes in the recently found prevalence of psychophysical laws working in the pathway in a multi-parameter setting. The issues related to • inference of hidden biological relations among the factors, that are yet to be discovered and • discovery of new causal relations using hypothesis testing, will be addressed in a subsequent manuscript. The current manuscript analyses the sensitivity indices for fold changes and deviations in fold changes in 17 different genes from a set of 74 genes as presented by Gujral and MacBeath^17^. An immediate followup of the manuscript is the analysis of the remaining^57^ genes which happens to the part B of this manuscript.

In order to address the above issues, sensitivity analysis (SA) is performed on either the datasets or results obtained from biologically inspired causal models. The reason for using these tools of sensitivity analysis is that they help in observing the behaviour of the output and the importance of the contributing input factors via a robust and an easy mathematical framework. In this manuscript both local and global SA methods are used. Where appropriate, a description of the biologically inspired causal models ensues before the analysis of results from these models. The approach taken here is that first a problem will be addressed and then the analysis of results and discussion ensues before working with the next issue.

### 2.1 Sensitivity analysis

Seminal work by Russian mathematician Sobol’^21^ lead to development as well as employment of SA methods to study various complex systems where it was tough to measure the contribution of various input parameters in the behaviour of the output. A recent unpublished review on the global SA methods by Iooss and Lemaître^22^ categorically delineates these methods with the following functionality • screening for sorting influential measures (Morris^23^ method, Group screening in Moon *et al.*^24^ & Dean and Lewis^25^, Iterated factorial design in Andres and Hajas^26^, Sequential bifurcation in Bettonvil and Kleijnen^27^ and Cotter^28^ design), • quantitative indicies for measuring the importance of contributing input factors in linear models (Christensen^29^, Saltelli *et al.*^30^, Helton and Davis^31^ and McKay *et al.*^32^) and nonlinear models (Homma and Saltelli^33^, Sobol^34^, Saltelli^35^, Saltelli *et al.*^36^, Saltelli *et al.*^37^, Cukier *et al.*^38^, Saltelli *et al.*^39^, & Tarantola *et al.*^40^ Saltelli *et al.*^41^, Janon *et al.*^42^, Owen^43^, Tissot and Prieur^44^, Da Veiga and Gamboa^45^, Archer *et al.*^46^, Tarantola *et al.*^47^, Saltelli *et al.*^41^ and Jansen^48^) and • exploring the model behaviour over a range on input values (Storlie and Helton^49^ and Da Veiga *et al.*^50^, Li *et al.*^51^ and Hajikolaei and Wang^52^). Iooss and Lemaître^22^ also provide various criteria in a flowchart for adapting a method or a combination of the methods for sensitivity analysis. Figure 2 shows the general flow of the mathematical formulation for computing the indices in the variance based Sobol method. The general idea is as follows - A model could be represented as a mathematical function with a multidimensional input vector where each element of a vector is an input factor. This function needs to be defined in a unit dimensional cube. Based on ANOVA decomposition, the function can then be broken down into *ƒ*_0_ and summands of different dimensions, if *ƒ*_0_ is a constant and integral of summands with respect to their own variables is 0. This implies that orthogonality follows in between two functions of different dimensions, if at least one of the variables is not repeated. By applying these properties, it is possible to show that the function can be written into a unique expansion. Next, assuming that the function is square integrable variances can be computed. The ratio of variance of a group of input factors to the variance of the total set of input factors constitute the sensitivity index of a particular group. Detailed derivation is present in the Appendix.

**Fig. 2.**
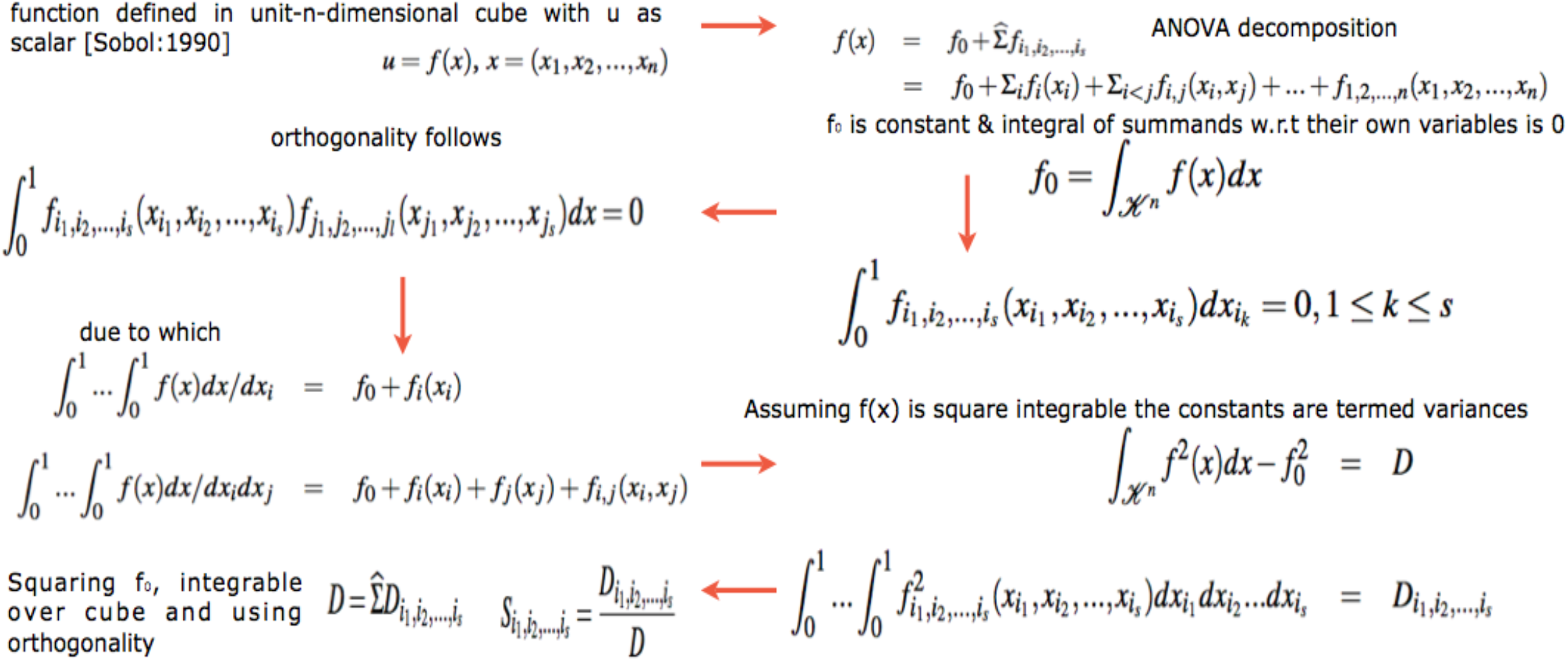
Computation of variance based sobol sensitivity indices. For detailed notations, see appendix.

Besides the above Sobol’^21^’s variance based indicies, more recent developments regarding new indicies based on density, derivative and goal-oriented can be found in Borgonovo^53^, Sobol and Kucherenko^54^ and Fort *et al.*^55^, respectively. In a latest development, Da Veiga^56^ propose new class of indicies based on density ratio estimation (Borgonovo^53^) that are special cases of dependence measures. This in turn helps in exploiting measures like distance correlation (Székely *et al.*^57^) and Hilbert-Schmidt independence criterion (Gretton *et al.*^58^) as new sensitivity indi-cies. The framework of these indicies is based on use of Csiszár *et al.*^59^ f-divergence, concept of dissimilarity measure and kernel trick Aizerman *et al.*^60^. Finally, Da Veiga^56^ propose feature selection as an alternative to screening methods in sensitivity analysis. The main issue with variance based indicies (Sobol’^21^) is that even though they capture importance information regarding the contribution of the input factors, they • do not handle multivariate random variables easily and • are only invariant under linear transformations. In comparison to these variance methods, the newly proposed indicies based on density estimations (Borgonovo^53^) and dependence measures are more robust. Figure 3 shows the general flow of the mathematical formulation for computing the indices in the density based HSIC method. The general idea is as follows - The sensitivity index is actually a distance correlation which incorporates the kernel based Hilbert-Schmidt Information Criterion between two input vectors in higher dimension. The criterion is nothing but the Hilbert-Schmidt norm of cross-covariance operator which generalizes the covariance matrix by representing higher order correlations between the input vectors through nonlinear kernels. For every operator and provided the sum converges, the Hilbert-Schmidt norm is the dot product of the orthonormal bases. For a finite dimensional input vectors, the Hilbert-Schmidt Information Criterion estimator is a trace of product of two kernel matrices (or the Gram matrices) with a centering matrix such that HSIC evalutes to a summation of different kernel values. Detailed derivation is present in the Appendix.

**Fig. 3.**
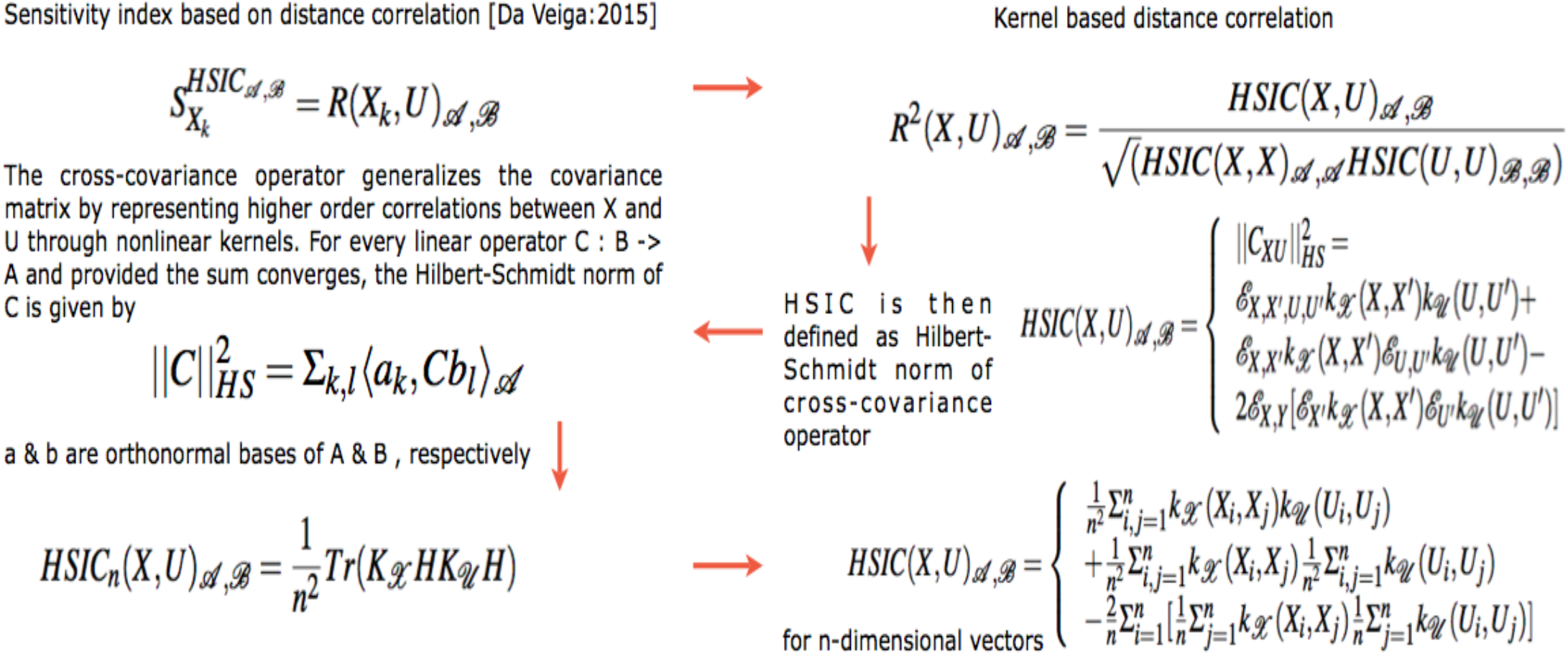
Computation of density based hsic sensitivity indices. For detailed notations, see appendix.

It is this strength of the kernel methods that HSIC is able to capture the deep nonlinearities in the biological data and provide reasonable information regarding the degree of influence of the involved factors within the pathway. Improvements in variance based methods also provide ways to cope with these nonlinearities but do not exploit the available strength of kernel methods. Results in the later sections provide experimental evidence for the same.

### 2.2 Relevance in systems biology

Recent efforts in systems biology to understand the importance of various factors apropos output behaviour has gained prominence. Sumner *et al.*^61^ compares the use of Sobol’^21^ variance based indices versus Morris^23^ screening method which uses a One-at-a-time (OAT) approach to analyse the sensitivity of *GSK*3 dynamics to uncertainty in an insulin signaling model. Similar efforts, but on different pathways can be found in Zheng and Rundell^62^ and Marino *et al.*^63^.

SA provides a way of analyzing various factors taking part in a biological phenomena and deals with the effects of these factors on the output of the biological system under consideration. Usually, the model equations are differential in nature with a set of inputs and the associated set of parameters that guide the output. SA helps in observing how the variance in these parameters and inputs leads to changes in the output behaviour. The goal of this manuscript is not to analyse differential equations and the parameters associated with it. Rather, the aim is to observe which input genotypic factors have greater contribution to observed phenotypic behaviour like a sample being normal or cancerous in both static and time series data. In this process, the effect of fold changes and deviations in fold changes in time is also considered for analysis in the light of the recently observed psychophysical laws acting downstream of the Wnt pathway (Goentoro and Kirschner^64^).

There are two approaches to sensitivity analysis. The first is the local sensitivity analysis in which if there is a required solution, then the sensitivity of a function apropos a set of variables is estimated via a partial derivative for a fixed point in the input space. In global sensitivity, the input solution is not specified. This implies that the model function lies inside a cube and the sensitivity indices are regarded as tools for studying the model instead of the solution. The general form of g-function (as the model or output variable) is used to test the sensitivity of each of the input factor (i.e expression profile of each of the genes). This is mainly due to its non-linearity, non-monotonicity as well as the capacity to produce analytical sensitivity indices. The g-function takes the form

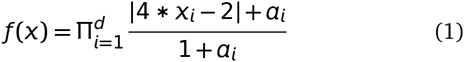

were, *d* is the total number of dimensions and *a*_*i*_ ≥ 0 are the indicators of importance of the input variable *x*_*i*_. Note that lower values of *a*_*i*_ indicate higher importance of *x*_*i*_. In our formulation, we randomly assign values of *x*_*i*_ € [0, 1]. For the static (time series) data *d* = 18(71) (factors affecting the pathway). Thus the expression profiles of the various genetic factors in the pathway are considered as input factors and the global analysis conducted. Note that in the predefined dataset, the working of the signaling pathway is governed by a preselected set of genes that affect the pathway. For comparison purpose, the local sensitivity analysis method is also used to study how the individual factor is behaving with respect to the remaining factors while working of the pathway is observed in terms of expression profiles of the various factors.

Finally, in context of Goentoro and Kirschner^64^’s work regarding the recent development of observation of Weber’s law working downstream of the pathway, it has been found that the law is governed by the ratio of the deviation in the input and the absolute input value. More importantly, it is these deviations in input that are of significance in studing such a phemomena. The current manuscript explores the sensitivity of deviation in the fold changes between measurements of fold changes at consecutive time points to explore in what duration of time, a particular factor is affecting the pathway in a major way. This has deeper implications in the fact that one is now able to observe when in time an intervention can be made or a gene be perturbed to study the behaviour of the pathway in tumorous cases. Thus sensitivity analysis of deviations in the mathematical formulation of the psychophysical law can lead to insights into the time period based influence of the involved factors in the pathway. Thus, both global and local anaylsis methods are employed to observe the entire behaviour of the pathway as well as the local behaviour of the input factors with respect to the other factors, respectively, via analysis of fold changes and deviations in fold changes, in time.

Given the range of estimators available for testing the sensitivity, it might be useful to list a few which are going to be employed in this research study. Also, a brief introduction into the fundamentals of the derivation of the three main indicies and the choice of sensitivity packages which are already available in literature, has been described in the Appendix.

## 3 Description of the dataset & design of experiments

Static data - A simple static dataset containing expression values measured for a few genes known to have important role in human colorectal cancer cases has been taken from Jiang *et al*^16^. Most of the expression values recorded are for genes that play a role in Wnt signaling pathway at an extracellular level and are known to have inhibitory affect on the Wnt pathway due to epigenetic factors. For each of the 24 normal mucosa and 24 human colorectal tumor cases, gene expression values were recorded for 14 genes belonging to the family of *SFRP, DKK, WIF*1 and *DACT*. Also, expression values of established Wnt pathway target genes like *LEF*1, *MYC, CD*44 and *CCND*1 were recorded per sample.

Time series data - Contrary to the static data described above, Gujral and MacBeath^17^ presents a bigger set of 71 Wnt-related gene expression values for 6 different times points over a range of 24-hour period using qPCR. The changes represent the fold-change in the expression levels of genes in 200 ng/mL *WNT*3*A*-stimulated HEK 293 cells in time relative to their levels in unstimulated, serum-starved cells at 0-hour. The data are the means of three biological replicates. Only genes whose mean transcript levels changed by more than two-fold at one or more time points were considered significant. Positive (negative) numbers represent up (down) -regulation.

Note that green (red) represents activation (repression) in the heat maps of data in Jiang *et al.*^16^ and Gujral and MacBeath^17^. Figures 4 and 5 represent the heat maps for the static and time series data respectively.

**Fig. 4.**
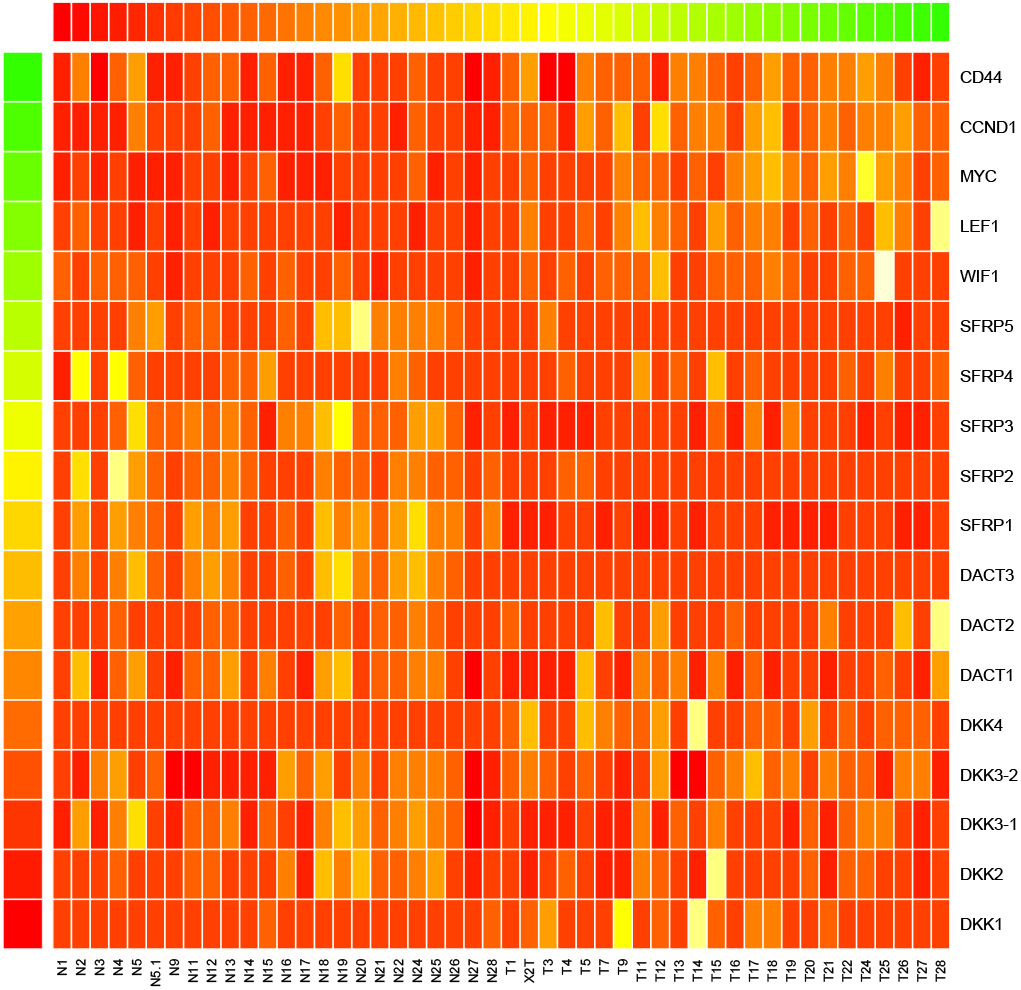
Heat map for gene expression values for each of the 24 normal mucosa and 24 human colorectal tumor cases from Jiang *et al.*^16^

**Fig. 5.**
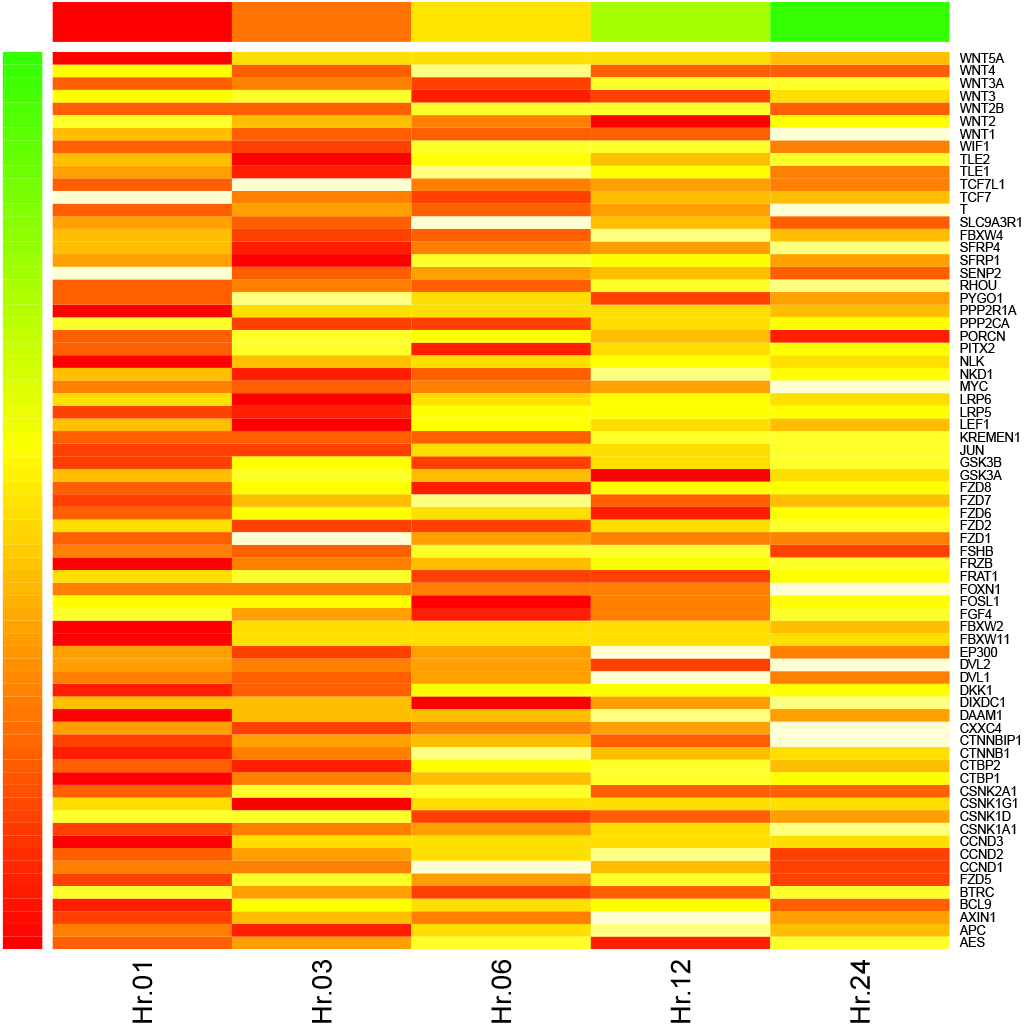
Heat map for gene expression values for 5 time points from Gujral and MacBeath^17^

general issues - • Here the input factors are the gene expression values for both normal and tumor cases in static data. For the case of time series data, the input factors are the fold change (deviations in fold change) expression values of genes at different time points (periods). Also, for the time series data, in the first experiment the analysis of a pair of the fold changes recorded at to different consecutive time points i.e 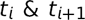 is done. In the second experiment, the analysis of a pair of deviations in fold changes recorded at 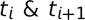 and 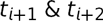. In this work, in both the static and the time series datasets, the analysis is done to study the entire model/pathway rather than find a particular solution to the model/pathway Thus global sensitivity analysis is employed. But the local sensitivity methods are used to observe and compare the affect of individual factors via 1^*st*^ order analysis w.r.t total order analysis (i.e global analysis). In such an experiment, the output is the sensitivity indices of the individual factors participating in the model. This is different from the general trend of observing the sensitivity of parameter values that affect the pathway based on differential equations that model a reaction. Thus the model/pathway is studied as a whole by observing the sensitivities of the individual factors.

• Static data - Note that the 24 normal and tumor cases are all different from each other. The 18 genes that are used to study in^16^ are the input factors and it is unlikely that there will be correlations between different patients. The phenotypic behaviour might be similar at a grander scale. Also, since the sampling number is very small for a network of this scale, large standard deviations can be observed in many results, especially when the Sobol method is used. But this is not the issue with the sampling number. By that analysis, large deviations are not observed in kernel based density methods. The deviations are more because of the fact that the nonlinearities are not captured in an efficient way in the variance based Sobol methods. Due to this, the resulting indi-cies have high variance in numerical value. For the same number of samplings, the kernel based methods don’t show high variance.

• Time series data - All the measurement data at each time point are generated by a normal distribution with fixed standard deviation of 0.005 plus a noise term. One might enquire as to how does this data generation match with the real experimental data? The kernel based density methods requires a distribution of data. The original experimental data of fold change was taken from each of the genes per time point. Gujral and MacBeath^17^ states that to determine the fold-change in gene expression induced by stimulation with Wnt3a, the normalized expression of each gene in the Wnt3a-stimulated sample was divided by the normalized expression of the same gene in the unstimulated sample. The qPCR data presented are mean of three biological replicates. By using a stringent margin of 0.005 and a noise term, the distribution of the data near the mean value is kept constricted. How much it deviates from the reality beyond the errors of measurement is not known to the author! Finally, 74 gene expression values are taken as input per time point for evaluting the sensitivity of each of the genetic factor that affect the model/pathway Again, one is not looking for a solution to the model in terms of good value for parameters but studying the degree of influence of each of the input factors that constitute the model/pathway

Design of experiments - The reported results will be based on scaled as well as unscaled datasets. For the static data, only the scaled results are reported. This is mainly due to the fact that the measurements vary in a wide range and due to this there is often an error in the computed estimated of these indices. The data for time series does not vary in a wide range and thus the results are reported for both the scaled and the non scaled versions. Total sensitivity indices and 1^*st*^ order indices will be used for sensitivity analysis. For addressing a biological question with known prior knowledge, the order of indices might be increased. While studying the interaction among the various genetic factors using static data, tumor samples are considered separated from normal samples. Bootstrapping without replicates on a smaller sample number is employed to generate estimates of indices which are then averaged. This takes into account the variance in the data and generates confidence bands for the indices. For the case of time series data, interactions among the contributing factors are studied by comparing (1) pairs of fold-changes at single time points and (2) pairs of deviations in fold changes between pairs of time points. Generation of distribution around measurements at single time points with added noise is done to estimate the indices.

## 4 Static data

To measure the strength of the contributing factors in the static dataset by Jiang *et al.*^16^, 1^*st*^ order and total sensitivity indices were generated. For each of the expression values of the genes recorded in the normal and tumor cases, the computation of the indices was done using bootstrapped samples in three different experiments each with a sample size of 8, 16 and 24, respectively. With only 24 samples in total, 20 bootstraps were generated for each set and the results were generated. From these replicates, the mean of the indices is reported along with the 95% confidence bands. Figure 6 represents the cartoon of the experimental setup followed to acheive the desired results. Note that plots of sensitivity indices have been relegated to Appendix.

**Fig. 6.**
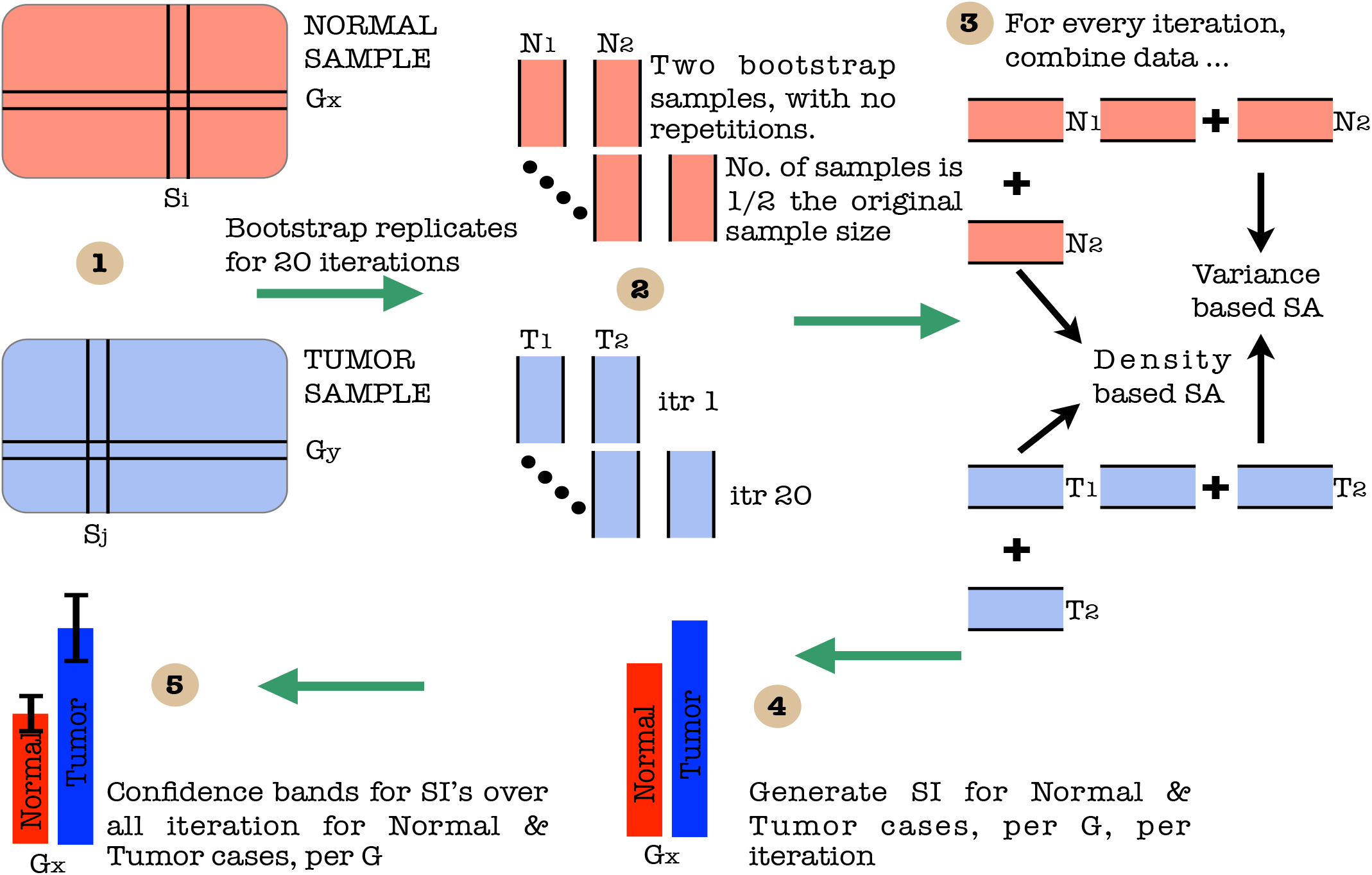
A cartoon of experimental setup. Step - (1) Segregation of data into normal and tumor cases. (2) Further data division per case and bootstrap sampling with no repetitions for different iterations. (3) Assembling bootstrapped data and application of SA methods. (4) Generation of SI’s for normal and tumor case per gene per iteration. (5) Generation of averaged SI and confidence bands per case per gene

Using the sensiFdiv, all indices are computed as positive and those nearing to zero indicate the contribution of a factor as independent from the behaviour under consideration. Here, while comparing the indices of the gene expression values for normal and tumor cases, it was found that most of the involved intra/extracellular factors had some degree of contribution in the normal case and almost negligible contribution in the tumor case (see figures 13, 15 and 16). Apart from the negative reading for the KL divergence 14 the interpretations remain the same. This implies that the basic Csiszár *et al.*^59^ f-divergence based indices might not capture the intrinsic genotypic effects for the normal and the tumorous cases. From the biological perspective, these graphs do not help in interpreting the strength of the contributions in normal and tumor cases. One might rank the indices for relative contributions, but this might not shed enough light on the how the factors are behaving in normal and tumor cases.

A more powerful way to analyse the contributions is the newly proposed HSIC based measures by Da Veiga^56^. These distances use the kernel trick which can capture intrinsic properties inherent in the recorded measurements by transforming the data into a higher dimensional space. Using these distances in sensiHSIC, it was found that the contributions of the various factors in the normal and the tumor cases vary drastically. This is shown in figures 17, 18 and 19. The laplace and the rbf kernels give more reliable sensitivity estimates for the involved factors than the linear kernel. Studying the results in figures 6 and 7 of Sinha^19^ based on prior biological knowledge encoded in the Bayesian network models along with the indices of aforementioned figures, it can be found that indices of the family of *DACT* – 1/2/3 show higher (lower) sensitivity in the normal (tumor) case where the activation (repression) happens. Again, of the *DACT* family, *DACT* – 1 has greater influence than *DACT* – 3 (than *DACT* – 2) based on the values of the sensitivity indices. These indices indicate the dependence of a factor on the output of the model characterized by the signaling being active (passive) in the normal (tumor) cases. 0(1) mean no (full) dependence of the output on the input factor. The laplace and the rbf kernels were found to give more consistent results than the linear kernel and the following description discusses the results from these kernels. For the *SFRP* family *SFRP* – 1/2/5 show higher (lower) sensitivity in normal (tumor) case where the activation (repression) happens (see figures 18 and 19). For *SFRP* – 3/4 the influence is higher (lower) in the tumor (normal) case. In all the three types of kernels, *WIF*1, *MYC* and *CCND*1 show stronger (weaker) influence of repression (activation) in the normal (tumor) case (see figures 18 and 19). *CD*44 showed variable influence while observing the normal and tumor cases. Sinha^19^ could not derive proper inferences for *LEF*1 but the sensitivity indices indicate that the influence of *LEF*1 in tumor samples to be higher than in normal samples. This points to the *LEF*1’*s* major role in tumor cases. Finally, for the family of *DKK*, *DKK*1 and *DKK*3 – 2 show similar behaviour of expression (repression) in normal (tumor cases) (see Sinha^19^). For the former, the prominence of the influence is shown in the higher (lower) sensitivity for tumor (normal) case. For the latter higher (lower) sensitivity was recorded for normal (tumor) case. This implies that the latter has more influential role in normal while the former has more influential role in tumor case. *DKK*3 – 1 was found to be expressed (repressed) in normal (tumor) and its dominant role is prominent from the higher bar sensitivity bar for normal than the tumor. Similar behavior of *DKK*2 was inferred by Sinha^19^ but the sensitivity indices point to varied results and thus a conclusion cannot be drawn. Note that greater the value of the sensitivity index, greater is an input factor’s contribution to the model.

**Fig. 7.**
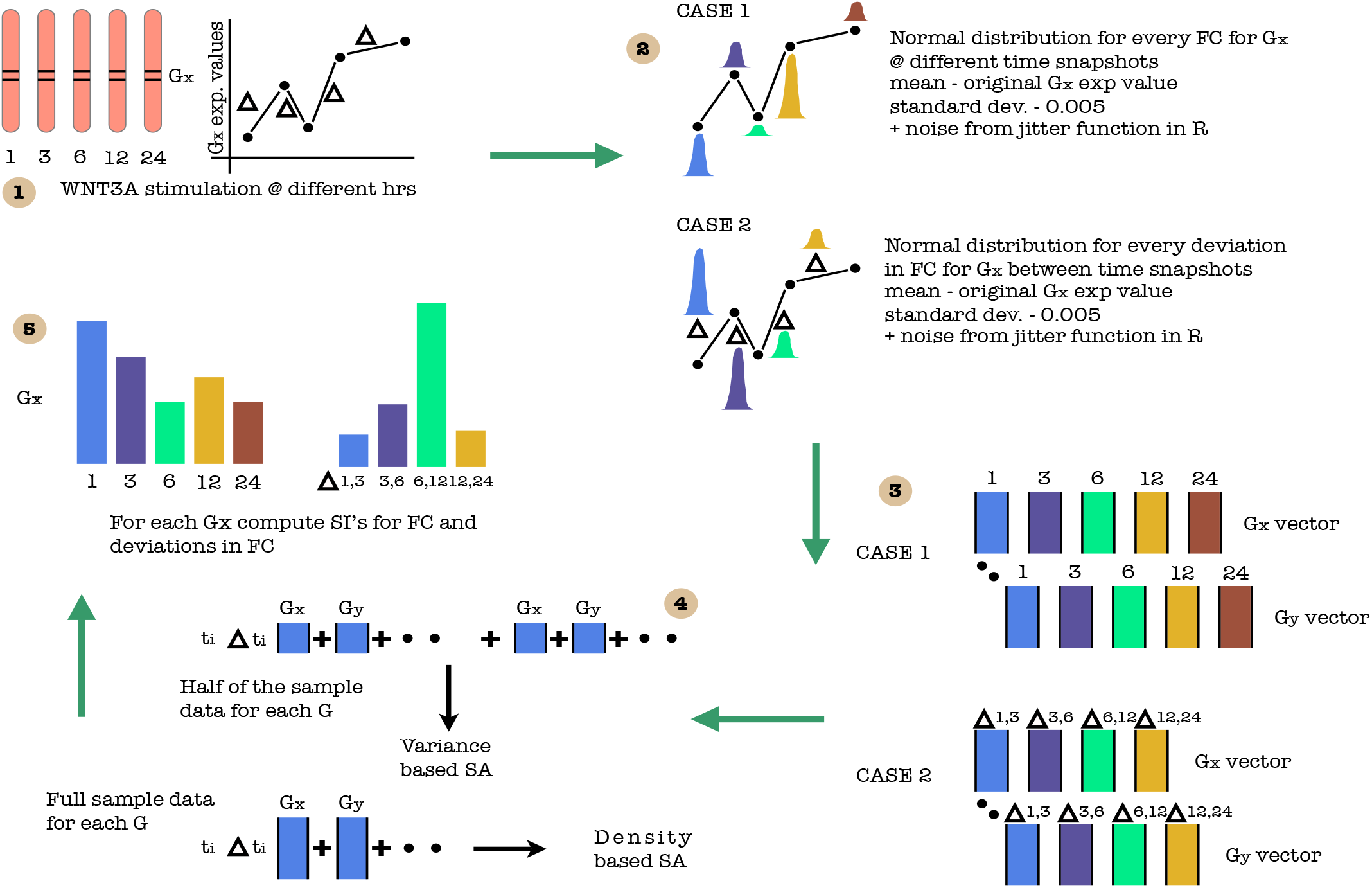
A cartoon of experimental setup. Step - (1) Time recordings of different gene expression values after WNT3A stimulation at different hrs. (2) Generation of normal distribution for every FC & ΔFC for Gx at & between different time snapshots, respectively. mean - original Gx exp value; standard dev - 0.005 + noise from jitter function in R (3) Generation of data set for FC & ΔFC. (4) Generation of samples for SA. (5) Compute SI for FC & ΔFC.

The first order indices generated by sobol functions implemented in sobol2002 (figure 20), sobol2007 (figure 21), sobol-jansen (figure 22), sobolmartinez (figure 23) and sobol (figure 24) do not point to significant dependencies of the input factors. This can be attributed to the fact that there are less number of samples that help in the estimation of the sensitivity indicies. Finally, the total order indices need to be investigated in the context of the first order indices. It can be observed, sobol2002 (figure 25) and sobol2007 (figure 25) give much better estimates than soboljansen (figure 27) and sobolmartinez (figure 28). Most importantly, it is the former two that closely match with the sensitivity indices estimated using the HSIC distance measures. Interpretations from sobol2002 (figure 25) and sobol2007 (figure 26) are the same as those described above using the laplace and the rbf kernels from density based HSIC measure.

In summary, the sensitivity indices confirm the inferred results in Sinha^19^ but do not help in inferring the causal relations using the static data. In combination with the results obtained from the Bayesian network models in Sinha^19^ it is possible to study the effect of the input factors for the pathway in both normal and tumor cases. The results of sensitivity indices indicate how much these factors influence the pathway in normal and tumor cases. Again, not all indices reveal important information. So users must be cautious of results and see which measures reveal information that are close to already established or computationally estimated biological facts. Here the density based sensitivity indices captured information more precisely than the variance based indices (except for the total order indices from sobol2002/7 which gave similar results as sensiHSIC). This is attributed to the analytical strength provided by the distance measures using the kernel trick via RKHS that capture nonlinear relations in higher dimensional space, more precisely. Finally, in a recent unpublished work by De Lozzo and Marrel^73^, it has been validated that the HSIC indices prove to be more sensitive to the global behaviour than the Sobol indices.

## 5 Time series data

Next, the analysis of the time series data is addressed using the sensitivity indices. There are two experiments that have been performed. First is related to the analysis of a pair of the fold changes recorded at to different consecutive time points i.e 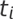 & 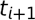. The second is related to the analysis of a pair of deviations in fold changes recorded at 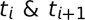 and 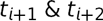. The former compares the measurements in time while the latter takes into account the deviations that happens in time. For each measurement at a time point a normal distribution was generated with original recorded value as the mean, a standard deviation of 0.005 and an added noise in the form of jitter (see function jitter in R langauge). For the time measurements of each of the genes recorded in Gujral and MacBeath^17^ an analysis of the sensitivity indices for both the scaled and the non-scaled data was done. Here the analysis for non-scaled data is presented. The reason for not presenting the scaled data is that the sample measurements did not vary drastically as found in the case of static data which caused troubles in the estimation of indices earlier. Another reason for not reporting the results on the scaled data is that the non-scaled ones present raw sensitive information which might be lost in scaling via normalization. Note that the authors use self organizing maps (SOM) to cluster data and use correlational analysis to derive their conclusions. In this work, the idea of clustering is abandoned and sensitivity indices are estimated for recorded factors participating in the pathway. Also the simple correlational analysis is dropped in favour of highly analytical kernel based distance measures. Figure 7 represents the experimental setup in a pictorial format.

Also, in a recent development, Goentoro and Kirschner^64^ point to two findings namely, • the robust fold changes of *β-catenin* and • the transcriptional machinery of the Wnt pathway depends on the fold changes in *β-catenin* instead of absolute levels of the same and some gene transcription networks must respond to fold changes in signals according to the Weber’s law in sensory physiology. The second study also carries a weight in the fact that due to the study of the deviations in the fold changes it is possible to check if the recently observed and reported natural psychophysical laws in the signaling pathway hold true or not. Finally, using the sensitivity indicies an effort is made to confirm the existing biological causal relations.

### 5.1 Analysis of fold changes at different time points

Lets begin with the gene *WNT*3*A* as changes in its concentration lead to recording of the measurements of the different genes by Gujral and MacBeath^17^. Of the list of genes recorded, the indices of the those which are influenced by the concentration of *WNT*3*A* are analysed. Next based on these confirmations and patterns of indices over time, conclusions for other enlisted genes are drawn. For the former list, the following genes *FZD*1, *FZD*2, *LEF*1, *TCF*7, *TCF*7*L*1, *LRP*6, *DVL*1, *SFRP*1, *SFRP*4, *CTBP*1, *CTBP*2, *PORCN, GSK*3*β*, *MYC*, *APC* and *CTNNB*1 are considered.

Figures 8 and 9 represent the indices computed over time. Columns represent the different kinds of indices computed while the rows show the respective genes. Each graph contains the sensitivity index computed at a particular time point (represented by a coloured bar). It should be observed from the aforementioned figures that the variants of the Sobol first order (FO) and the total order (TO) indices computed under different formulations were not very informative. This can be seen in graphs were some indices are negative and at some places the behaviour across time and genes remain the same. In contrast to this, the indices generated via the original Sobol function (under the column Sobol-SBL) as well as the sensiHSIC were found to be more reliable. Again, the rbf and laplace kernels under the HSIC formulations showed similar behaviour in comparison to the use of the linear kernel.

Gujral and MacBeath^17^ simulate the serum starved HEK293 cells with 200 ng/mL of *WNT*3*A* at different lengths of time. After the first hour (*t*_1_), (under HSIC-rbf/laplace) it was observed that the sensitivity of *WNT*3*A* was low (red bar). The maximum contribution of *WNT*3*A* can be recoreded after the 12^*th*^ stimulation. But due to increased stimulation by *WNT*3*A* later on, there is an increased sensitivity of *FZD*-1/2 as well as *LRP*6. The *FZD* or the frizzled family of 7-transmember protein (Ueno *et al.*^74^) works in tandem with *LRP*-5/6 as binding parameters for the Wnt ligands to initiate the Wnt signaling. Consistent with the findings of Holcombe *et al*.^75^ and Planutis *et al*.^76^, *FZD*1 was found to be expressed. But there is a fair decrease in the contribution of the same in the next two time frames i.e after 3^*rd*^ and the 6^*th*^ hour. The maximum contribution of *FZD*1 is found after the *WNT*3A simulation at 12^*th*^ hour. This probably points to repetitive involvement of *FZD* 1 after a certain period of time to initiate the working of the signaling pathway. *FZD*2 showed increasing significance in contribution after the first two time frames. The contribution drops significantly after the 3^*rd*^ simulation and gradually increases in the next two time frames. The repetitive behaviour is similar to *FZD*1, yet it’s role is not well studied as it appears to bind to both *WNT*3*A* which promotes *Wnt/beta-catenin* signaling and *WNT*5*A* which inhibits it as shown by Sato *et al.*^77^, respectively.

**Fig. 8.**
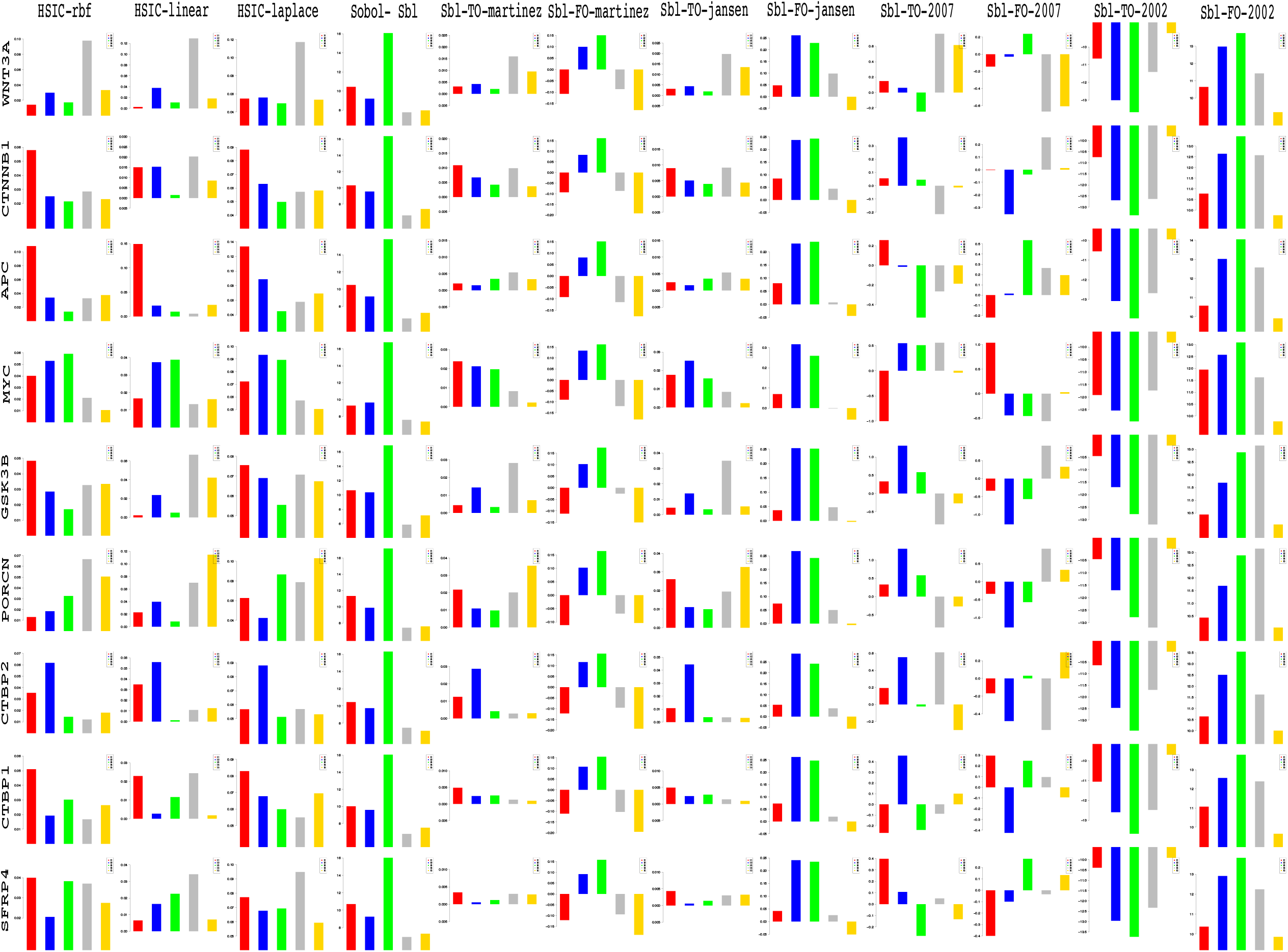
Column wise - methods to estimate sensitivity indices. Row wise - sensitivity indicies for each gene. For each graph, the bars represent sensitivity indices computed at t1 (red), t2 (blue), t3 (green), t4 (gray) and t5 (yellow). Indices were computed using non scaled time series data. TO - total order; FO - first order; SBL - Sobol

**Fig. 9.**
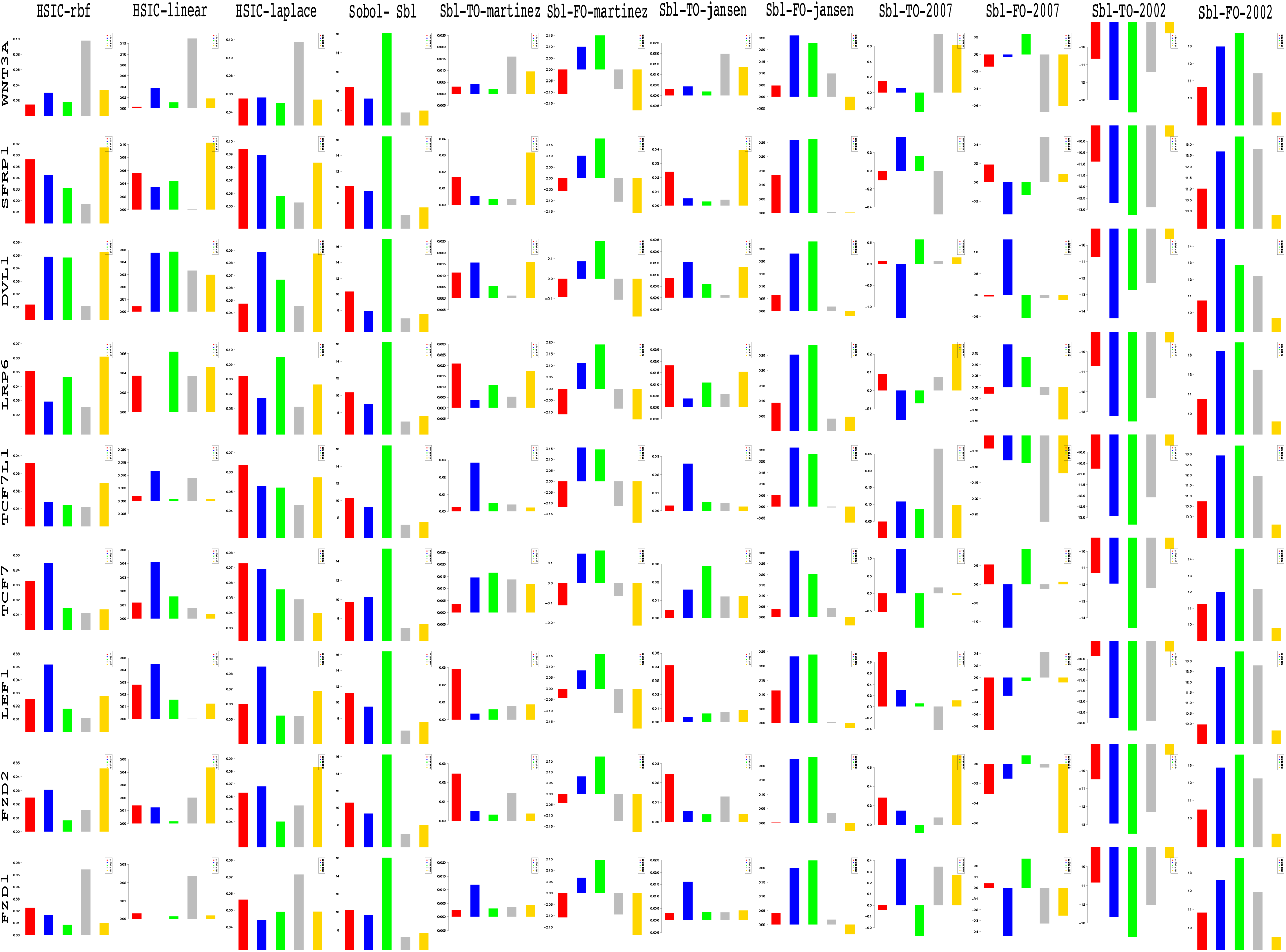
Column wise - methods to estimate sensitivity indices. Row wise - sensitivity indicies for each gene. For each graph, the bars represent sensitivity indices computed at t1 (red), t2 (blue), t3 (green), t4 (gray) and t5 (yellow). Indices were computed using non scaled time series data. TO - total order; FO - first order; SBL - Sobol

Klapholz-Brown *et al*.^78^ and Yokoyama *et al.*^79^ show that there is increased *β-catenin* due to *WNT*3*A* stimulation which is depicted by the increased sensitivity of *CTNNB*1 expression in one of the above mentioned figures. *MYC* (i.e *c – MYC*) is known to be over expressed in colorectal cancer cases mainly due to the activation of *TCF –* 4 transcription factor via intra nuclear binding of *β-catenin* (He *et al.*^80^), either by *APC* mutations (Korinek *et al.*^81^) or *β-catenin* mutations (Morin *et al.*^82^). The sensitivity of *MYC* increased monotonically but after the 6^*th*^ hour it dropped significantly. Probably *MYC* does not play important role at later stages. As found in Hino *et al*.^83^ and You *et al*.^84^, *DVL* family interacts with the frizzled *FDZ* members leading to disassembly of the *β-catenin* destruction complex and subsequent translocation of *β-catenin* to the nucleus. Development on *DVL* family have been extensively recorded in González-Sancho *et al*.^85^ and^86^, and significance of *DVL*1 in Taiwanese colorectal cancer in Huang *et al.*^87^. *DVL*1 shows a marked increase in sensitivity as the concentration of the *WNT*3*A* increases in time. This is supported by the fact that ligand binding at the membrane leads to formation of complex including *DVL*1, *FZD* and *AXIN.*

Negative regulators like *SFRP*4 were found to have lower sensitivity as *WNT*3*A* concentration increases, but remained constant for most period. Meanwhile the significance of Wnt antagonist *SFRP*1 (Galli *et al*.^88^, Suzuki *et al*.^89^ and Caldwell *et al*.^90^) decreases over the period as the concentration of *WNT*3*A* increases. Chinnadurai^91^ reviews the co-repressor ability of the *CTBP* family, while Hamada and Bienz^92^ shows *CTBP* as a binding factor that interacts with *APC* thus lowering the availability of free nuclear *β-catenin*. This interaction is further confirmed in the recent research work by Schneikert *et al*.^93^. As shown by Yokoyama *et al*.^79^ *CTBP*1 showed increased sensitivity with increased stimulation of *WNT*3*A* in the first hour. The latter stages show a decreased contribution of *CTBP*1 as the concentration of *WNT*3*A* was increased. This is in line with what Gujral and MacBeath^17^ show in their manuscript and indicate the lowering of the co-repressor effect of *CTBP* at later stages. On the other hand, *CTBP*2 showed reverse behaviour of sensitivity in comparison to *CTBP*1 across different time points. Increased significance of *CTBP*2 was observed in the first two time frames, i.e after 1^*st*^ and 3^*rd*^ hour of stimulation, followed by lower contribution to the pathway at the latter stages. In both cases, the diminishing co-repressive nature of *CTBP* in time is observed. Contrary to these finding, recent results in Patel *et al*.^94^ suggest that both *CTBP*1 and *CTBP*2 are up-regulated in colon cancer stem cells.

*PORCN* showed less sensitivity in the initial stages than in final stages indicating its importance in the contribution to Wnt secretion which is necessary for signaling (Willert and Nusse^95^). The sensitivity of *GSK*3*β* and *APC* decreased in time indicating the lowering of its significance in later stages due to no formation of the degradation complex. Activity of *TCF* gains greater prominence in the first and the second time frames after the initial *WNT*3*A* stimulation. This is in conjugation with the pattern showed by *CTBP*2. Regarding *TCF*7*L*2, the activity is observed to be maximum during the first time frame with decrease in the contribution in the later time frames.

Indicies for remaining 57 genes as well as analysis of the same will be presented in the following B part of this manuscript. Graphs for these 57 genes have been presented in figures 29 and 30 in Appendix.

### 5.2 The logarithmic psychophysical law

In a recent development, Goentoro and Kirschner^64^ point to two findings namely, • the robust fold changes of *β*-catenin and • the transcriptional machinery of the Wnt pathway depends on the fold changes in *β*-catenin instead of absolute levels of the same and some gene transcription networks must respond to fold changes in signals according to the Weber’s law in sensory physiology. In an unpublished work by Sinha^96^, preliminary analysis of results in Sinha^19^ shows that the variation in predictive behaviour of *β*-catenin based transcription complex conditional on gene evidences follows power and logarithmic psychophysical law crudely, implying deviations in output are proportional to increasing function of deviations in input and showing constancy for higher values of input. This relates to the work of Adler *et al*.^97^ on power and logarithmic law albeit at a coarse level. A description of these laws ensues before the analysis of the results.

Masin *et al*.^98^ states the Weber’s law as follows - Consider a sensation magnitude *γ* determined by a stimulus magnitude *β*. Fechner^99^ (vol 2, p. 9) used the symbol Δ*γ* to denote a just noticeable sensation increment, from *γ* to *γ* + Δ*γ*, and the symbol *Δβ* to denote the corresponding stimulus increment, from *β to β* + Δ*β*. Fechner^99^ (vol 1, p. 65) attributed to the German physiologist Ernst Heinrich Weber the empirical finding Weber^100^ that Δ*γ* remains constant when the relative stimulus increment 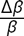 remains constant, and named this finding Weber’s law. Fechner^99^ (vol 2, p. 10) underlined that Weber’s law was empirical. ■

It has been found that Bernoulli’s principle (Bernoulli^101^) is different from Weber’s law (Weber^100^) in that it refers to Δ*γ* as any possible increment in *γ*, while the Weber’s law refers only to just noticeable increment in *γ*. Masin *et al*.^98^ shows that Weber’s law is a special case of Bernoulli’s principle and can be derived as follows - Equation 2 depicts the Bernoulli’s principle and increment in sensation represented by Δ*γ* is proportional to change in stimulus represented by Δ*β.*

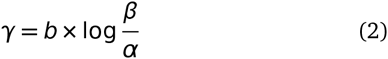

were *b* is a constant and *α* is a threshold. To evaluate the increment, the following equation 3 and the ensuing simplification gives -

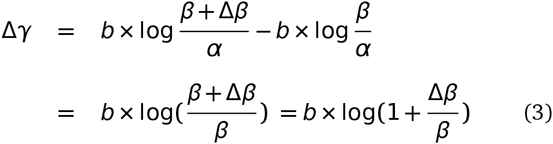

Since *b* is a constant, equation 3 reduces to 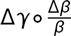 were o means "is constant when there is constancy of from Masin *et al*.^98^. The final reduction is a formulation of Weber’s laws in wordings and thus Bernoulli’s principles imply Weber’s law as a special case. Using Fechner^99^ derivation, it is possible to show the relation between Bernoulli’s principles and Weber’s law. Starting from the last line of equation 3, the following steps yield the relation -

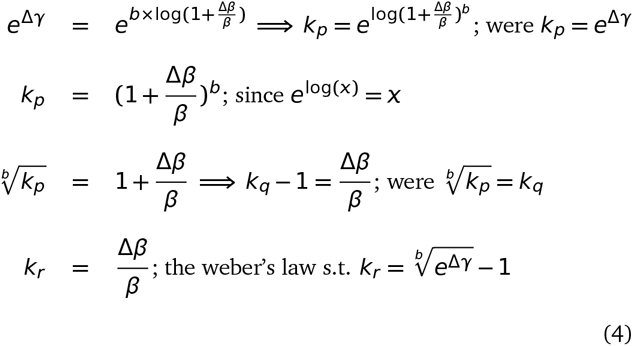

The reduction 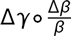 holds true given the last line of equation 4. By observation, it is important to note that the deviation Δ in the stimulus *β* plays a crucial role in the above depicted formulations. In the current study, instead of computing the sensitivity of the laws for each involved factor, the sensitivity of the deviations in the fold changes of each factor is taken into account. This is done in order to study the affect of deviations in fold changes in time as concentrations of *WNT*3*A* changes at a constant rate. Without loss of generality, it was observed over time that most involved factors had sensitivity indices or strength of contributions, parts or whole of whose graphs follow a convex or a concave curvature. These are usually represented by either an exponentially increasing or decreasing curve or nonlinear curves. This points towards the fact that with increasing changes in stimulated concentration of *WNT*3*A* the deviations in fold changes of an involved factor behave either in an increasing or decreasing fashion. Thus deviations in fold changes of various involved factors does affect the working of the signaling pathway over time. Finally, these deviations approximately capture the difference in fold changes recorded between two time frames and are thus a measure of how much the involvement of a factor affects the pathway due to these differences. This measure of involvement is depicted via the estimated sensitivity indices. The study of deviations in fold changes might help in deciding when a therapeutic drug could be administered in time. Details on the contributions of each involved factor are discussed below.

### 5.3 Analysis of deviations in fold changes

In comparison to the contributions estimated via the sensitivity indices using fold changes at different times separately, this section analyzes the contributions due to deviations in the fold changes recorded between two time points i.e 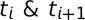. These analysis are also a way to test the efficacy of deviations in fold changes versus the absolute levels that have been stressed upon in Goentoro and Kirschner^64^. As with the analysis of the fold changes at different time points, the estimates obtained using the rbf, linear and laplacian kernels in the HSIC based sensitivity analysis have been used here. Of these, the rbf and laplacian kernels give almost similar results. Plots of the time series expression profiles from Gujral and MacBeath^17^ have been relegated to the Appendix.

■ wnt3a - Figure 33 shows the profile of mRNA expression levels of *WNT*3*A* after external stimulation. There is a series of (+−++) deviations in the fold change recordings at different time points. A repetitive behaviour is observed in the contribution of the deviations in fold changes for *WNT*3*A* estimated via the sensitivity indices. For intervals in *t*_1_, *t*_3_ and *t*_6_ there is an increase in the significance of the contribution of *WNT*3*A* in figure 10 (see the first two bars for <*t*_1_, *t*_3_ > & <*t*3, *t*6>), even though in the first three time frames levels of *WNT*3*A* are shown to be down-regulated (see figure 33). This behaviour is again repeated in figure 10 for intervals *t*_6_, *t*_12_ and *t*_24_ (see the next two bars for <*t*_6_,*t*_12_> & <*t*_12_,*t*_24_>). In both cases one finds an increase in the contribution of the deviation in the fold change. Comparing the contribution of levels of fold changes in figure 8 were it was found that there is a dip in contribution of *WNT*3*A* after *t*_3_ and then a further increase in the contribution at a latter time frame, one finds that the deviations in fold changes involving <*t*_1_,*t*_3_> & <*t*_3_,*t*_6_> have higher significance than the deviations in fold changes involving <*t*_6_,*t*_12_> & <*t*_12_,*t*_24_>. It can be noted that even in the deregulated state from <*t*_1_, *t*_3_ and *t*_6_> the deviations are minimal and the contributions are significantly high. In case of the regulated states from <*t*_6_, *t*_12_ and *t*_24_> the deviation is extremely high between the first two time frames and low in the next two. This results in greater significant contribution in the latter deviation than the former deviation. Thus when deviations are low and the fold changes over time do not vary much, the contributions of the involved factor to the signaling pathway is expected to be high and vice versa. This points to the fact that low variations in fold changes over time have a stablizing influence of *WNT*3*A* rather than abrupt high variations in fold changes that might not have the same influence. Thus measur-ments of deviations in fold changes provide greater support for studying the affect of *WNT*3*A* over time.

■ ctnnb1 - Figure 34 shows the profile of mRNA expression levels of *CTNNB*1 after external stimulation. There is a series of (++−+) deviations in the fold change recordings at different time points. An initial increase in the influence of *CTNNB*1 is observed from <*t*_1_,*t*_3_> to <*t*_3_,*t*_6_> (first two bars in figure 10) followed by a gradual decrease of influence from <*t*_3_,*t*_6_> to <*t*_6_,*t*_12_> to <*t*_12_,*t*_24_> (last three bars in figure 10). This is observed even though there is an up-regulation in levels of *CTNNB*1 with a slight dip at *t*_12_ (see figure 34). In comparison to the contribution of levels of fold changes in figure 8 were it was found that there is a gradual decrease in the influence of *CTNNB*1 till *t*_6_ and then a further increase in the contribution at latter time frames, one finds that the deviations in fold changes involving <*t*_3_,*t*_6_> have the highest significance with an almost exponential decrease in the deviations in fold changes involving <*t*_6_,*t*_12_> & <*t*_12_,*t*_24_>. Even though in a regulated state the influence of deviations in the fold changes indicate a different scenario altogether in comparison to influences of fold changes at distinct time frames. This might point to the fact that the affect of *CTNNB*1 is maximum during <*t*_3_,*t*_6_> in comparison to other stages even after constantly increasing external stimulation with *WNT*3*A* at different time points. Exponential decrease in the influence in the deviations in latter time frames points to the ineffectiveness of *CTNNB*1 in the pathway at later stages. Finally, in contrast to the behaviour of influence of *WNT*3*A* in the foregoing paragraph, *CTNNB*1 showed higher (lower) influence for greater (lesser) deviations in fold changes.

■ apc - Figure 35 shows the profile of mRNA expression levels *of APC* after external stimulation. The profile of the deviations of *APC* in a deregulated state show the following (−++−) pattern. While the *CTNNB*1 expression profile shows non-monotonic increase in levels of fold changes in upregulated state, *APC* expression profile shows a nonlinear behaviour in levels of fold changes in down regulated state. The significance of deviation in fold changes for *APC* is maximum during <*t*_3_,*t*_6_> when the down-regulation is weakened. Further weaking of the downregulation during <*t*_6_,*t*_12_> does not have much significance. This attenuation in significance of deviations in fold change might support the fact that *A PC’s* weaking in downregulation amplifies shutting down of the Wnt pathway after the intial strong downregulation (where Wnt activity is high). This is corroborated by the finding of Gujral and MacBeath^17^ which observes the initial (later) positive (negative) feedback that strenghtens (weakens) the Wnt pathway activity. An initial increase in the influence of *APC* is observed from <*t*_1_,*t*_3_> to <*t*_3_,*t*_6_> (first two bars in figure 10) followed by a gradual decrease of influence from <*t*_3_,*t*_6_> to <*t*_6_,*t*_12_> to <*t*_12_,*t*_24_> (last three bars in figure 10). This is observed even though there is an down-regulation in levels of *APC* with slight weaking at *t*_6_, *t*_12_ and *t*_24_ in comparison to recordings at other time frames (see figure 35). In comparison to the contribution of levels of fold changes in figure 8 where it was found that there is a gradual decrease in the influence of *APC* till *t*_6_ and then a further increase in the contribution at latter time frames, one finds that the deviations in fold changes involving <*t*_3_,*t*_6_> have the highest significance with an almost exponential decrease in the deviations in fold changes involving < *t*_6_,*t*_12_ > & <*t*_12_,*t*_24_>.

**Fig. 10.**
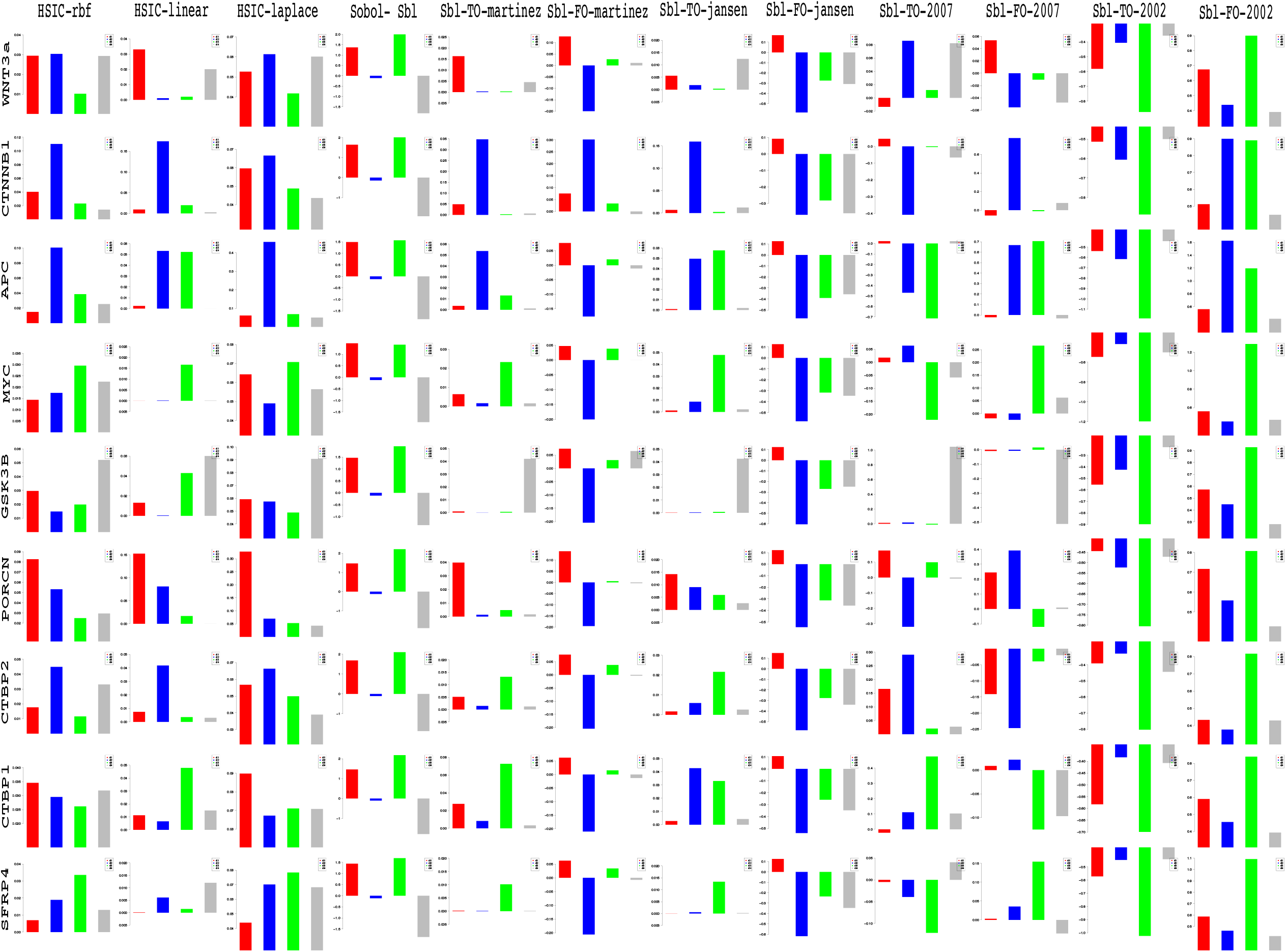
Column wise - methods to estimate sensitivity indices. Row wise - sensitivity indicies for each gene on deviations in fold change. For each graph, the bars represent sensitivity indices computed at <t1,t2> (red), <t2,t3> (blue), <t3,t4> (green) and <t4,t5> (gray). Indices were computed using non scaled time series data. TO - total order; FO -first order; SBL - Sobol

**Fig. 11.**
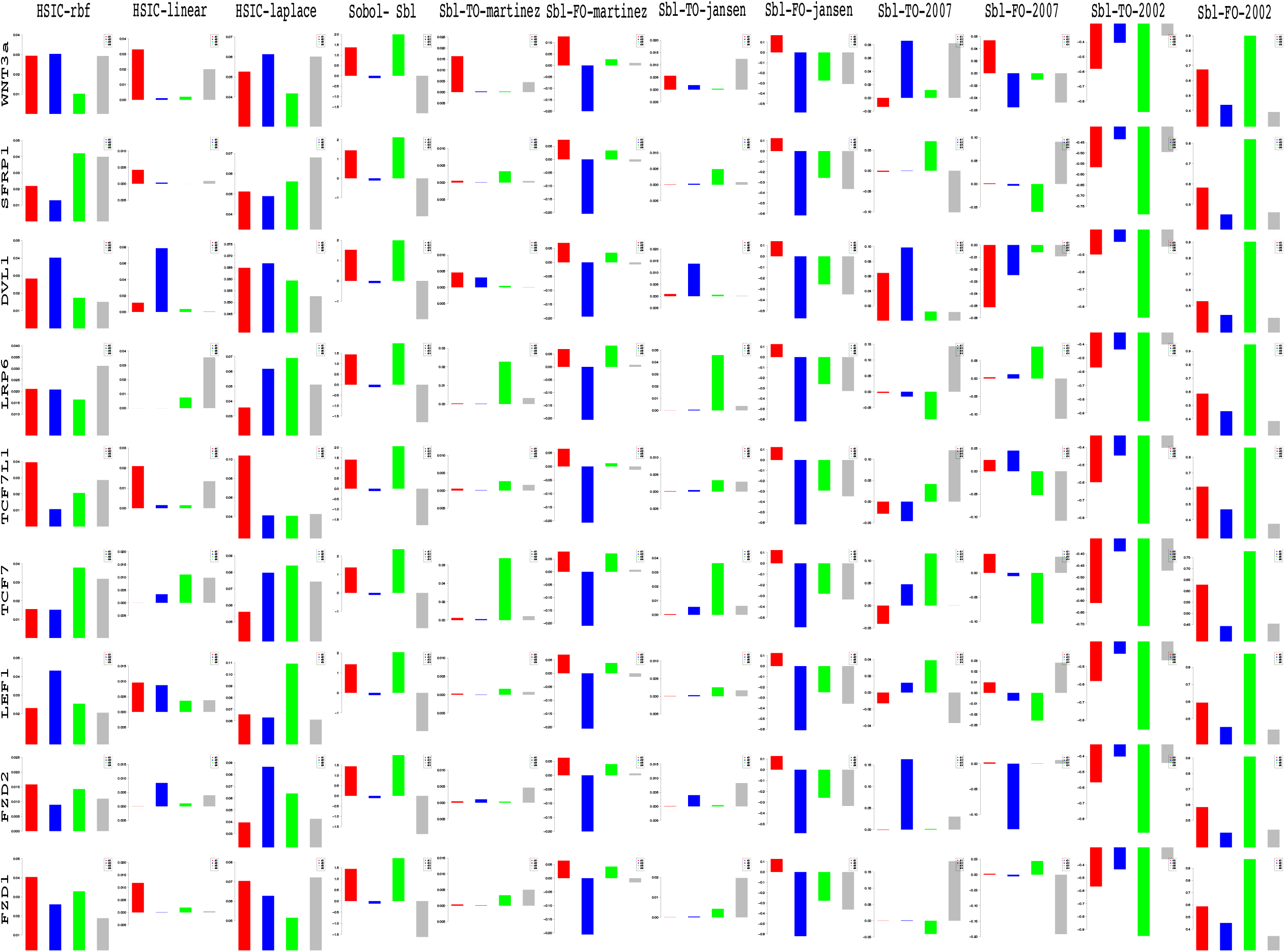
Column wise - methods to estimate sensitivity indices. Row wise - sensitivity indicies for each gene on deviations in fold change. For each graph, the bars represent sensitivity indices computed at <t1,t2> (red), <t2,t3> (blue), <t3,t4> (green) and <t4,t5> (gray). Indices were computed using non scaled time series data. TO - total order; FO -first order; SBL-Sobol

■ myc - Figure 36 shows the profile of mRNA expression levels of *MYC* after external stimulation. The profile of the deviations in fold changes of *MYC* in an up-regulated state show the following (−+++) pattern. After an initial dip in the up-regulation at *t*_6_ there is an exponential increase in the fold changes of *MYC* as time progresses. While figure 8 shows an increasing sensitivity of *MYC* for the first three time frames, later up-regulated state of *MYC* due to increasing *WNT*3*A* stimulations do not hold much significance. In contrast, it is not possible to observe a pattern in the sensitivity of deviations in fold changes for *MYC* except for the fact that the maximum contribution of deviation in fold change is observed for the period of < *t*_6_, *t*_12_ >. This is a period when *MYC’s* significance in the pathway is maximum.

■ gsk3b - Figure 37 shows the profile of mRNA expression levels of *GSK*3*β* after external stimulation. The profile of the deviations in fold changes of *GSK*3*β* in an varied regulated state show the following (−+++) pattern. After an initial up-regulation at *t*_3_ there is down regulation at *t*_6_ before which up-regulation follows for latter stages. It is widely known that *WNT* stimulation leads to inhibition of *GSK*3*β*. In contrast to this regard *GSK*3*β* shows a up-regulated levels at *t*_3_, *t*_12_ and *t*_24_. The author is currently unaware of why this contasting behaviour is exhibited. Later upregulation might point to the fact that the effectiveness of Wnt stimulation has decreased and *GSK*3*β* plays the role of stabilizing and controlling the behaviour of the pathway by working against the Wnt stimulation and preventing further degradation. While work by Gujral and MacBeath^17^ does not shed light on this aspect, contrasting models of inhibitions for *GSK*3 has been recently proposed in Metcalfe and Bienz^102^ which might support this behaviour. Figure 8 shows an decreasing sensitivity of *GSK*3*β* for the first two time frames, after which there is an increasing sensitivity for the next three time frames. Comparing this with plots in figure 10 it is found that there is greater significance of deviations in fold changes of *GSK*3*β* during later stages of <*t*_6_,*t*_12_> and <*t*_12_,*t*_24_>.

■ porcn - Figure 38 shows the profile of mRNA expression levels of *PORCN* after external stimulation. The profile of the deviations in fold changes of *PORCN* in an up regulated state show the following (+−−−) pattern. After an initial hike in up-regulation at *t*_3_ there is continuous decrease in the up regulation. *PORCN* is known to help in the secretion of the Wnt ligands that later on help in the instigation of the signaling activity. Sustanined stimulation by *WNT*3*A* over a period of time might lead to decrease in the up regulation of *PORCN* which helps in Wnt secretion. Graph for *PORCN* in figure 8 shows increasing significance of the influence of *PORCN* as time passes, even though there is lower regulation of the same at later stages (fig 38). The highly significant influence of lower regulation at later stages indicates the lessened effectiveness of *PORCN* due to sustained *WNT*3*A* stimulation that might have suppressed the functionality of secretion carried out via *PORCN*. Contrary to this, the influences of the deviations in the fold changes over time show the reverse behaviour. The maximum influence is during the first two time frames of <*t*_1_,*t*_3_> and this influence of deviations decreases at later stages. This points to the fact that the deviations in the fold changes at intial stage has greater significance in the pathway than the deviations at later stages. It follows that in initial stages of Wnt stimulation the expression of *PORCN* has significant influence.

■ ctbp2 - Figure 39 shows the profile of mRNA expression levels of *CTBP*2 after external stimulation. The profile of the deviations in fold changes of *CTBP*2 in an up regulated state show the following (−++−) pattern. It is known that *CTBP*2 shows co-repressive nature and the pattern of sensitivity indicates this heigthened effect at <*t*_3_,*t*_6_> and <*t*_12_, *t*_24_>. In contrast to this in figure 8 one finds that the significance of upregulation at *t*_12_ and *t*_24_ is minimal and yet the sensitivity for the deviations in fold changes for this period is second to <*t*_3_, *t*_6_>. The probable explanation for this might be that for higher upregulation (in terms of numeric representations), even small deviations might play a sigificant role while the sensitivity at individual time frames remains low.

■ ctbp1 - Figure 40 shows the profile of mRNA expression levels of *CTBP*1 after external stimulation. The profile of the deviations in fold changes of *CTBP*1 in an up regulated state show the following (+++−) pattern. As with the heightened sensitivity at time frame *t*_1_ the sensitivity of deviations in the fold changes exhibits heightened effect in the pathway at <*t*_1_, *t*_3_>. Further analysis might not be possible as one finds lowered sensitivity even at heightened up regulation for individual recordings as well as deviations.

■ sfrp4 - Figure 41 shows the profile of mRNA expression levels of *SFRP*4 after external stimulation. The profile of the deviations in fold changes of *SFRP*4 in an down regulated (except in the last time frame) state show the following (−+++) pattern. Known to be a negative regulator of the Wnt pathway, it was found that its sensitivity in extremely high as it is down regulated during the stimulation. This is depicted in the figures plotted in 8. This heightened sensitivity during most of the down regulation points towards the significant role of hypermethylation that leads to silencing of this gene. Contrary to this, there is a monotonically increasing sensitivity for deviations in the fold changes during down regulation from *t*_1_ to *t*_12_. A dip in the sensitivity of the deviation for the final time frame < *t*_12_, *t*_24_ > happens when the up regulation is recorded in the last time frame. Based on the maximum sensitivity for deviation in fold change during <*t*_6_, *t*_12_>, up regulation of *SFRP*4 in this period is expected to have greatest reverse effect on the activation of the Wnt pathway. It appears that the hypermethylation that causes the silencing of the *SFRP*4 is maximal during this stage and thus a potential period for the pathway to be inhibited via reversal of silencing (Suzuki *et al.*^89^).

■ sfrp1 - Figure 42 shows the profile of mRNA expression levels of *SFRP*1 after external stimulation. The profile of the deviations in fold changes of *SFR P*4 in an up regulated (except in during the second time frame) state show the following (−+−−) pattern. It is widely known that *SFRP*1 is a Wnt antagonist and is known for inactivation in the canconical Wnt pathway due to hypermethylation thus leading to upregulation of the pathway (Caldwell *et al*.^90^). Suzuki *et al*.^89^ further indicates that *SFRP*1 is thought to silence ligand-dependent Wnt signaling by binding of the cysteine rich-domain (CRD) to Wnt proteins, thus preventing interaction with *FRZ* receptors. Recent in silico results by Kim and Kim^103^ confirm hypermethylation of *SFRP*1 in colorectal cancers. Given the above profile, it is possible to see that there is a down regulation at *t*3 but the significance of its influence on the pathway is not much as revealed in figure 9. Figure 9 shows a decreasing significance in the influence of the *SFRP*1 with the maximum influence in the last stage of the *WNT*3*A* stimulation, where there is an up regulation. In similar way, in figure 11 there is significance of the sensitivity in the deviations in fold changes during the up regulation of *SFRP*1 during the <*t*_6_, *t*_12_> and <*t*_12_,*t*_24_>. The activation at later stages show that *SFRP*1 has a greater antagonistic effect on the Wnt pathway. In comparison to up regulation, one finds the down regulation at *t*_3_ does not play a significant role. These are in line with Gujral and MacBeath^17^’s claim regarding reversal of behaviour at different time stages.

■ dvl1 - Figure 43 shows the profile of mRNA expression levels of *DVL*1 after external stimulation. The profile of the deviations in fold changes of *DVL*1 in an up regulated state show the following (−++−) pattern. *DVL*1 is an adaptor protein that helps in signal transmission that leads to stabilization of cytosolic *β-catenln* for further processing. Huang *et al.*^87^ report high expression of *DVL*1 in Taiwanese colorectal cancer patients with liver metastasis and it has also been observed as a potential biomarker in CRC (Wu *et al.*^104^). In figure 9 the time frame *t*_12_ at which DVL1 shows maximum up regulation is the most insignificant one due to the lowest sensitivity while moderate upregulation during *t*_3_ and *t*_6_ show high sensitivity. The same is true for upregulation at *t*_24_. Comparing this with the deviations in the fold changes in 11, one finds that there is maximum sensitivity during the period of <*t*_3_, *t*_6_> preceeded by a lower sensitivity index for the perioud of < *t*_1_, *t*_3_ >. At other intervals there was a decreased sensitivity even though the deviations in the fold changes were very high. This indicates that the high deviations might not influence the signaling activity significantly. Also the best period of intervention is at <*t*_3_, *t*_6_>.

■ lrp6 - Figure 44 shows the profile of mRNA expression levels of *Lrp*6 after external stimulation. The profile of the deviations in fold changes of *Lrp6* in an up regulated state (except at *t*_3_) show the following (−++−) pattern. In an extensive work on the molecular differences of *LRPs* MacDonald *et al.*^105^ investigate and show that *LRP*5/6 along with the frizzled family members form a Wnt inducible co-receptor complex that helps in signal transmission after *LRP* phosphorylation. Earlier wet lab work by Liu *et al.*^106^ and in silico findings by Watanabe *et al.*^107^ have shown highly expressed participation of *LRP*5/6 in Wnt signaling pathway. Latest work by Lemieux *et al.*^108^ shows that KRAS signaling promotes canonical Wnt activity via *LRP*6. In figure 9, *LRP*6 shows significant influences during the *t*_1_, *t*_6_ and *t*_12_. The only period in which it is down regulated during *t*_3_ has little significance in comparison to the significance of the up regulated states. It is not known why *LRP*6 shows down regulation at this stage. Finally, for unkown reasons, the influence of *LRP*6 during *t*_12_ was found to be the lowest. It was not possible to read into the sensitivity of *LRP*6 for deviations in fold changes the HSIC based indices. The laplace kernel shows a pattern of increasing sensitivity with the heightest during < *t*_6_, *t*_12_ >. But this is not so in the other two formulations. Thus wet lab experiments might aid in confirming these results and shedding more light on the duration during which a drug could be administered.

■ tcf7l16 - Figure 45 shows the profile of mRNA expression levels of *TCF*7*L*1 (also known as *TCF*3) after external stimulation. The profile of the deviations in fold changes of *TCF*7*L*1 in a down regulated state (except at *t*_3_) show the following (+−+−) pattern. It is known that Wnt stimulation promotes the phosphorylation of repressor-acting *TCF*7*L*1 by homeodomain-interacting protein kinase (HIPK2), which results in its dissociation from the WRE (Hikasa and Sokol^109^). Gujral and MacBeath^17^’s results also indicate the same repression of *TCF*7*L*1 during *WNT*3*A* simulation as shown in figure 45. But in contradiction to this recent findings of Leushacke *et al*.^110^, *TCF*7*L*1 is found to be expressed in the colon crypt and in colon cancer. Their results indicate that *TCF*7*L*1 may have an as yet unidentified role in transmission of tumor-related *β-catenin* signals. Evidence of this up regulation is found at *t*_3_ time period as shown in figure 45. From figure 9 it can be seen that the sensivi-tiy of *TCF*7*L*1 is maximum during the first time period. Later on the sensitivity subsides as time passes untill the sensitivity shots up in the last time frame *t*_24_. In comparison to this with respect to the deviations in fold changes over time in figure 11, <*t*_1_,*t*_3_> showed the maximum period of influence. Later on there is a drop in the sensitivity which is followed by an approximately monotonic increase. It is the first transition from down regulation to up regulation <*t*_1_, *t*_3_> that might be the time for intervention. Also the last stage might be of some value but during down regulation only.

■ tcf7 - Figure 46 shows the profile of mRNA expression levels of *TCF*7 after external stimulation. The profile of the deviations in fold changes of *TCF*7 in an up regulated state show the following (−−++) pattern. *TCF*7 is found to be regulated upon Wnt stimulation as it binds with *LEFs* to activate the transcription procedure after interacting with *β-catenin* Cadigan and Waterman^111^. In figure 9, the sensitivity of the activation of *TCF*7 decreases monotonically as time progresses. But this behaviour is not the same for deviations in the fold changes. The maximum influence is found for the duration of <*t*_6_, *t*_12_>. The next best consistent influence is in the duration <*t*_12_, *t*_24_>. These are the two time periods when the influence of the deviations in the fold changes is maximum and thus susceptible to therapeutic interference.

■ lef1 - Figure 47 shows the profile of mRNA expression levels of *LEF*1 after external stimulation. The profile of the deviations in fold changes of *LEF*1 in an up regulated state (except at *t*_31_ show the following (−−++) pattern. Generally, *LEF*1 is found to be up regulated upon Wnt stimulation when it works in tandem with *TCF*7 Cadigan and Waterman^111^. Yet, in figure 9, the sensitivity of the activation of *LEF*1 is not similar to that of *TCF*7. In contradiction to what is observed, one finds a down regulation during the time period *t*_3_. More importantly this is the period in which the most significant influence of down regulated *LEF*1 is observed. The Initial down regulation at this subinterval indicates that *LEF*1 is not facilitating the Wnt pathway positiviely Conclusive results cannot be stated regarding the deviations in fold changes from figure 11.

■ fzd2 - Figure 48 shows the profile of mRNA expression levels of *FZD*2 after external stimulation. The profile of the deviations in fold changes of *FZD*2 in an up regulated state (except at *t*_3_ and *t*_6_) show the following (−−++) pattern. The *FZD* or the frizzled family of 7-transmember protein (Ueno *et al.*^74^) works in tandem with *LRP*-5/6 as binding parameters for the Wnt ligands to initiate the Wnt signaling. In comparison to the repetitive behavior shown in figure 9 it is not possible to draw conclusions on the deviations in fold changes.

■ fzd1 - Figure 49 shows the profile of mRNA expression levels of *FZD*1 after external stimulation. The profile of the deviations in fold changes of *FZD*1 in an down regulated state (except at *t*_3_) show the following (+−−+) pattern. Consistent with the findings of Holcombe *et al*.^75^ and Planutis *et al.*^76^, *FZD*1 was found to be expressed at *t*_3_. In the rest of the time periods, it was down regulated. But the significance of the influence shows a different pattern in figure 9 with the down regulation at *t*_12_ being the most influencial. In contrast to this, while observing the deviations in the fold changes, it was found that the first two durations <*t*_1_,*t*_3_> and <*t*_3_, *t*_6_> showed consistent decreasing behaviour in terms of influence. It is during the first period that the deviations in fold changes are significant and thus it is possible to intervene therapeutically during the activation stage.

Indicies for remaining 57 genes as well as analysis of the same will be presented in the following B part of this manuscript. Graphs for these 57 genes have been presented in figures 31 and 32 in the Appendix.

## 6 Conclusion

### COMPUTATIONAL SIGNIFICANCE

Local and global sensitivity analysis on static and time series measurements in Wnt signaling pathway for colorectal cancer is done. Density based Hilbert-Schmidt Information Criterion indices outperformed the variance based Sobol indices. This is attributed to the employment of distance measures & the kernel trick via Reproducing kernel Hilbert space (RKHS) that captures nonlinear relations among various intra/extracellular factors of the pathway in a higher dimensional space. The gained advantage is confirmed on the inferred results obtained via a Bayesian network model based on prior biological knowledge and static gene expression data. In time series data, using these indices it is now possible to observe when and in which period of time and to what degree a factor gets influenced & contributes to the pathway, as changes in concentration of the another factor is made. This facilitates in time based administration of target therapeutic drugs & reveal hidden biological information within colorectal cancer samples.

### DEVIATIONS IN FORMULATION OF PSYCHOPHYSICAL LAW

In context of Goentoro and Kirschner^64^’s work regarding the recent development of observation of Weber’s law working downstream of the pathway, it has been found that the law is governed by the ratio of the deviation in the input and the absolute input value. More importantly, it is these deviations in input that are of significance in studing any phemomena. The current manuscript explores the sensitivity of deviation in the fold changes between measurements of fold changes at consecutive time points to explore in what duration of time i.e < *t*_*i*_, *t*_*i*+1_ >, a particular factor is affecting the pathway in a major way. This has deeper implications in the fact that one is now able to observe when in time an intervention can be made or a gene be perturbed to study the behaviour of the pathway in tumorous cases. Thus sensitivity analysis of deviations in mathematical formulations of the psychophysical law can lead to insights into the time period based influence of the involved factors in the pathway. This will also shed light on the duration in which the psychophysical laws might be most prevalent.

### SPECIFIC EXAMPLES OF BIOLOGICAL INTERPRETATIONS

gsk3*β* - It is widely known that *WNT* stimulation leads to inhibition of *GSK*3*β*. In contrast to this regard *GSK*3*β* shows a up-regulated levels at *t*_3_, *t*_12_ and *t*_24_. The author is currently unaware of why this contasting behaviour is exhibited. Later up-regulation might point to the fact that the effectiveness of Wnt stimulation has decreased and *GSK*3*β* plays the role of stabilizing and controlling the behaviour of the pathway by working against the Wnt stimulation and preventing further degradation. While work by Gujral and MacBeath^17^ does not shed light on this aspect, contrasting models of inhibitions for *GSK*3 has been recently proposed in Metcalfe and Bienz^102^ which might support this behaviour. Considering analysis of fold changes at different time points, decreasing sensitivity of *GSK*3*β* was observed for the first two time frames, after which there is an increasing sensitivity for the next three time frames. Comparing this with plots of analysis of deviations in fold changes, it is observed that there is greater significance of deviations in fold changes of *GSK*3*β* during later stages of < *t*_6_, *t*_12_ > and < *t*_12_, *t*_24_ >. It is in these periods that one might be able to pertube and study significant affects on the pathway.

porcn - *PORCN* is known to help in the secretion of the Wnt ligands that later on help in the instigation of the signaling activity. Sustanined stimulation by *WNT*3*A* over a period of time might lead to decrease in the up regulation of *PORCN* which helps in Wnt secretion. Graph for *PORCN* in analysis of fold changes shows increasing significance of the influence of *PORCN* as time passes, even though there is lower regulation of the same at later stages. The highly significant influence of lower regulation at later stages indicates the lessened effectiveness of *PORCN* due to sustained *WNT*3*A* stimulation that might have suppressed the functionality of secretion carried out via *PORCN*. Contrary to this, the influences of the deviations in the fold changes over time show the reverse behaviour. The maximum influence is during the first two time frames of < *t*_1_,*t*_3_ > and this influence of deviations decreases at later stages. This points to the fact that the deviations in the fold changes at intial stage has greater significance in the pathway than the deviations at later stages. It follows that in initial stages of Wnt stimulation the expression of *PORCN* has significant influence.

## 7 Acknowledgement

Sincere thanks to anonymous reviwers who have provided input to refine this manuscript. Part of this work has been accepted for poster presentation at the International conference for Systems Biology of Human Disease 2016. The author thanks Harvard Program in Therapeutics Sciences for granting registration fee scholarship for this work after evaluation of the poster abstract. Also the author thanks Mrs. Rita Sinha and Mr. Prabhat Sinha for supporting him financially on this project.

## Appendix

### Sensitivity indices

#### Variance based indices

The variance based indices as proposed by Sobol’^21^ prove a theorem that an integrable function can be decomposed into summands of different dimensions. Also, a Monte Carlo algorithm is used to estimate the sensitivity of a function apropos arbitrary group of variables. It is assumed that a model denoted by function 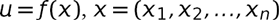, is defined in a unit *n*-dimensional cube 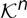 with *u* as the scalar output. The requirement of the problem is to find the sensitivity of function *ƒ*(*x*) with respect to different variables. If *u*\* =*ƒ*(*x*\*) is the required solution, then the sensitivity of *u*\* apropos *x*_*k*_ is estimated via the partial derivative 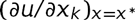. This approach is the local sensitivity. In global sensitivity, the input *x* = *x*\* is not specified. This implies that the model *ƒ*(*x*) lies inside the cube and the sensitivity indices are regarded as tools for studying the model instead of the solution. Detailed technical aspects with examples can be found in Homma and Saltelli^33^ and Sobol^65^.

Let a group of indices 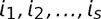 exist, where 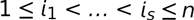 and 1 ≤ *s* ≤ *n*. Then the notation for sum over all different groups of indices is -

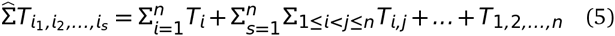

Then the representation of *ƒ*(*x*) using equation 5 in the form -

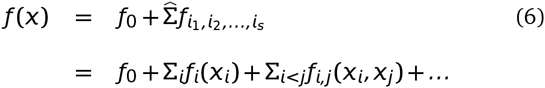

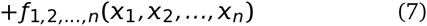

is called ANOVA-decomposition from Archer *et al.*^46^ or expansion into summands of different dimensions, if *ƒ*_0_ is a constant and integrals of the summands 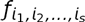 with respect to their own variables are zero, i.e,

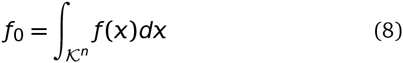

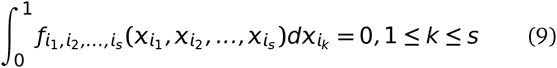

It follows from equation 7 that all summands on the right hand side are orthogonal, i.e if at least one of the indices in 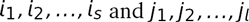 is not repeated i.e

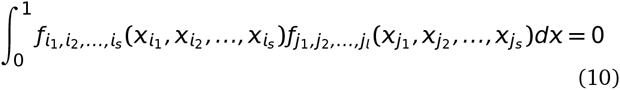

Sobol’^21^ proves a theorem stating that there is an existence of a unique expansion of equation 7 for any*ƒ*(x) integrable in 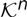. In brief, this implies that for each of the indices as well as a group of indices, integrating equation 7 yields the following -

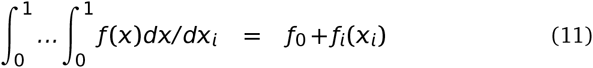

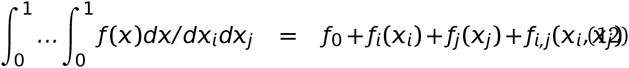

were, *dx*/*dx*_*i*_ is 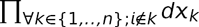 and 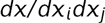 is 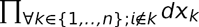. For higher orders of grouped indices, similar computations follow. The computation of any sum-mand 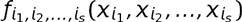 is reduced to an integral in the cube 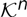. The last summand 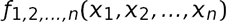 is *ƒ*(*x*) – *ƒ*_0_ from equation 7. Homma and Saltelli^33^ stresses that use of Sobol sensitivity indices does not require evaluation of any 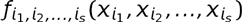 nor the knowledge of the form of *ƒ*(*x*) which might well be represented by a computational model i.e a function whose value is only obtained as the output of a computer program.

Finally, assuming that *ƒ*(*x*) is square integrable, i.e 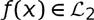, then all of 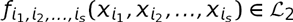. Then the following constants

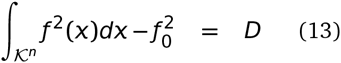

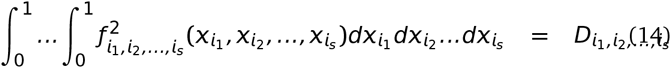

are termed as variances. Squaring equation 7, integrating over 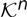 and using the orthogonality property in equation 10, *D* evaluates to -

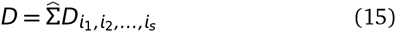

Then the global sensitivity estimates is defined as -

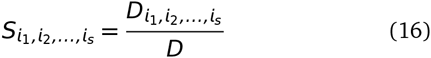

It follows from equations 15 and 16 that

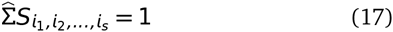

Clearly, all sensitivity indices are non-negative, i.e an index 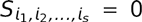 if and only if 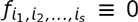. The true potential of Sobol indices is observed when variables 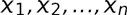 are divided into *m* different groups with *y*_1_, *y*_2_, …, *y*_*m*_ such that *m* < *n*. Then*ƒ*(*x*)☰*ƒ*(*y*_1_, *y*_2_, …, *y*_*m*_). All properties remain the same for the computation of sensitivity indices with the fact that integration with respect to *y*_*k*_ means integration with respect to all the *x_i_’s* in *y*_*k*_. Details of these computations with examples can be found in Sobol^65^. Variations and improvements over Sobol indices have already been stated in section 2.1.

#### Density based indices

As discussed before, the issue with variance based methods is the high computational cost incurred due to the number of interactions among the variables. This further requires the use of screening methods to filter out redundant or unwanted factors that might not have significant impact on the output. Recent work by Da Veiga^56^ proposes a new class of sensitivity indicies which are a special case of density based indicies Borgonovo^53^. These indicies can handle multivariate variables easily and relies on density ratio estimation. Key points from Da Veiga^56^ are mentioned below.

Considering the similar notation in previous section, 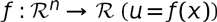 is assumed to be continuous. It is also assumed that *X*_*k*_ has a known distribution and are independent. Baucells and Borgonovo^66^ state that a function which measures the similarity between the distribution of *U* and that of *U|X*_*k*_ can define the impact of *X*_*k*_ on *U.* Thus the impact is defined as -

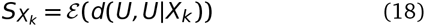

were *d*(·,·) is a dissimilarity measure between two random variables. Here *d* can take various forms as long as it satisfies the criteria of a dissimilarity measure. Csiszár *et al.*^59^’s f-divergence between *U* and *U|X*_*k*_ when all input random variables are considered to be absolutely continuous with respect to Lebesgue measure on 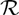 is formulated as -

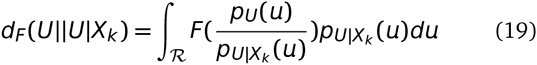

were *F* is a convex function such that *F*(1) = 0 and *p*_*U*_ and 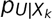 are the probability distribution functions of *U* and *U|X_k_.* Standard choices of *F* include Kullback-Leibler divergence *F*(*t*) = −log_*e*_(*t*), Hellinger distance 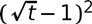, Total variation distance *F*(*t*) = |*t*−1|, Pearson *χ*^2^ divergence *F*(*t*) = *t*^2^ − 1 and Neyman *χ*^2^ divergence *F*(*t*) = (1 − *t*^2^)/t. Substituting equation 19 in equation 18, gives the following sensitivity index -

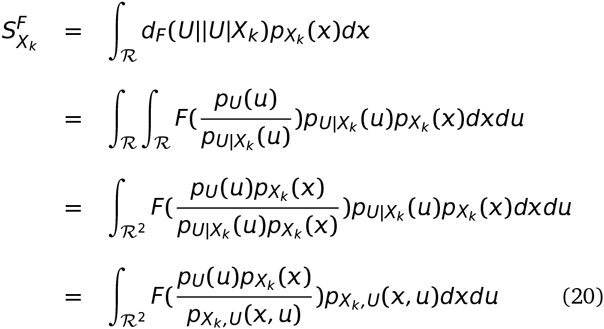

were 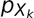 and 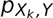 are the probability distribution functions of *X*_*k*_ and (*X*_*k*_, *U*), respectively. Csiszár *et al.*^59^ f-divergences imply that these indices are positive and equate to 0 when *U* and *X*_*k*_ are independent. Also, given the formulation of 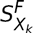, it is invariant under any smooth and uniquely invertible transformation of the variables *X*_*k*_ and *U* (Kraskov *et al.^67^).* This has an advantage over Sobol sensitivity indices which are invariant under linear transformations.

By substituting the different formulations of *F* in equation 20, Da Veiga^56^’s work claims to be the first in establishing the link that previously proposed sensitivity indices are actually special cases of more general indices defined through Csiszár *et al.*^59^’s f-divergence. Then equation 20 changes to estimation of ratio between the joint density of (*X_k_, U*) and the marginals, i.e -

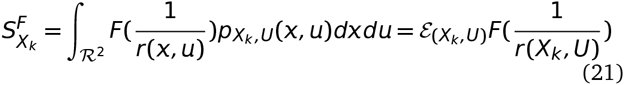

were, 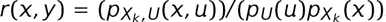. Multivariate extensions of the same are also possible under the same formulation.

Finally, given two random vectors 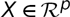 and 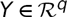, the dependence measure quantifies the dependence between *X* and *Y* with the property that the measure equates to 0 if and only if *X* and *Y* are independent. These measures carry deep links (Sejdinovic *et al.*^68^) with distances between embeddings of distributions to reproducing kernel Hilbert spaces (RHKS) and here the related Hilbert-Schmidt independence criterion (HSIC by Gretton *et al.*^58^) is explained.

In a very brief manner from an extremely simple introduction by Daumé III^69^ - “We first defined a field, which is a space that supports the usual operations of addition, subtraction, multiplication and division. We imposed an ordering on the field and described what it means for a field to be complete. We then defined vector spaces over fields, which are spaces that interact in a friendly way with their associated fields. We defined complete vector spaces and extended them to Banach spaces by adding a norm. Banach spaces were then extended to Hilbert spaces with the addition of a dot product.” Mathematically, a Hilbert space 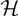 with elements 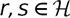 has dot product 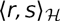 and *r · s.* When 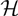 is a vector space over a field F, then the dot product is an element in 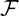. The product 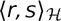 follows the below mentioned properties when 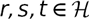 and for all 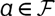 -

- Associative: (*ar*)·*s* = *a*(*r·s*)
- Commutative: *r·s* = *s·r*
- Distributive: r·(*s* + *t*) = *r*·*s* + *r·t*

Given a complete vector space 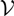 with a dot product 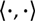, the norm on 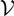 defined by 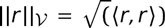 makes this space into a Banach space and therefore into a full Hilbert space.

A reproducing kernel Hilbert space (RKHS) builds on a Hilbert space 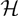 and requires all Dirac evaluation functionals in 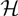 are bounded and continuous (on implies the other). Assuming 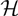 is the 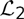 space of functions from 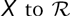 for some measurable *X.* For an element 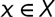, a Dirac evaluation functional at x is a functional 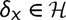 such that 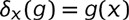. For the case of real numbers, *x* is a vector and *g* a function which maps from this vector space to 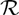. Then 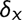 is simply a function which maps *g* to the value *g* has at *x*. Thus, *δ*_*x*_ is a function from 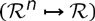 into 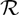.

The requirement of Dirac evaluation functions basically means (via the Riesz^70^ representation theorem) if *ϕ* is a bounded linear functional (conditions satisfied by the Dirac evaluation functionals) on a Hilbert space *H*, then there is a unique vector 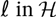 such that 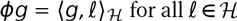 for all 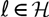. Translating this theorem back into Dirac evaluation functionals, for each *δ*_*x*_ there is a unique vector *k*_*x*_ in 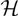 such that 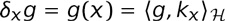. The reproducing kernel *K* for 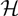 is then defined as: 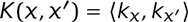, were *k*_*x*_ and *k*_*x′*_ are unique representatives of *δ*_*x*_ and *δ_x′_.* The main property of interest is 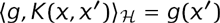. Furthermore, *k*_*x*_ is defined to be a function 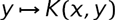 and thus the reproducibility is given by 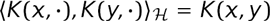.

Basically, the distance measures between two vectors represent the degree of closeness among them. This degree of closeness is computed on the basis of the discriminative patterns inherent in the vectors. Since these patterns are used implicitly in the distance metric, a question that arises is, how to use these distance metric for decoding purposes?

The kernel formulation as proposed by Aizerman *et al.*^60^, is a solution to our problem mentioned above. For simplicity, we consider the labels of examples as binary in nature. Let 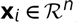, be the set of *n* feature values with corresponding category of the example label (*y*_*i*_) in data set 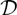. Then the data points can be mapped to a higher dimensional space *H* by the transformation *ϕ:*

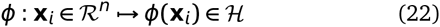

This 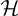 is the *Hilbert Space* which is a strict inner product space, along with the property of completeness as well as separability. The inner product formulation of a space helps in discriminating the location of a data point w.r.t a separating hyperplane in 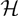. This is achieved by the evaluation of the inner product between the normal vector representing the hyperplane along with the vectorial representation of a data point in 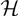 (Figure 12 represents the geometrical interpretation). Thus, the idea behind equation (22) is that even if the data points are nonlinearly clustered in space *R*^*n*^,the transformation spreads the data points into 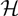, such that they can be linearly separated in its range in 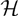.

**Fig. 12.**
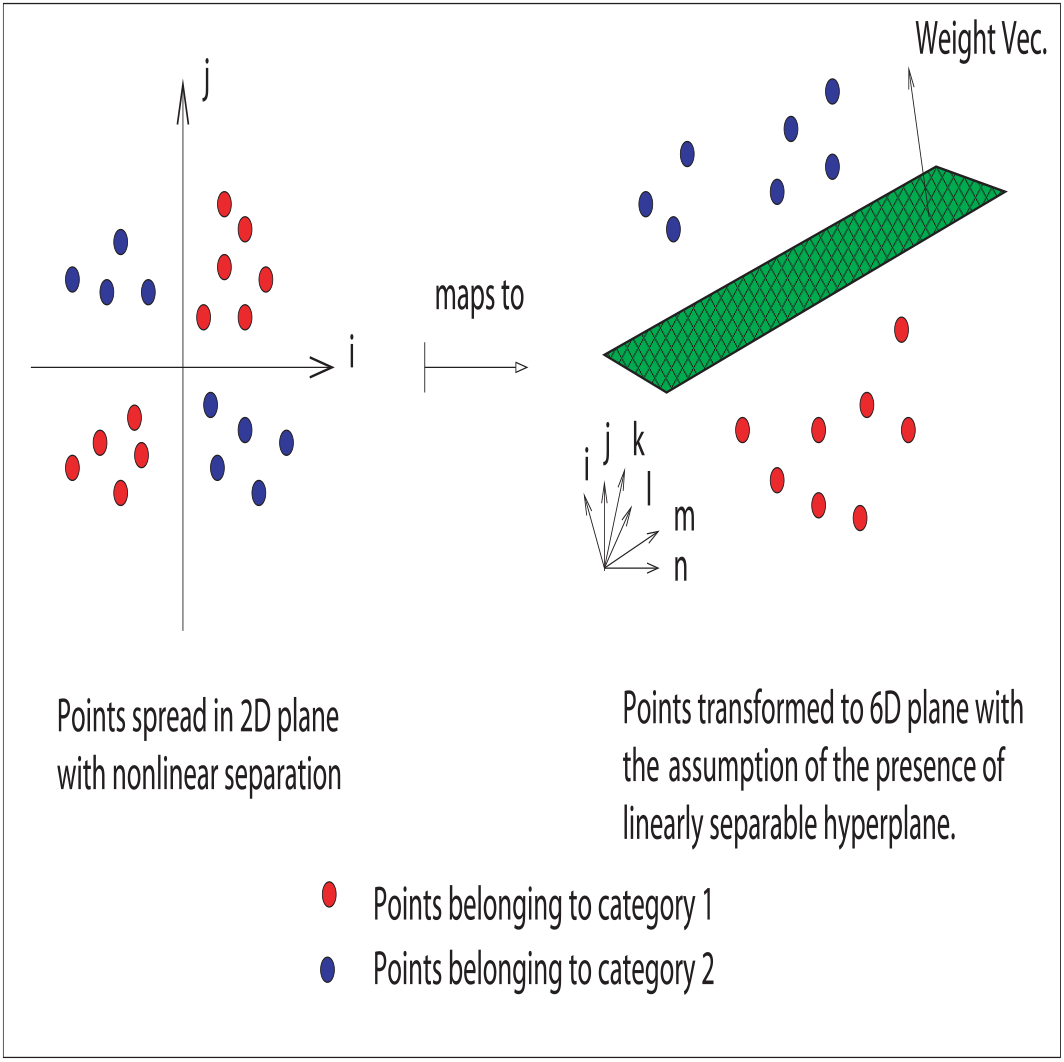
A geometrical interpretation of mapping nonlinearly separable data into higher dimensional space where it is assumed to be linearly separable, subject to the holding of dot product.

Often, the evaluation of dot product in higher dimensional spaces is computationally expensive. To avoid incurring this cost, the concept of kernels in employed. The trick is to formulate kernel functions that depend on a pair of data points in the space 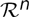, under the assumption that its evaluation is equivalent to a dot product in the higher dimensional space. This is given as:

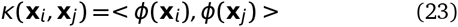

Two advantages become immediately apparent. First, the evaluation of such kernel functions in lower dimensional space is computationally less expensive than evaluating the dot product in higher dimensional space. Secondly, it relieves the burden of searching an appropriate transformation that may map the data points in 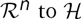. Instead, all computations regarding discrimination of location of data points in higher dimensional space involves evaluation of the kernel functions in lower dimension. The matrix containing these kernel evaluations is referred to as the *kernel* matrix. With a cell in the kernel matrix containing a kernel evaluation between a pair of data points, the kernel matrix is square in nature.

As an example in practical applications, once the kernel has been computed, a pattern analysis algorithm uses the kernel function to evaluate and predict the nature of the new example using the general formula:

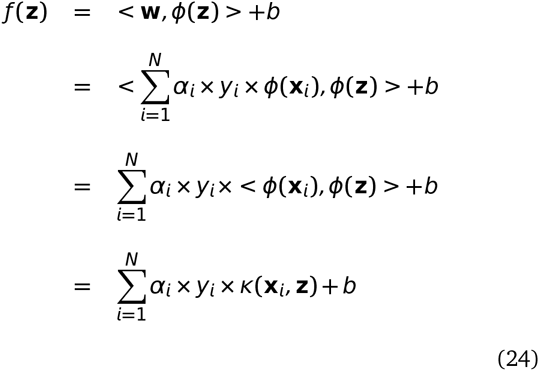

where **w** defines the hyperplane as some linear combination of training basis vectors, **z** is the test data point, *y*_*i*_ the class label for training point **x**_*i*_, *α*_*i*_ and *b* are the constants. Various transformations to the kernel function can be employed, based on the properties a kernel must satisfy. Interested readers are referred to Taylor and Cristianini^71^ for description of these properties in detail.

The Hilbert-Schmidt independence criterion (HSIC) proposed by Gretton *et al.*^58^ is based on kernel approach for finding dependences and on cross-covariance operators in RKHS. Let 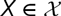 have a distribution *P*_*X*_ and consider a RKHS 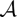 of functions 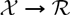 with kernel 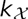 and dot product 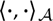. Similarly, Let 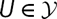 have a distribution 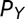 and consider a RKHS 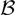 of functions 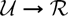 with kernel 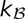 and dot product 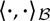. Then the cross-covariance operator 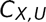 associated with the joint distribution 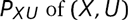 is the linear operator 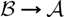 defined for every 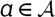 and 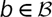 as -

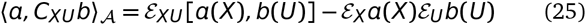

The cross-covariance operator generalizes the covariance matrix by representing higher order correlations between *X* and *U* through nonlinear kernels. For every linear operator 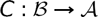 and provided the sum converges, the Hilbert-Schmidt norm of *C* is given by -

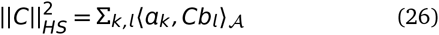

were *a*_*k*_ and *b*_*l*_ are orthonormal bases of 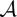 and 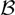, respectively. The HSIC criterion is then defined as the Hilbert-Schmidt norm of cross-covariance operator -

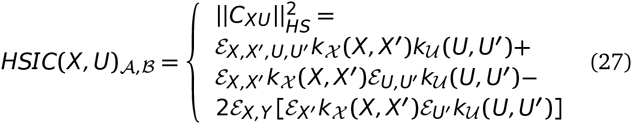

were the equality in terms of kernels is proved in Gretton *et al.*^58^. Finally, assuming 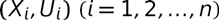 is a sample of the random vector (*X, U*) and 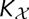 and 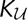 denote the Gram matrices with entries 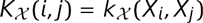 and 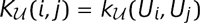. Gretton *et al.*^58^ proposes the following estimator for 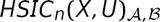 -

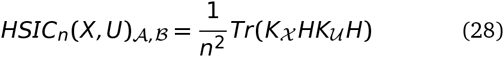

were *H* is the centering matrix such that 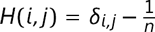. Then 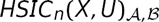 can be expressed as -

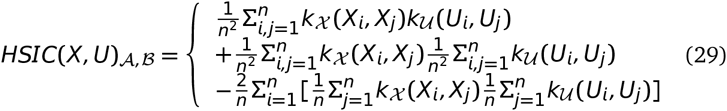

Finally Da Veiga^56^ proposes the sensitivity index based on distance correlation as -

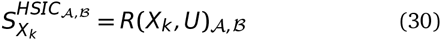

were the kernel based distance correlation is given by -

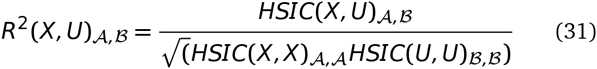

were kernels inducing *A* and *B* are to be chosen within a universal class of kernels. Similar multivariate formulation for equation 28 are possible.

## Choice of sensitivity indices

The sensitivity package (Faivre *et al.*^72^ and Iooss and Lemaître^22^) in R langauge provides a range of functions to compute the indices and the following indices will be taken into account for addressing the posed questions in this manuscript.

1. sensiFdiv - conducts a density-based sensitivity analysis where the impact of an input variable is defined in terms of dissimilarity between the original output density function and the output density function when the input variable is fixed. The dissimilarity between density functions is measured with Csiszar f-divergences. Estimation is performed through kernel density estimation and the function kde of the package ks. (Borgonovo^53^, Da Veiga^56^)
2. sensiHSIC - conducts a sensitivity analysis where the impact of an input variable is defined in terms of the distance between the input/output joint probability distribution and the product of their marginals when they are embedded in a Reproducing Kernel Hilbert Space (RKHS). This distance corresponds to HSIC proposed by Gretton *et al.*^58^ and serves as a dependence measure between random variables.
3. soboljansen - implements the Monte Carlo estimation of the Sobol indices for both first-order and total indices at the same time (all together 2p indices), at a total cost of (p+2) × n model evaluations. These are called the Jansen estimators. (Jansen^48^ and Saltelli *et al.*^41^)
4. sobol2002 - implements the Monte Carlo estimation of the Sobol indices for both first-order and total indices at the same time (all together 2p indices), at a total cost of (p+2) × n model evaluations. These are called the Saltelli estimators. This estimator suffers from a conditioning problem when estimating the variances behind the indices computations. This can seriously affect the Sobol indices estimates in case of largely non-centered output. To avoid this effect, you have to center the model output before applying "sobol2002". Functions "soboljansen" and "sobolmartinez" do not suffer from this problem. (Saltelli^35^)
5. sobol2007 - implements the Monte Carlo estimation of the Sobol indices for both first-order and total indices at the same time (all together 2p indices), at a total cost of (p+2) × n model evaluations. These are called the Mauntz estimators. (Saltelli *et al.*^41^)
6. sobolmartinez - implements the Monte Carlo estimation of the Sobol indices for both first-order and total indices using correlation coefficients-based formulas, at a total cost of (p + 2) × n model evaluations. These are called the Martinez estimators.
7. sobol - implements the Monte Carlo estimation of the Sobol sensitivity indices. Allows the estimation of the indices of the variance decomposition up to a given order, at a total cost of (N + 1) × n where N is the number of indices to estimate. (Sobol’^21^)

**Fig. 13.**
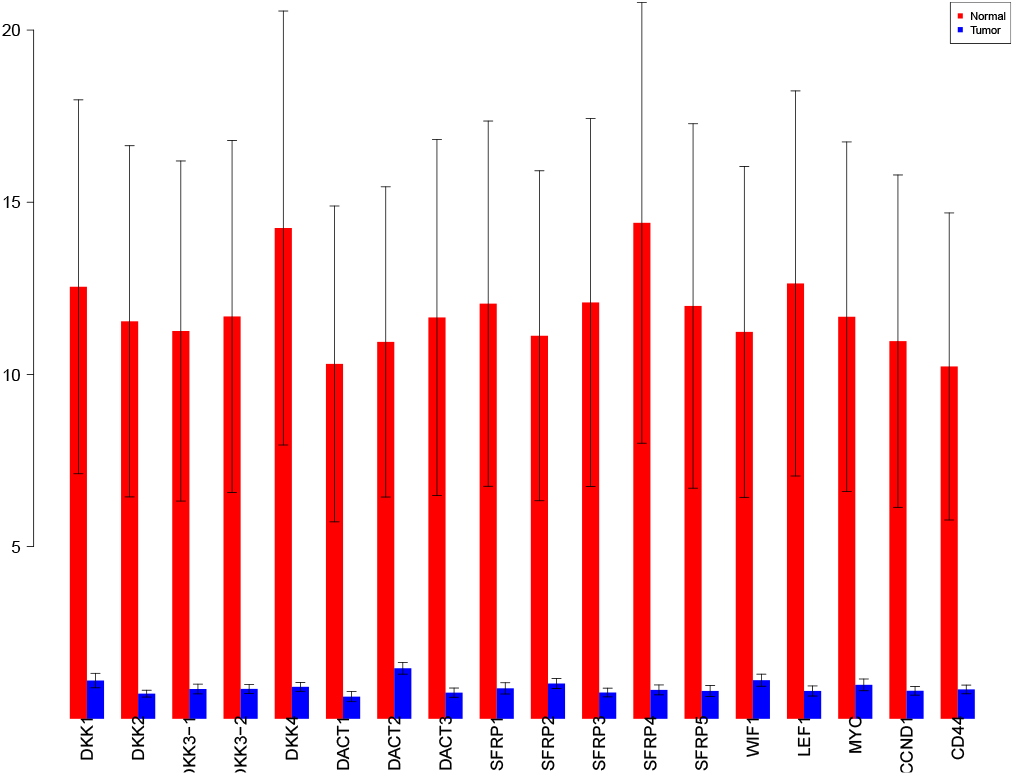
sensiFdiv indices using Total Variation distance. Red - indices for normal. Blue - indices for tumor.

**Fig. 14.**
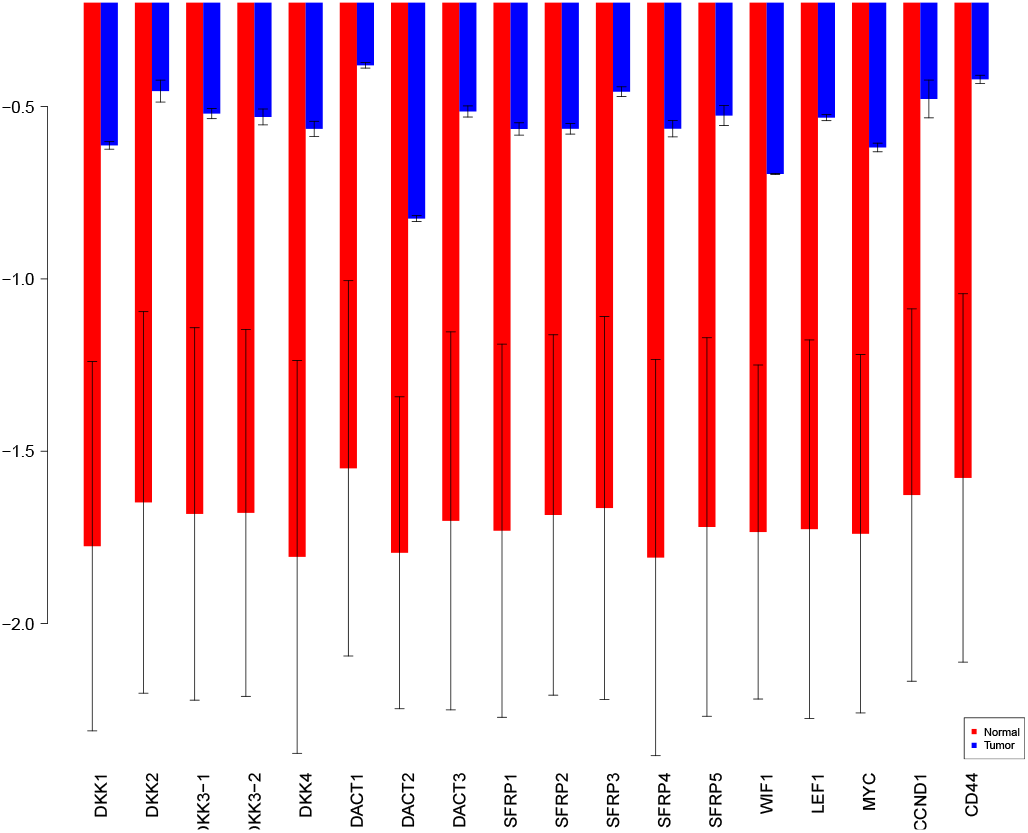
sensiFdiv indices for Kullback-Leibler divergence. Red - indices for normal. Blue - indices for tumor.

**Fig. 15.**
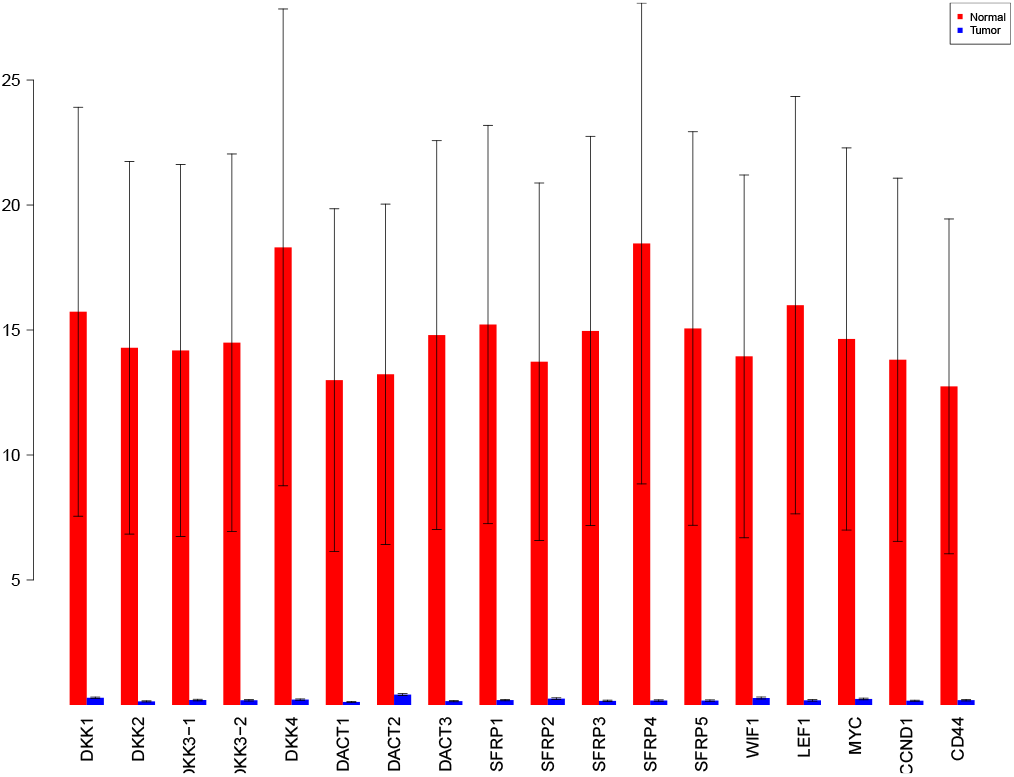
sensiFdiv indices for Hellinger distance. Red - indices for normal. Blue - indices for tumor.

**Fig. 16.**
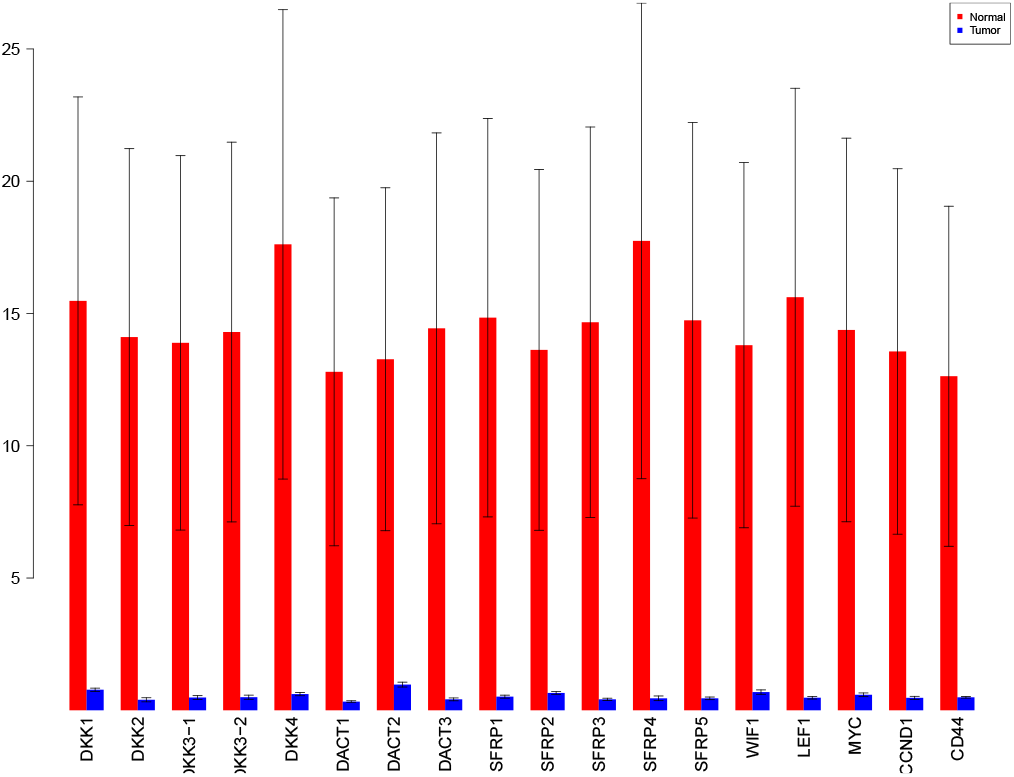
sensiFdiv indices for Pearson *χ*^2^ distance. Red - indices for normal. Blue - indices for tumor.

**Fig. 17.**
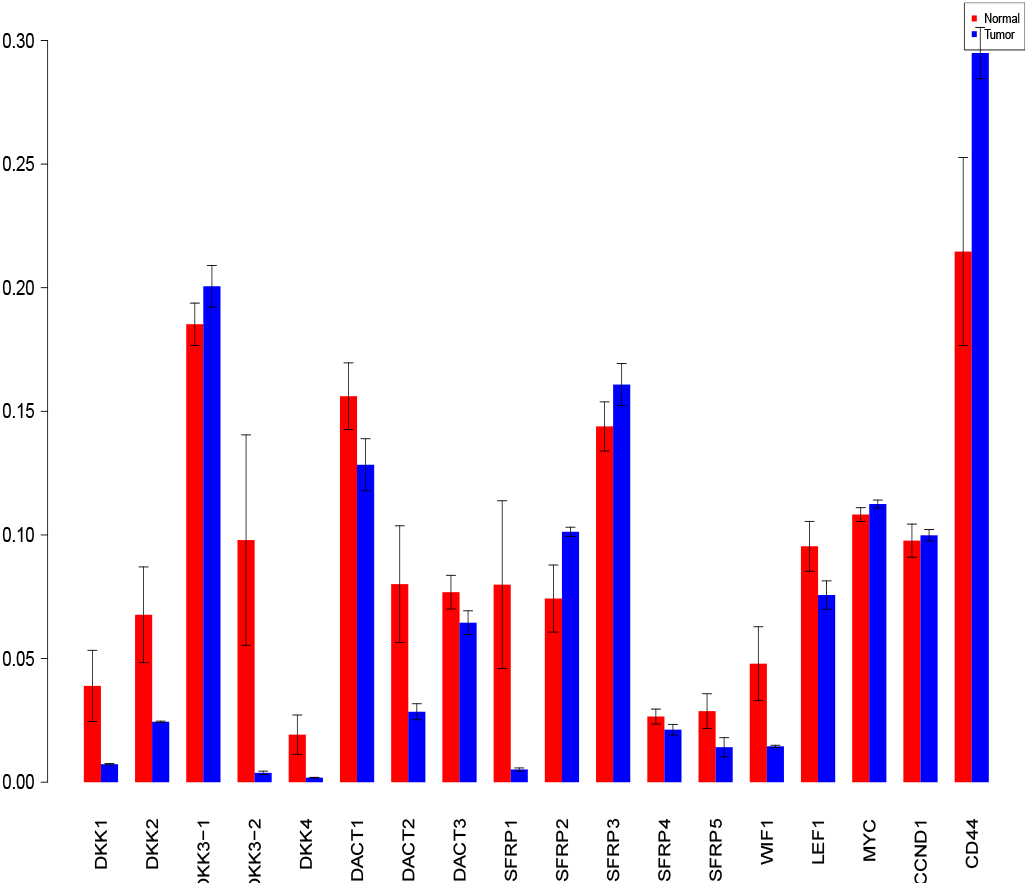
sensiHSIC indices for linear kernel. Red - indices for normal. Blue - indices for tumor.

**Fig. 18.**
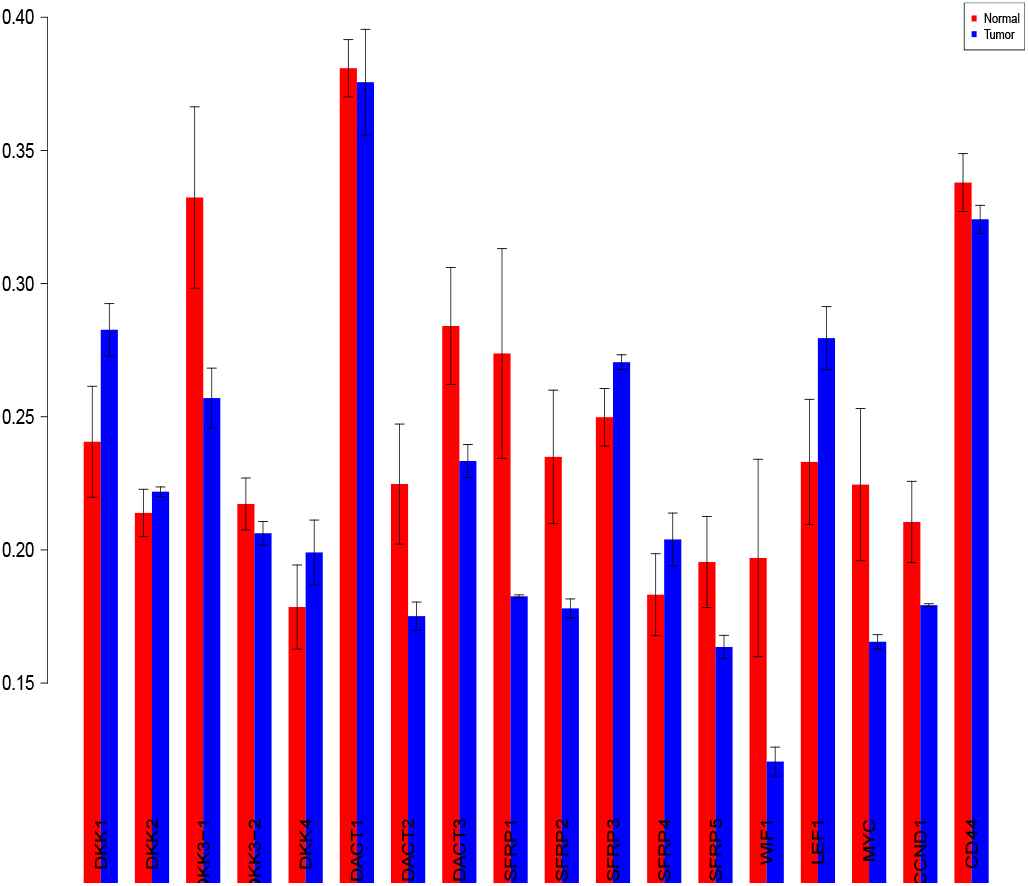
sensiHSIC indices for laplace kernel. Red - indices for normal. Blue - indices for tumor.

**Fig. 19.**
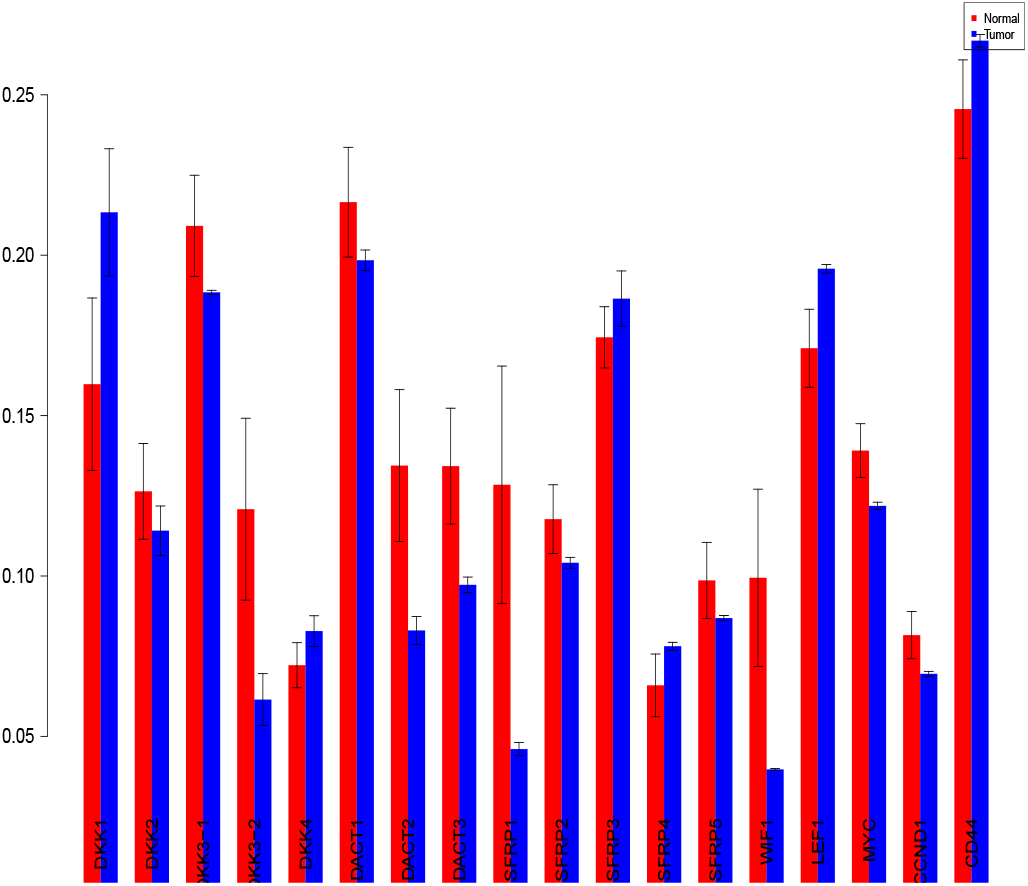
sensiHSIC indices for rbf kernel. Red - indices for normal. Blue - indices for tumor.

**Fig. 20.**
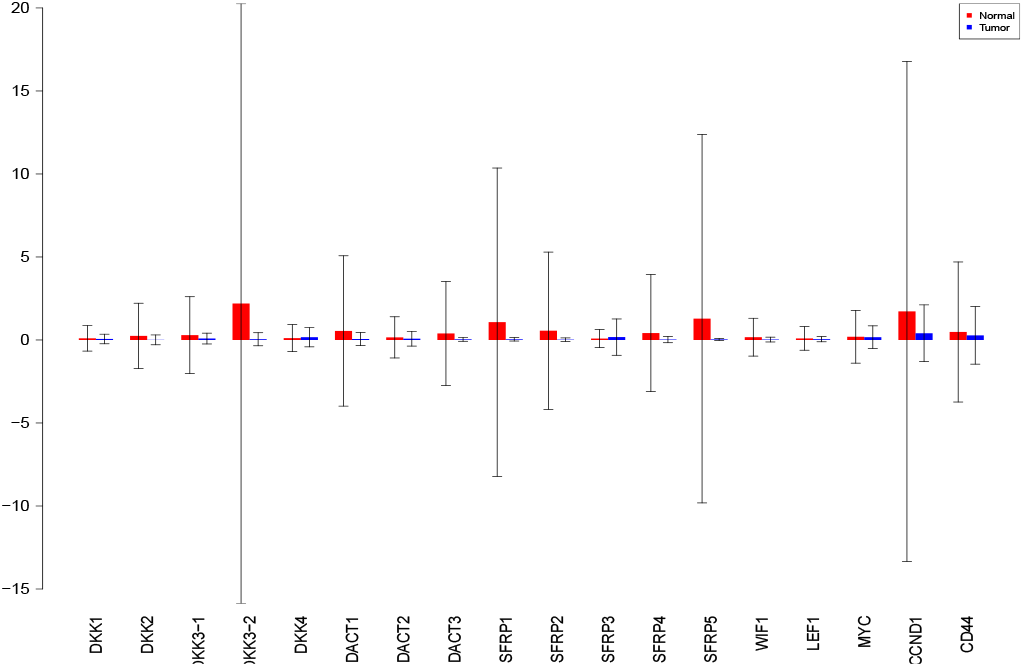
Sobol 2002 first order indices. Red - indices for normal. Blue -indices for tumor.

**Fig. 21.**
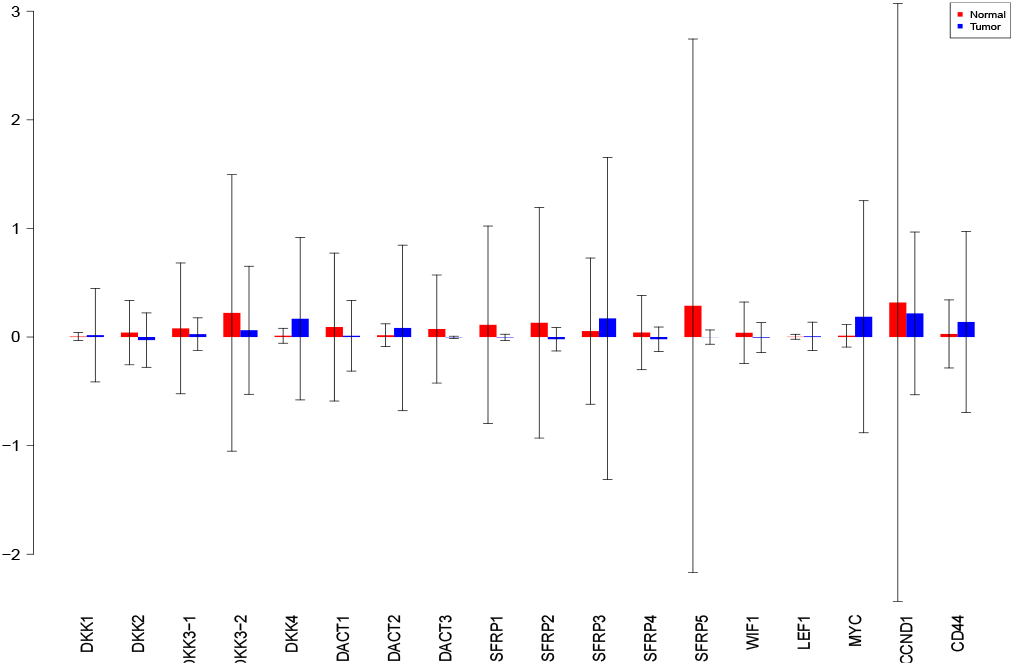
Sobol 2007 first order indices. Red - indices for normal. Blue -indices for tumor.

**Fig. 22.**
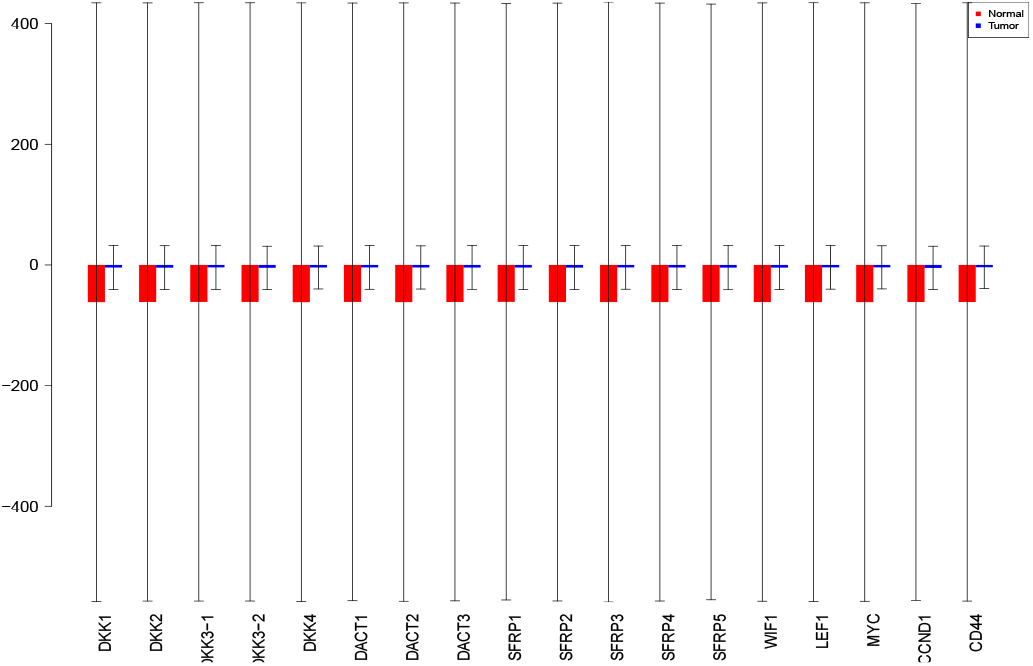
Sobol jansen first order indices. Red - indices for normal. Blue -indices for tumor.

**Fig. 23.**
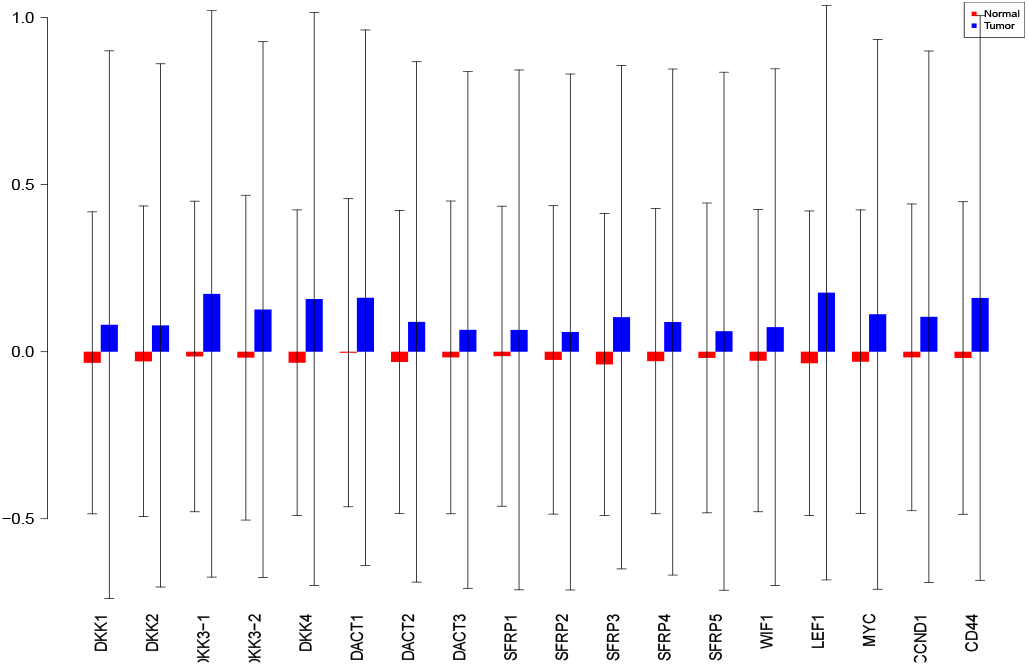
Sobol martinez first order indices. Red - indices for normal. Blue - indices for tumor.

**Fig. 24.**
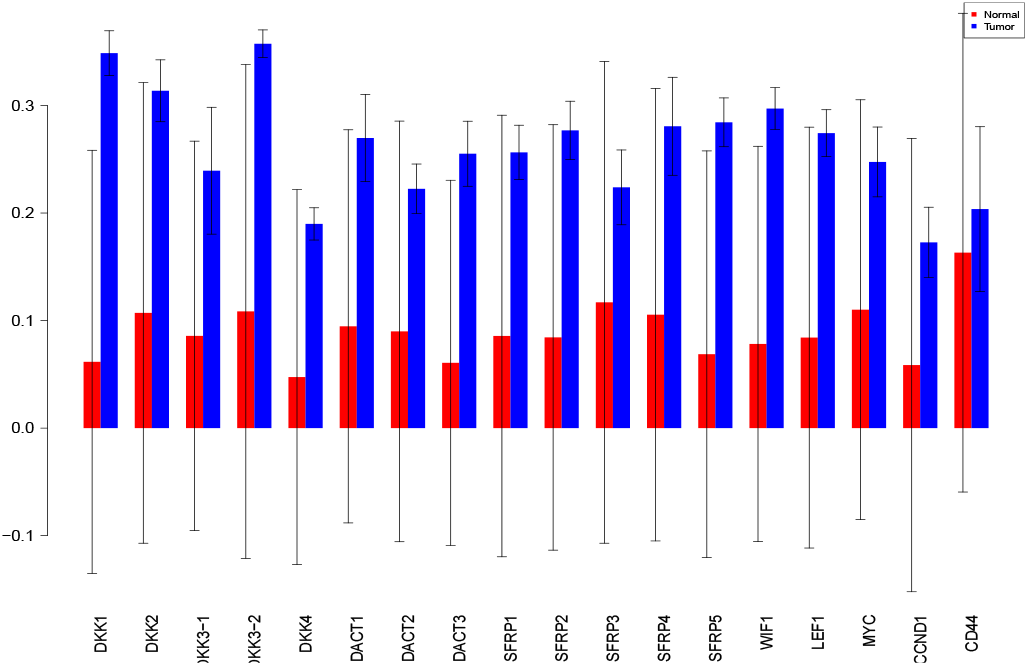
Sobol first order indices. Red - indices for normal. Blue - indices for tumor.

**Fig. 25.**
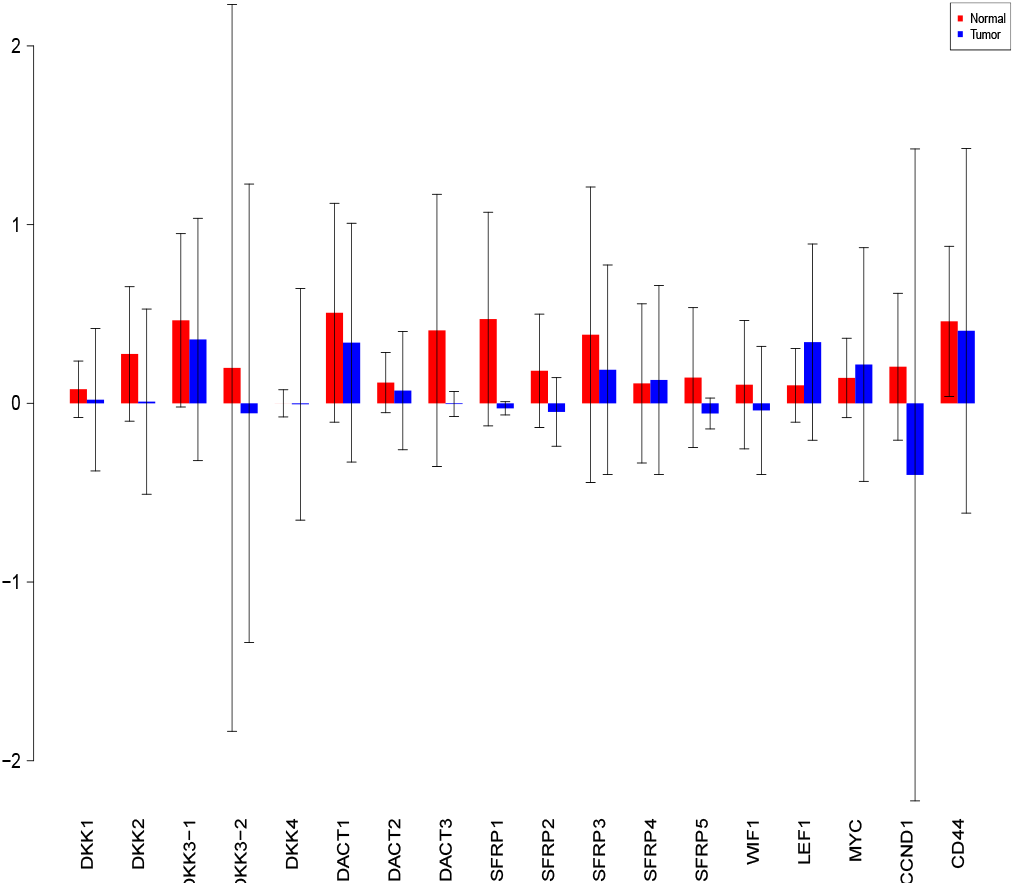
Sobol 2002 total order indices. Red - indices for normal. Blue -indices for tumor.

**Fig. 26.**
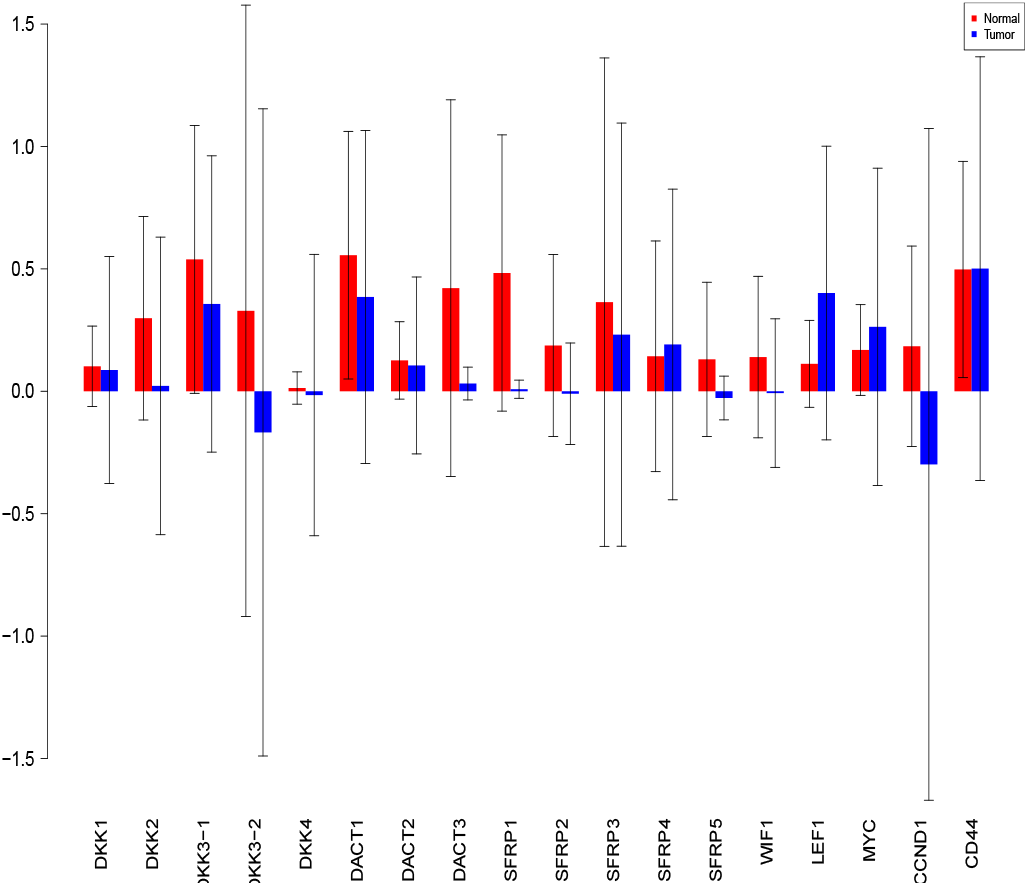
Sobol 2007 total order indices. Red - indices for normal. Blue -indices for tumor.

**Fig. 27.**
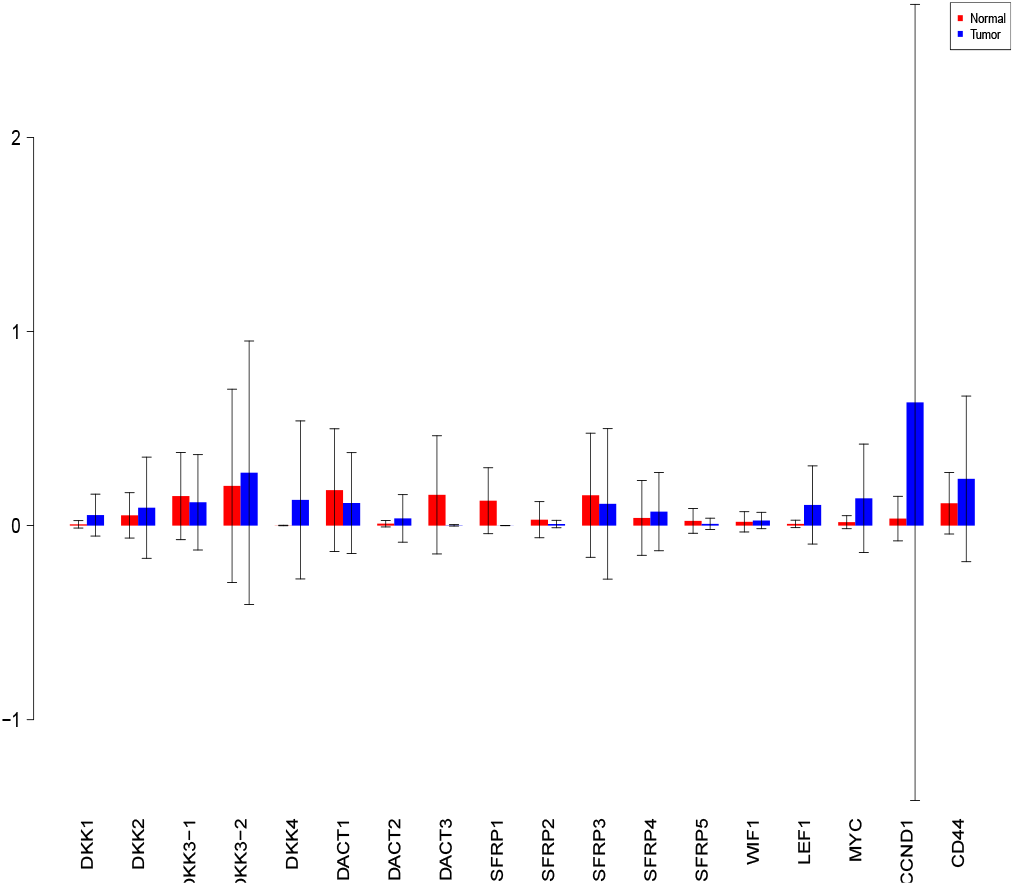
Sobol jansen total order indices. Red - indices for normal. Blue -indices for tumor.

**Fig. 28.**
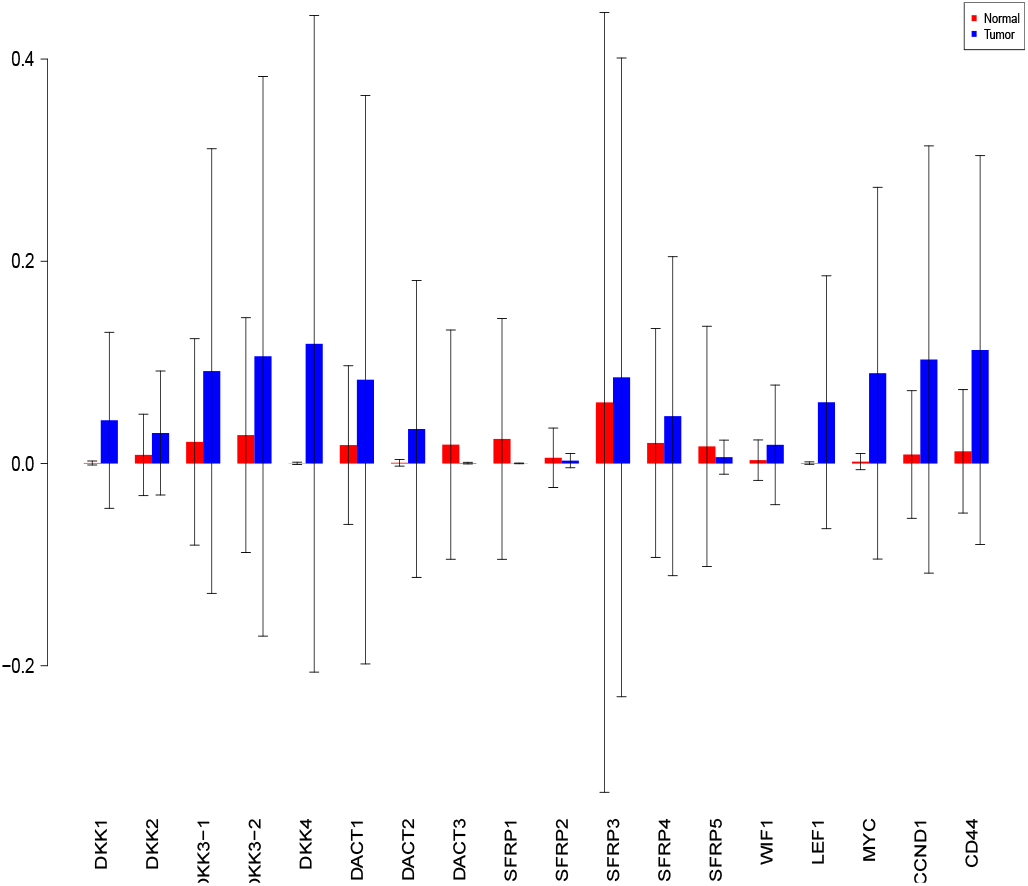
Sobol martinez total order indices. Red - indices for normal. Blue -indices for tumor.

**Fig. 29.**
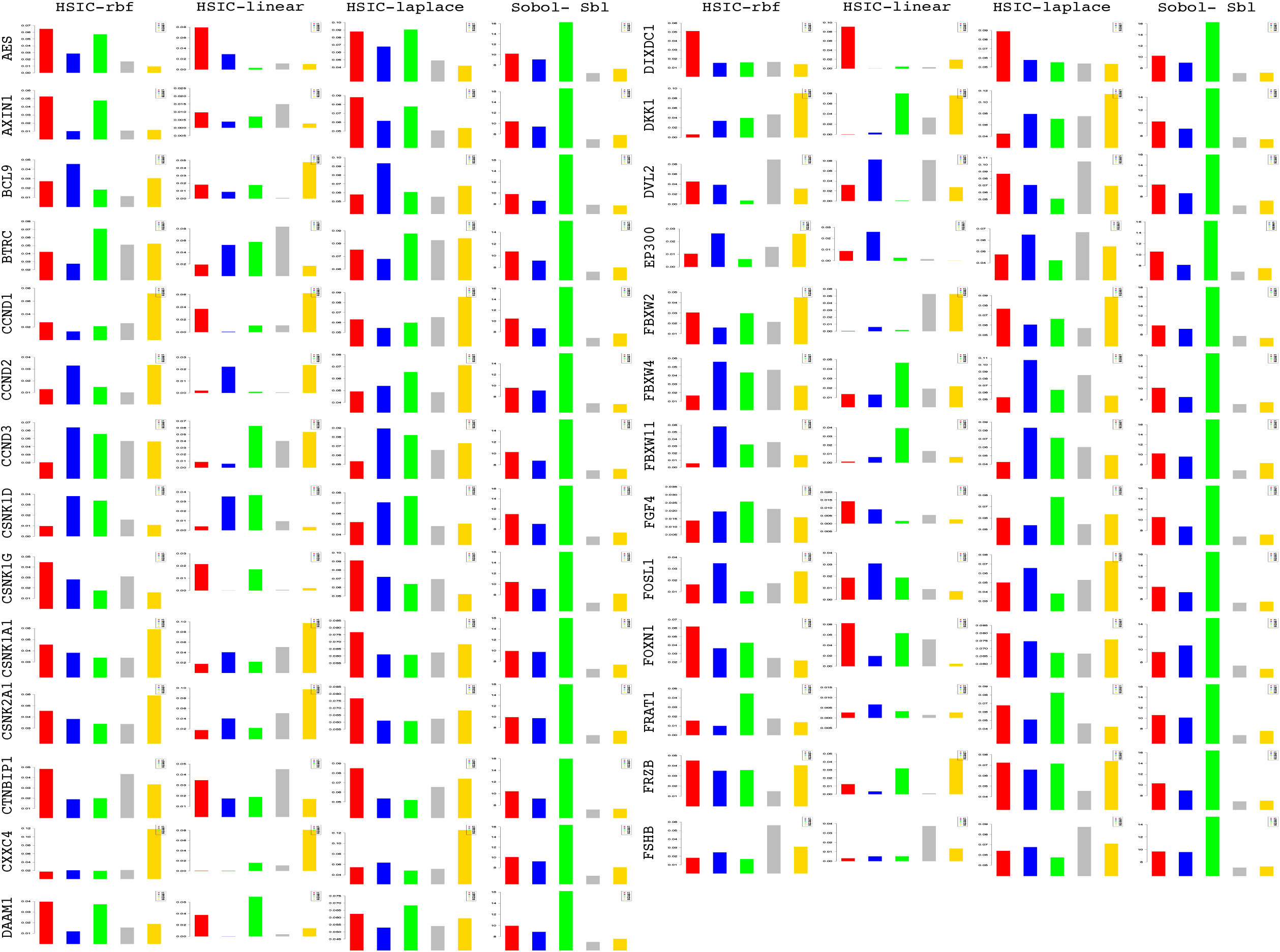
Column wise - methods to estimate sensitivity indices. Row wise - sensitivity indicies for each gene. For each graph, the bars represent sensitivity indices computed at t1 (red), t2 (blue), t3 (green), t4 (gray) and t5 (yellow). Indices were computed using non scaled time series data. TO - total order; FO - first order; SBL - Sobol

**Fig. 30.**
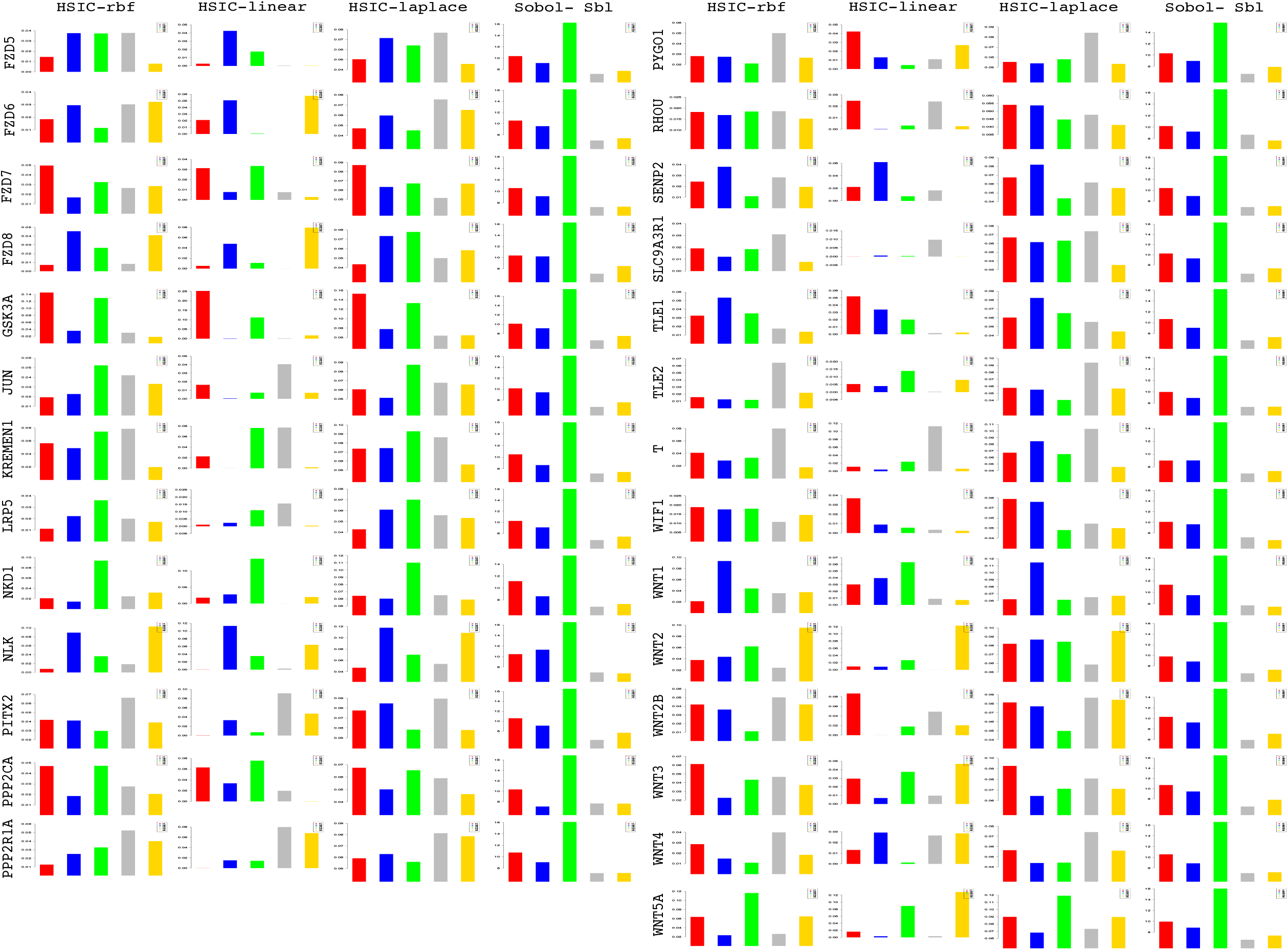
Column wise - methods to estimate sensitivity indices. Row wise - sensitivity indicies for each gene. For each graph, the bars represent sensitivity indices computed at t1 (red), t2 (blue), t3 (green), t4 (gray) and t5 (yellow). Indices were computed using non scaled time series data. TO - total order; FO - first order; SBL - Sobol

**Fig. 31.**
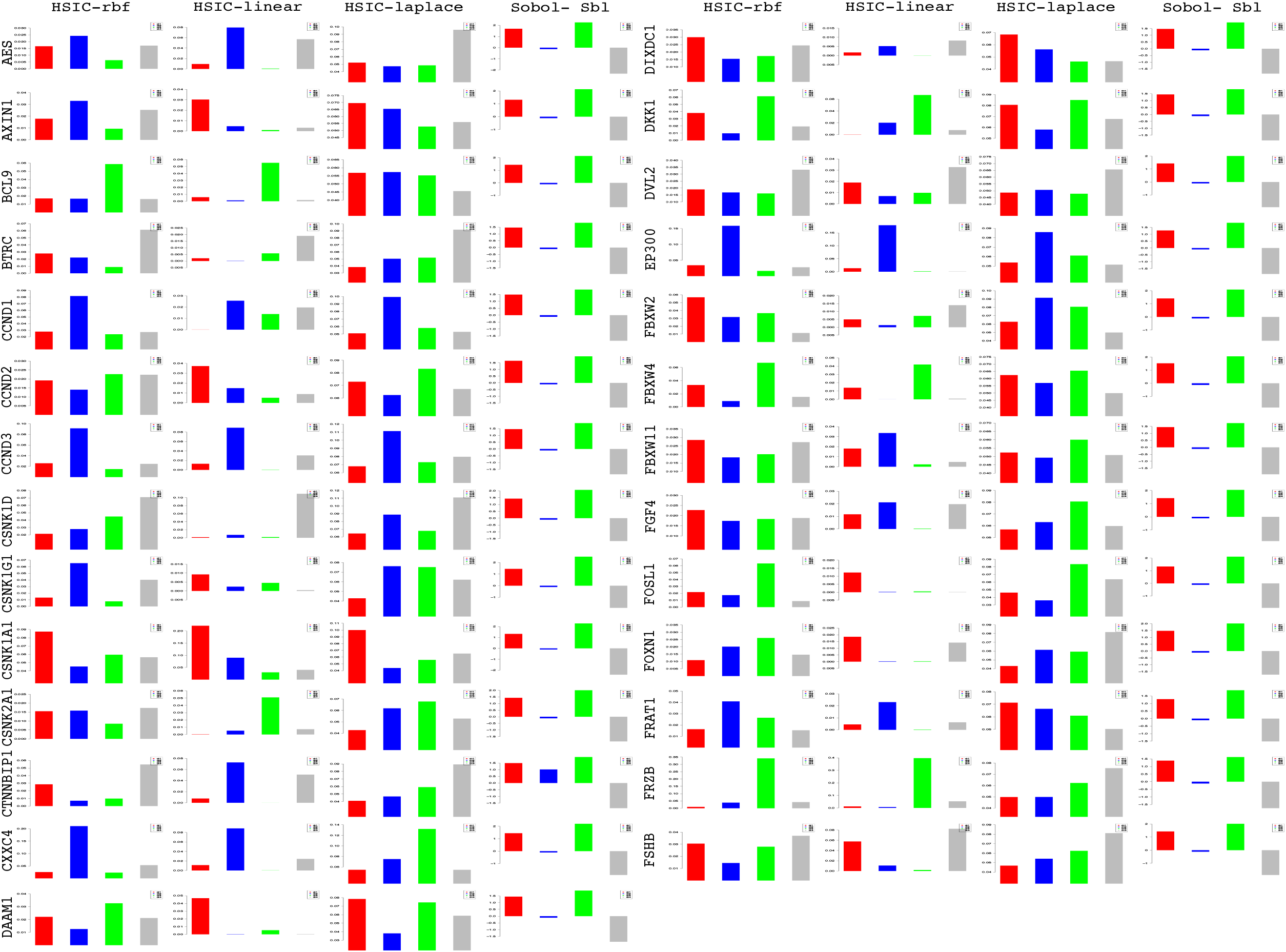
Column wise - methods to estimate sensitivity indices. Row wise - sensitivity indicies for each gene on deviations in fold change. For each graph, the bars represent sensitivity indices computed at <t1, t2> (red), <t2,t3> (blue), <t3,t4> (green) and <t4,t5> (gray). Indices were computed using non scaled time series data. TO - total order; FO -first order; SBL-Sobol

**Fig. 32.**
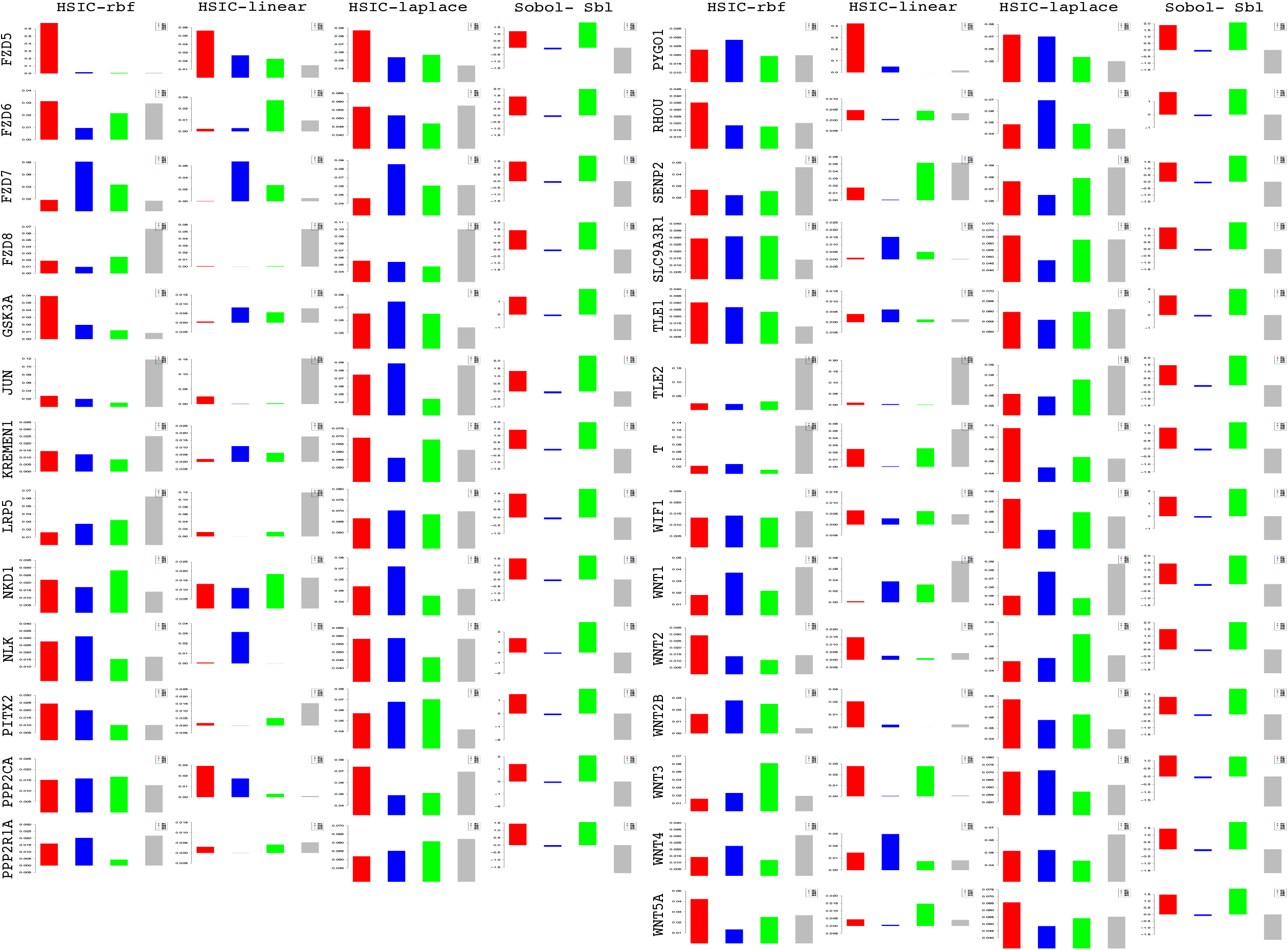
Column wise - methods to estimate sensitivity indices. Row wise - sensitivity indicies for each gene on deviations in fold change. For each graph, the bars represent sensitivity indices computed at <t1, t2> (red), <t2,t3> (blue), <t3,t4> (green) and <t4,t5> (gray). Indices were computed using non scaled time series data. TO - total order; FO -first order; SBL - Sobol

**Fig. 33.**
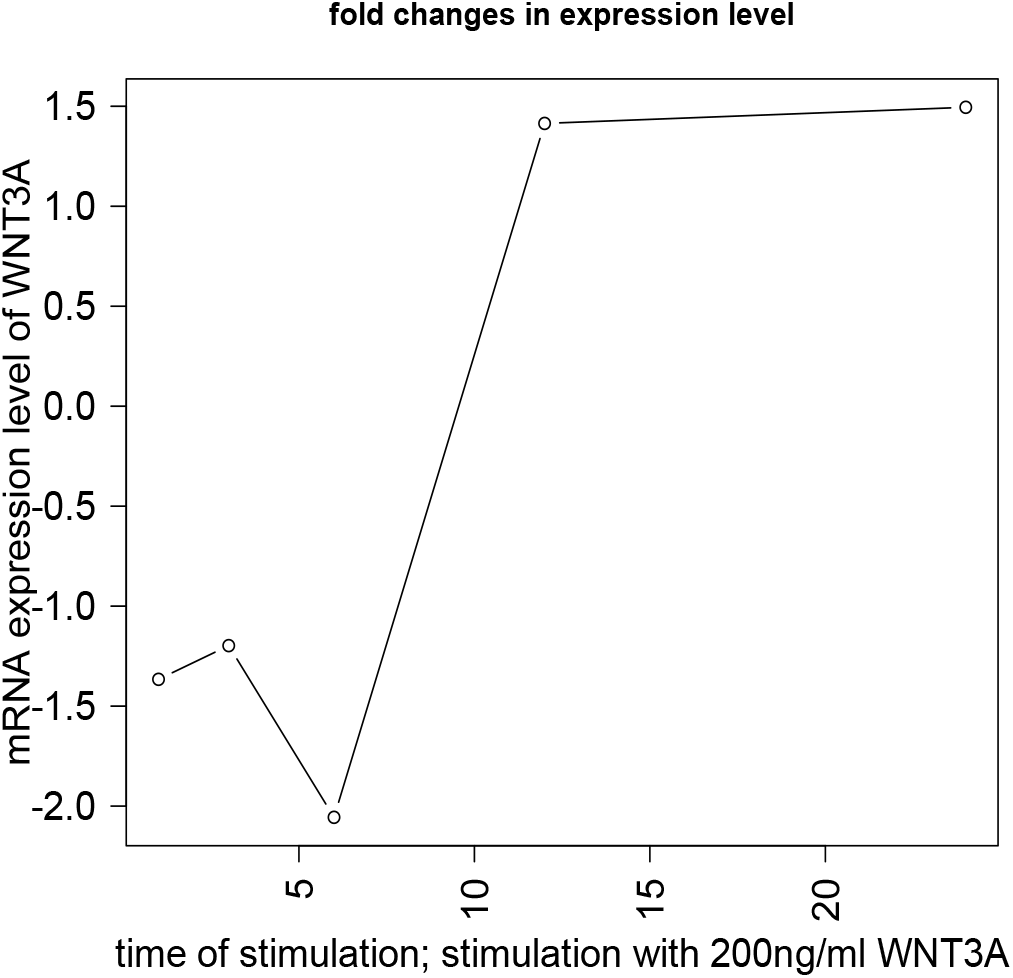
mRNA expression levels of *WNT*3*A* at 1^*st*^, 3^*rd*^, 6^*th*^, 12^*th*^ and 24^*th*^ hour from Gujral and MacBeath^17^.

**Fig. 34.**
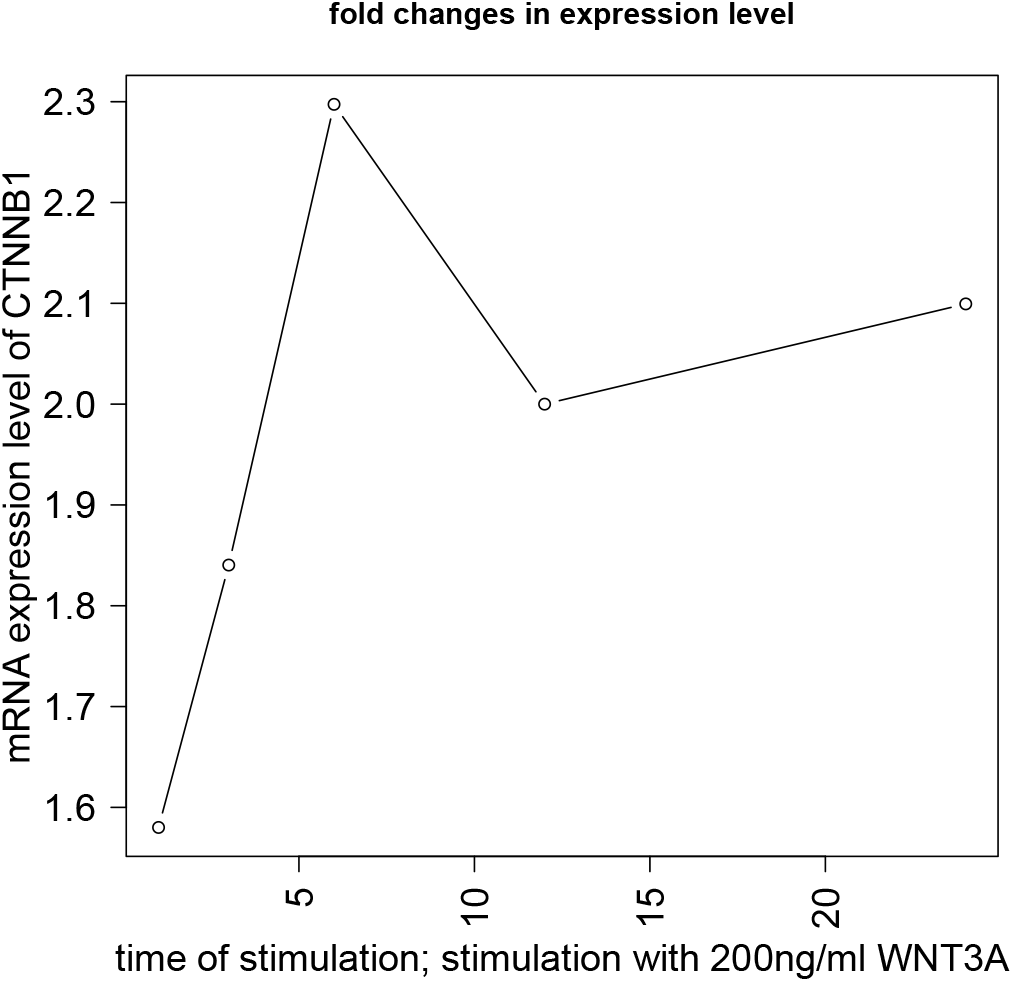
mRNA expression levels of CTNNB1 at 1^*st*^, 3^*rd*^, 6^*th*^, 12^*th*^ and 24^*th*^ hour from Gujral and MacBeath^17^.

**Fig. 35.**
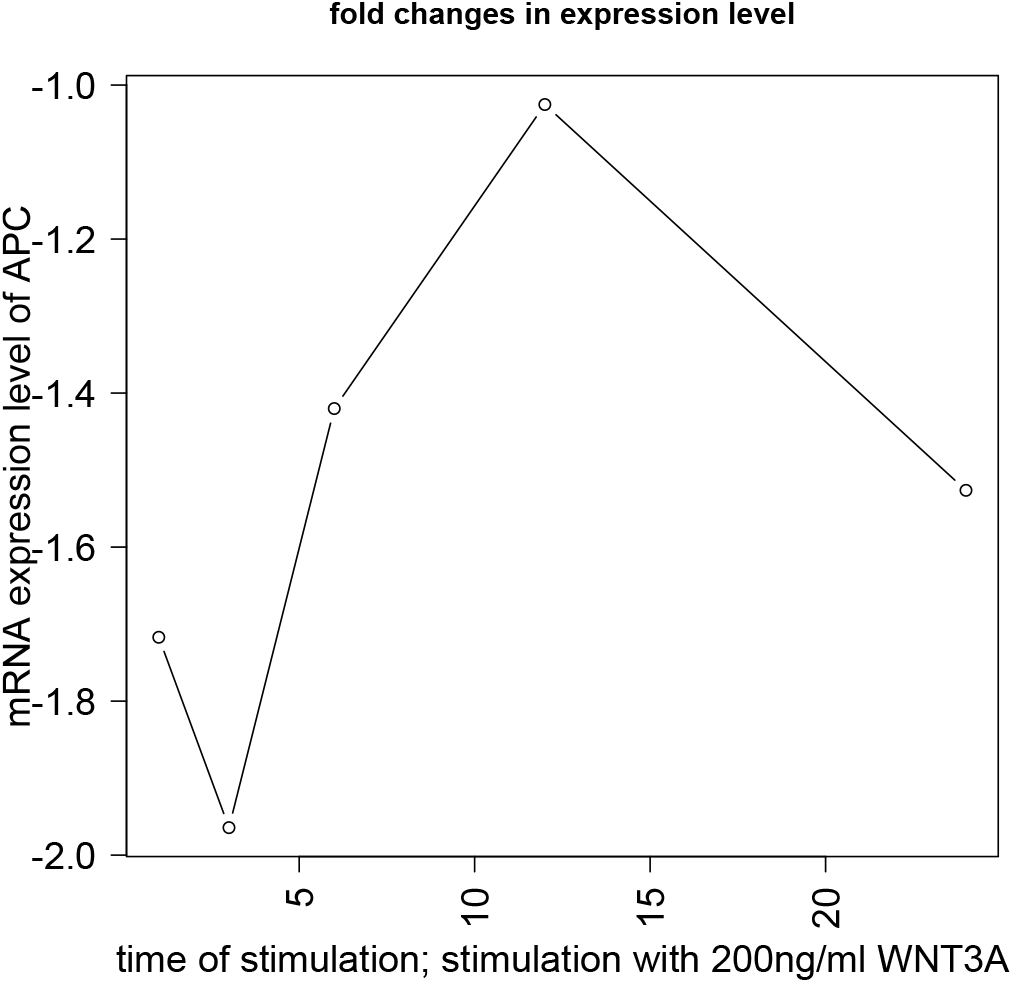
mRNA expression levels of APC at 1^*st*^, 3^*rd*^, 6^*th*^, 12^*th*^ and 24^*th*^ hour from Gujral and MacBeath^17^.

**Fig. 36.**
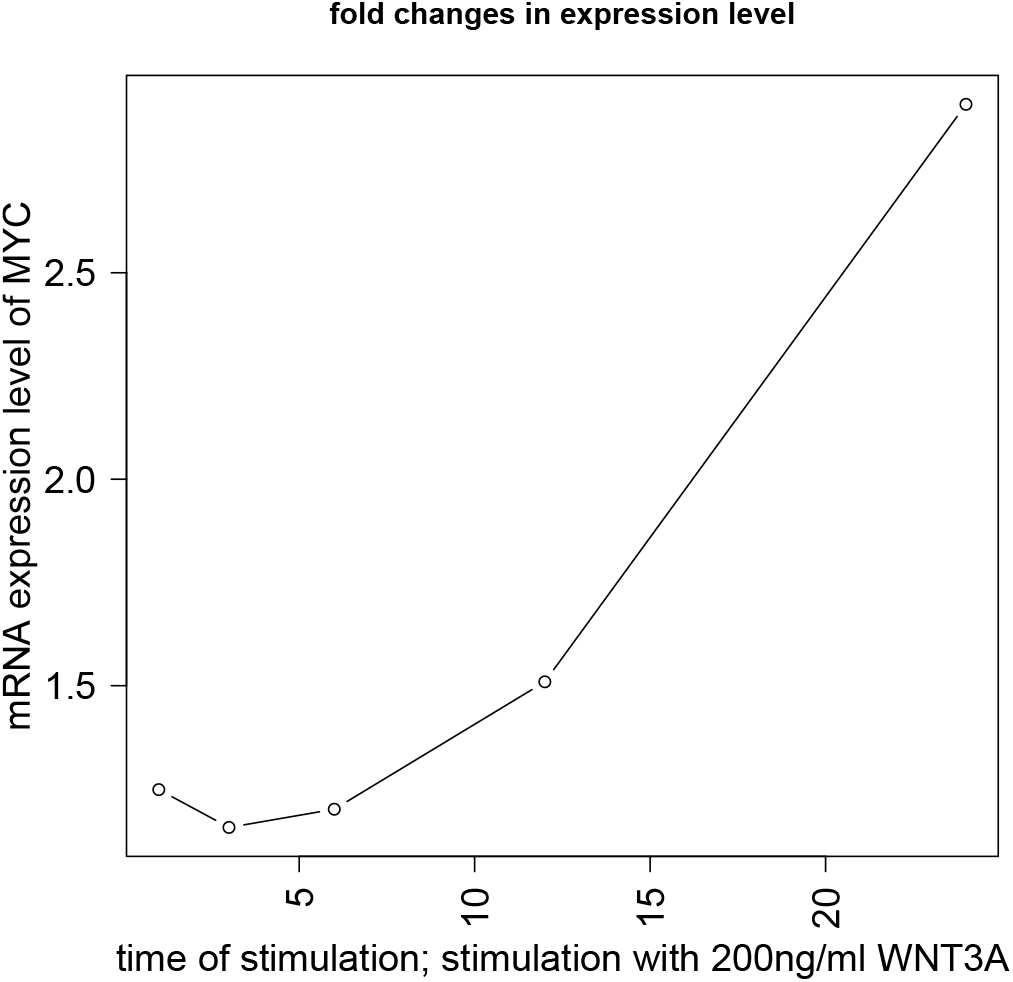
mRNA expression levels of MYC at 1^*st*^, 3^*rd*^, 6^*th*^, 12^*th*^ and 24^*th*^ hour from Gujral and MacBeath^17^.

**Fig. 37.**
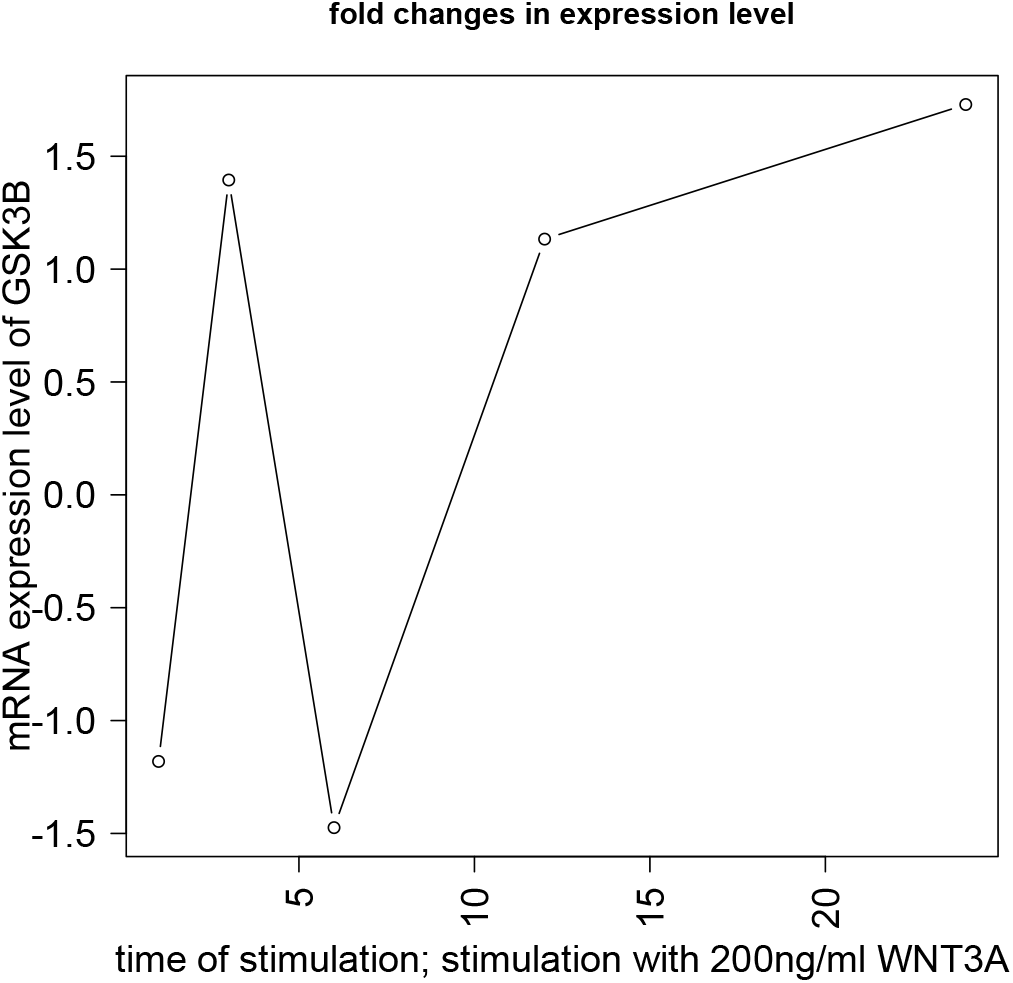
mRNA expression levels of *GSK*3*β* at 1^*st*^, 3^*rd*^, 6^*th*^, 12^*th*^ and 24^*th*^ hour from Gujral and MacBeath^17^.

**Fig. 38.**
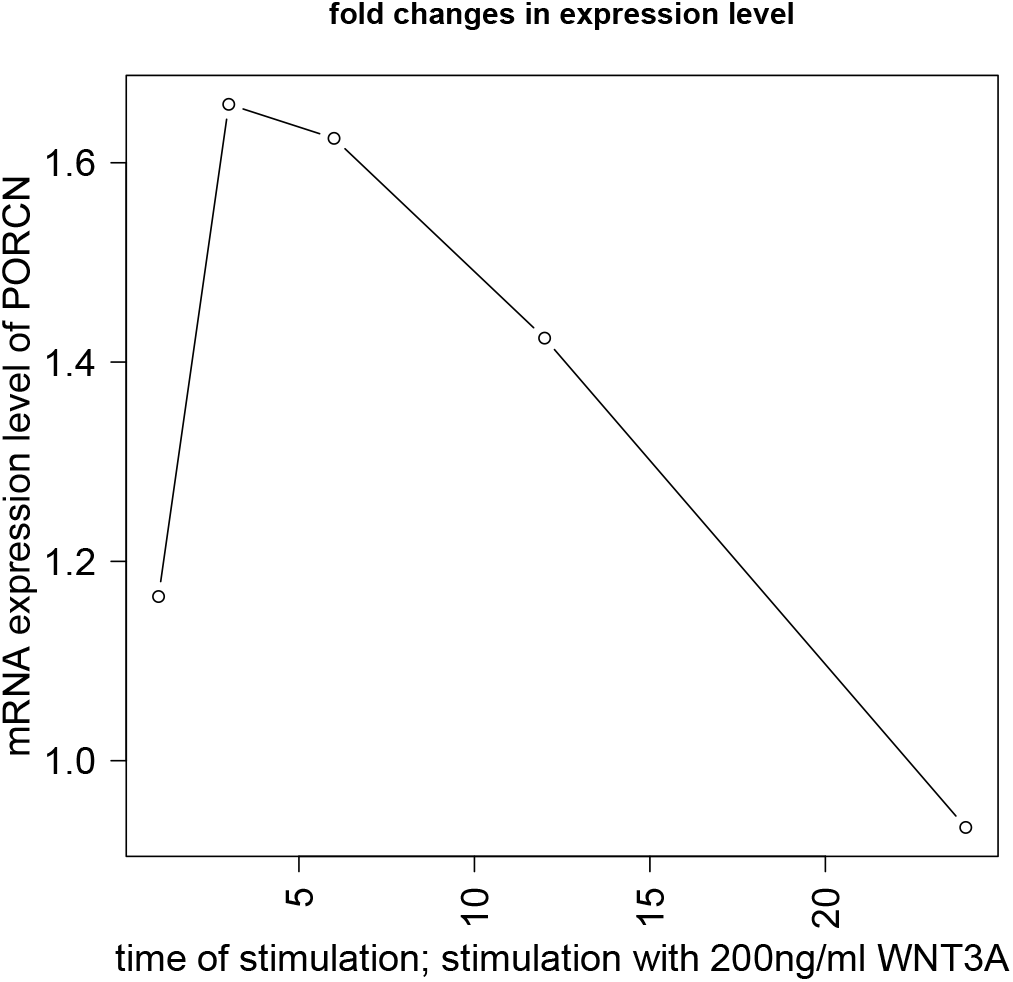
mRNA expression levels of *PORCN* at 1^*st*^, 3^*rd*^, 6^*th*^, 12^*th*^ and 24^*th*^ hour from Gujral and MacBeath^17^.

**Fig. 39.**
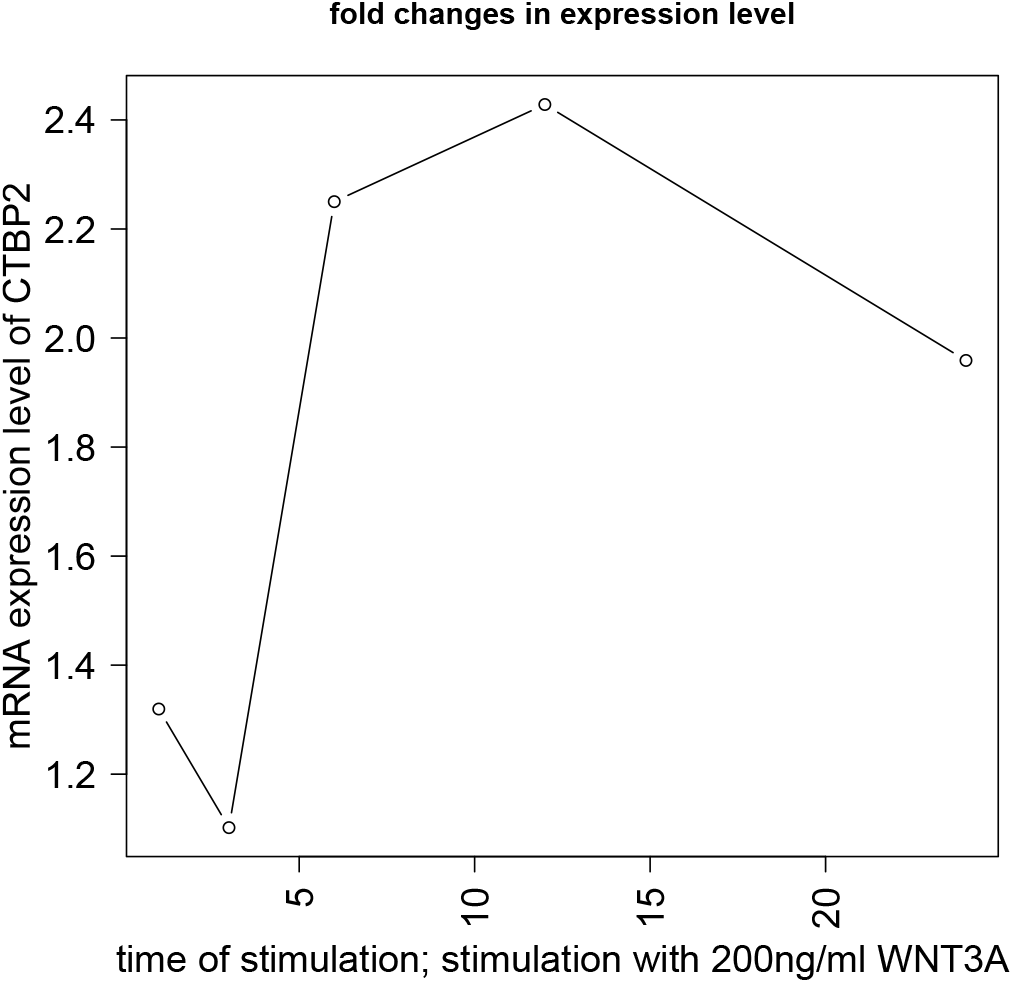
mRNA expression levels of *CTBP*2 at 1^*st*^, 3^*rd*^, 6^*th*^, 12^*th*^ and 24^*th*^ hour from Gujral and MacBeath^17^.

**Fig. 40.**
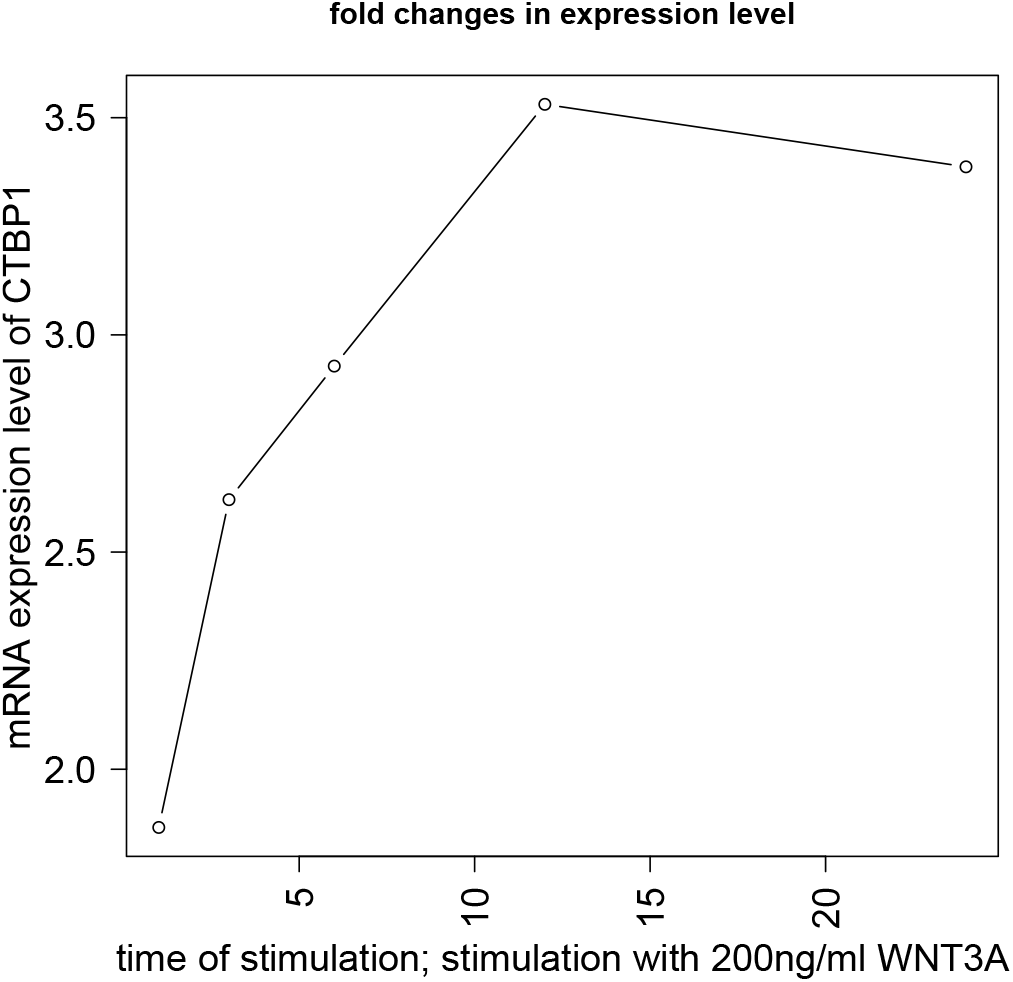
mRNA expression levels of *CTBP*1 at 1^*st*^, 3^*rd*^, 6^*th*^, 12^*th*^ and 24^*th*^ hour from Gujral and MacBeath^17^.

**Fig. 41.**
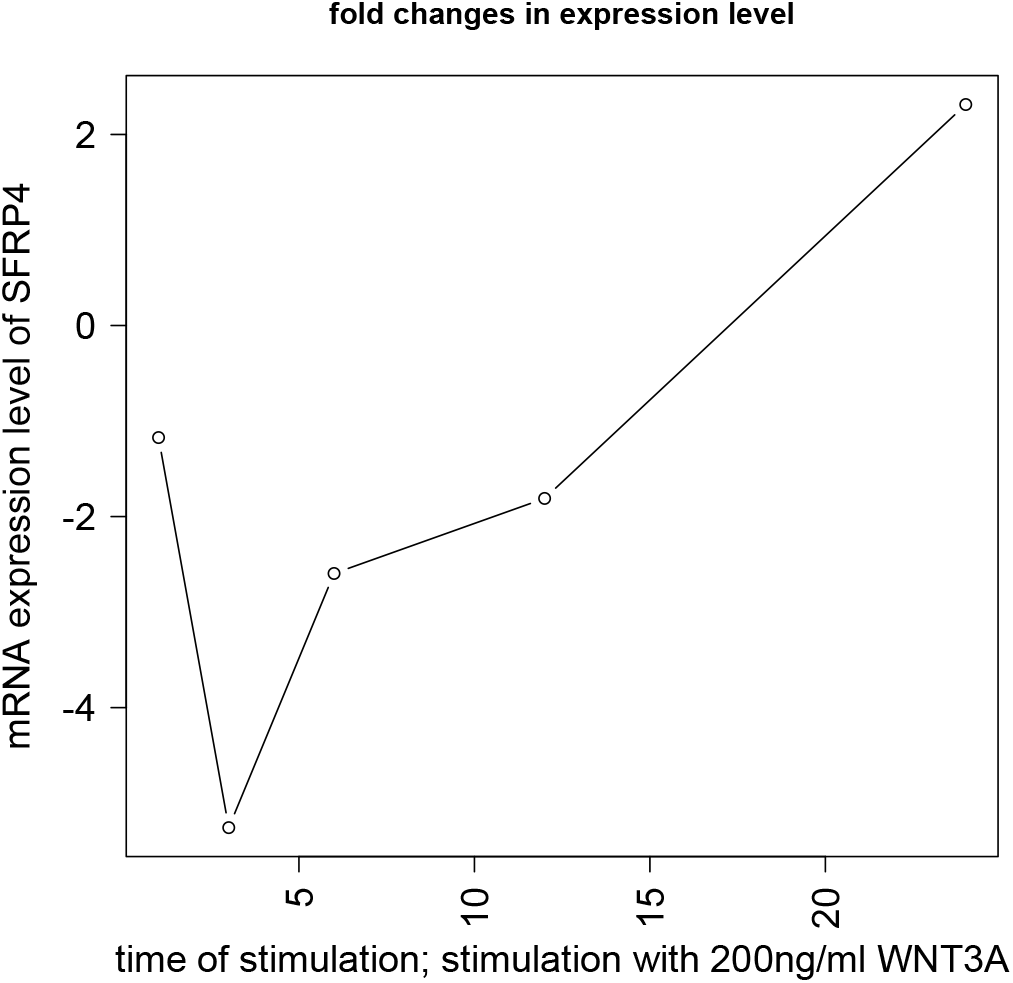
mRNA expression levels of *SFRP*4 at 1^*st*^, 3^*rd*^, 6^*th*^, 12^*th*^ and 24^*th*^ hour from Gujral and MacBeath^17^.

**Fig. 42.**
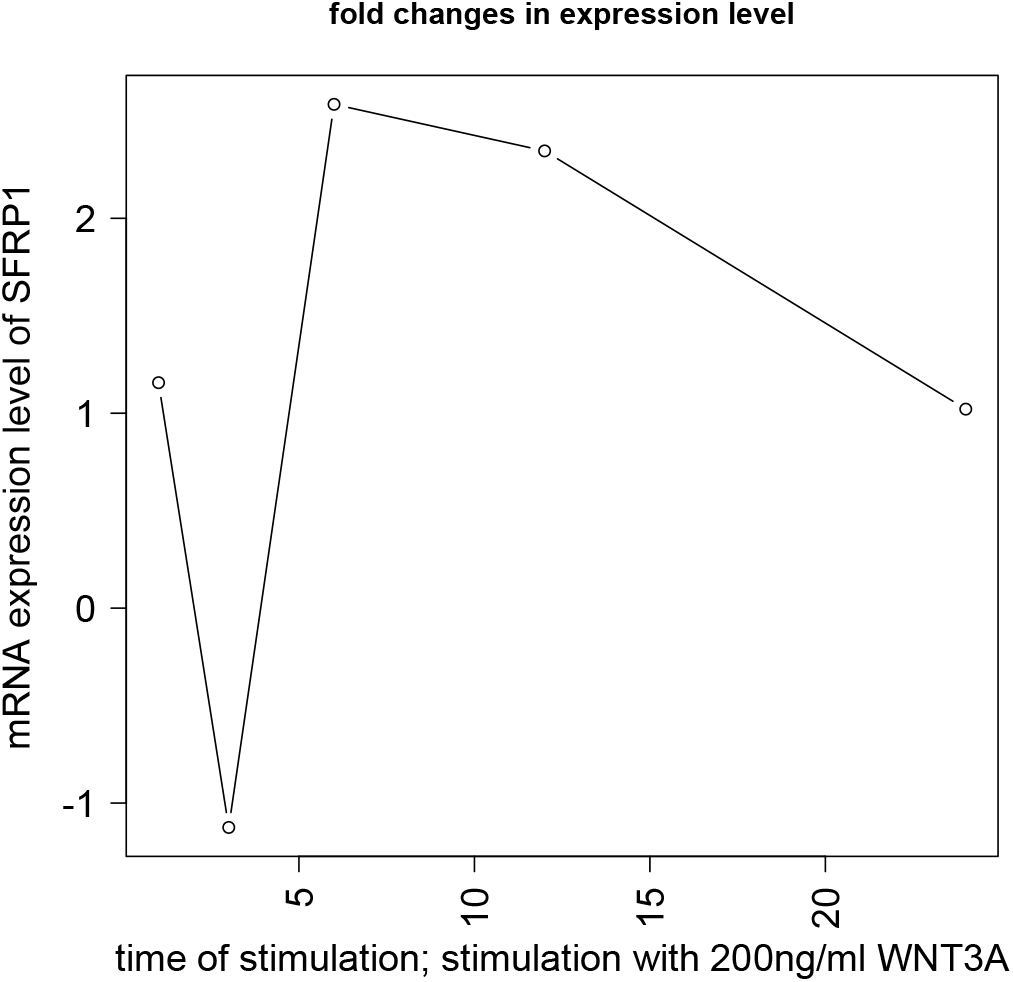
mRNA expression levels of *SFRP*1 at 1^*st*^, 3^*rd*^, 6^*th*^, 12^*th*^ and 24^*th*^ hour from Gujral and MacBeath^17^.

**Fig. 43.**
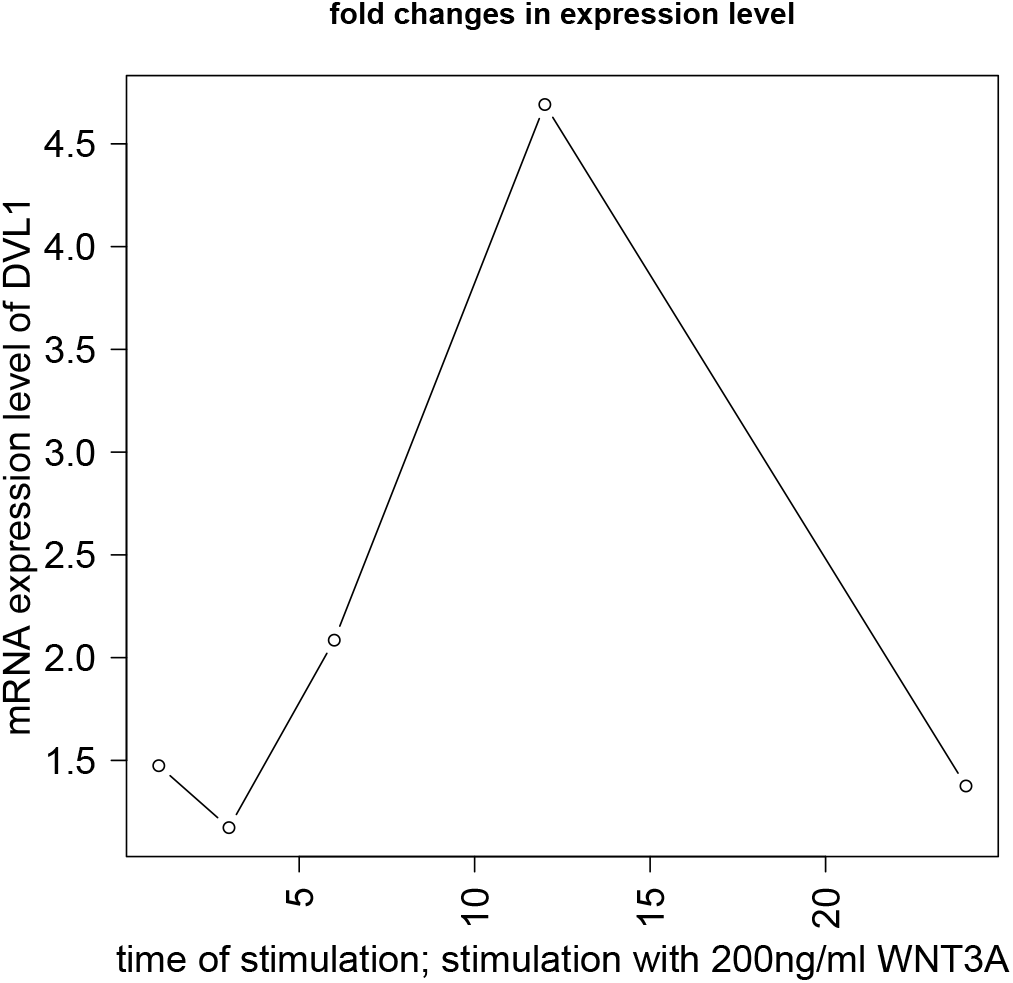
mRNA expression levels of *DVL*1 at 1^*st*^, 3^*rd*^, 6^*th*^, 12^*th*^ and 24^*th*^ hour from Gujral and MacBeath^17^.

**Fig. 44.**
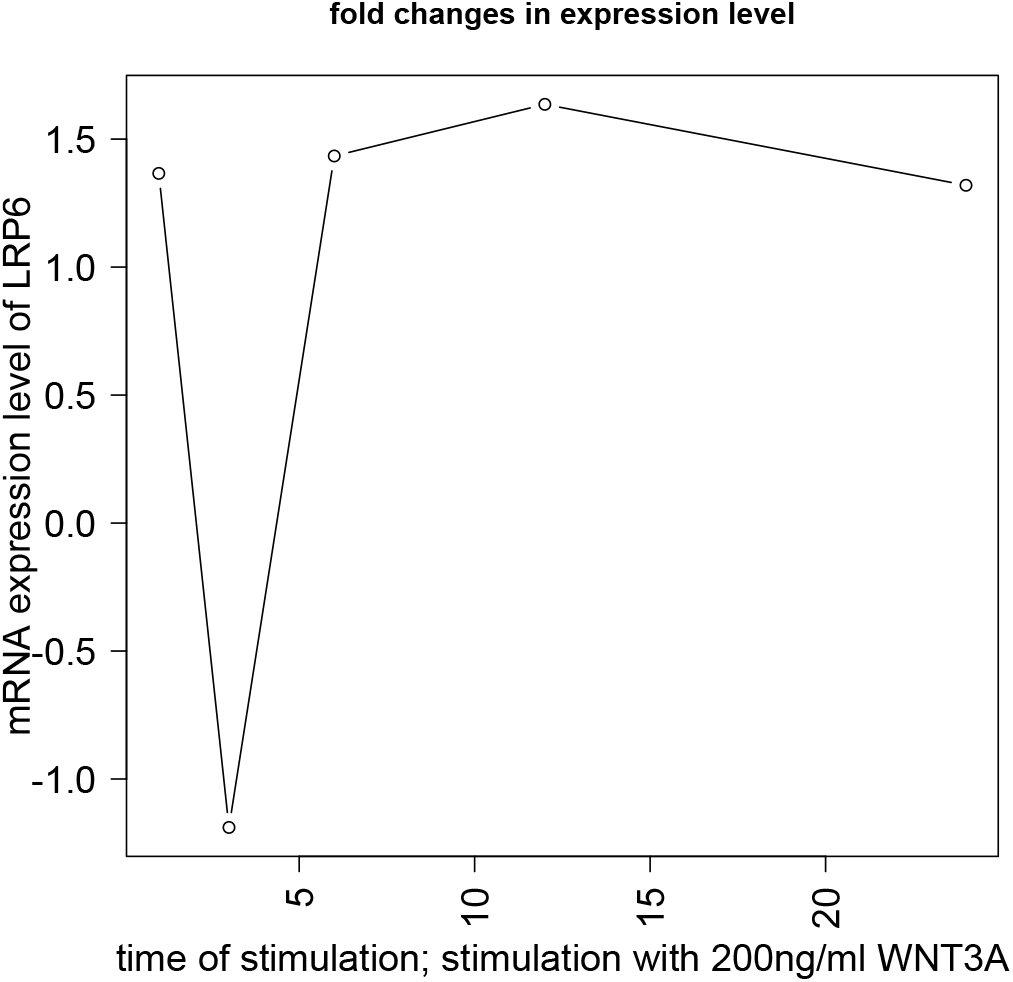
mRNA expression levels of *LRP*6 at 1^*st*^, 3^*rd*^, 6^*th*^, 12^*th*^ and 24^*th*^ hour from Gujral and MacBeath^17^.

**Fig. 45.**
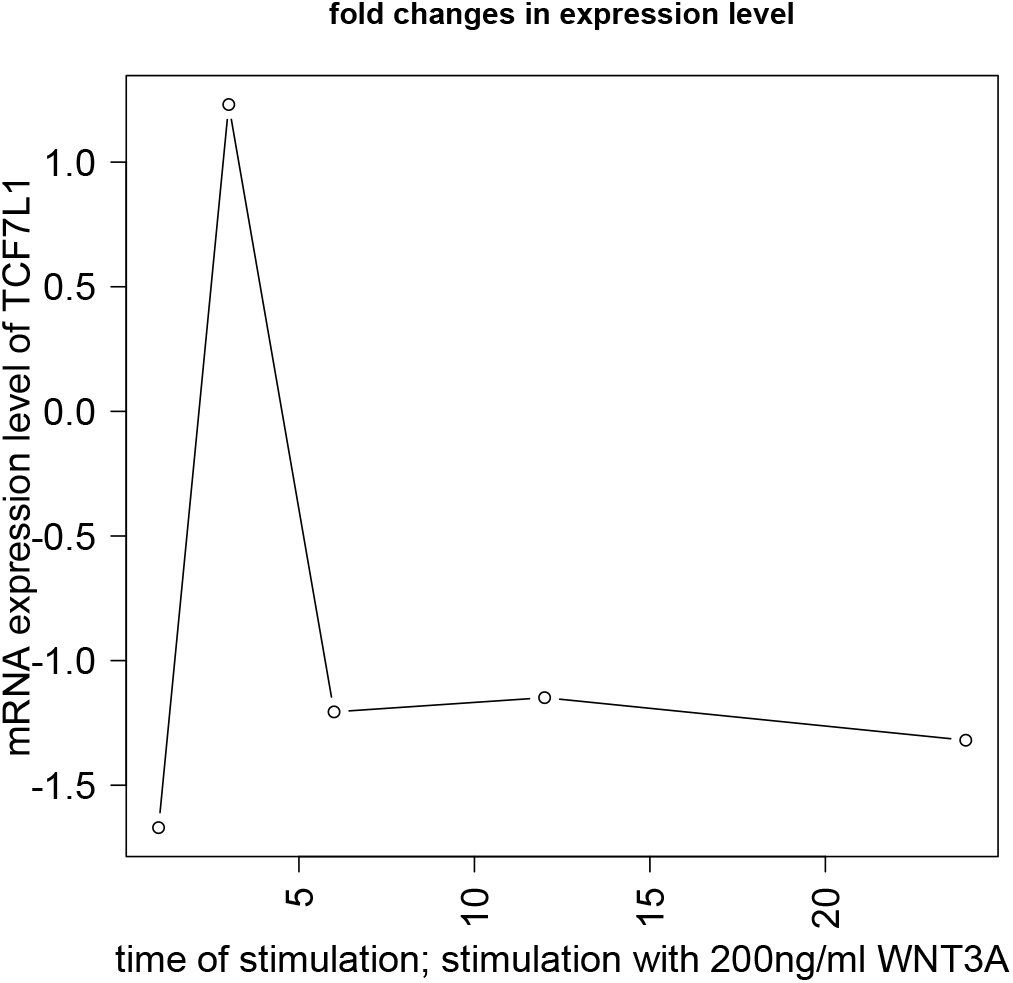
mRNA expression levels of *TCF*7*L*1 at 1^*st*^, 3^*rd*^, 6^*th*^, 12^*th*^ and 24^*th*^ hour from Gujral and MacBeath^17^.

**Fig. 46.**
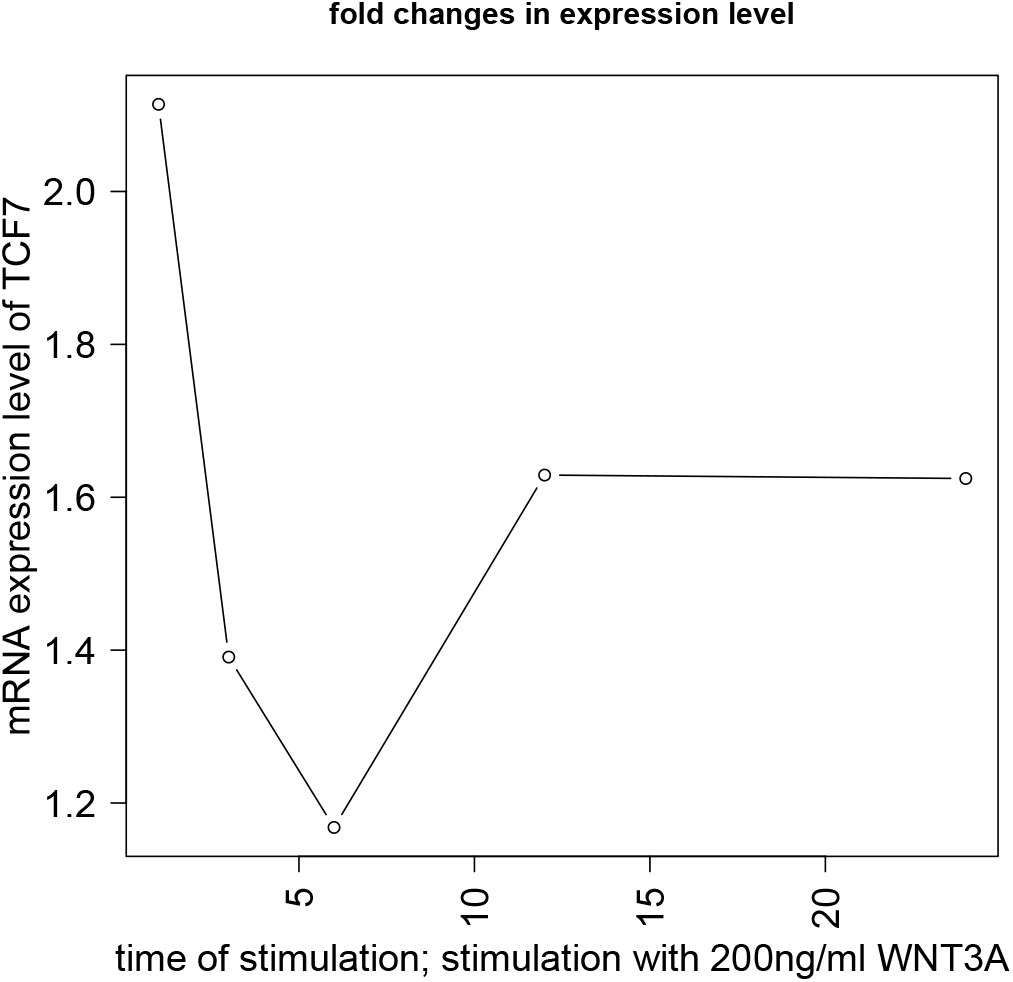
mRNA expression levels of *TCF*1 at 1^*st*^, 3^*rd*^, 6^*th*^, 12^*th*^ and 24^*th*^ hour from Gujral and MacBeath^17^.

**Fig. 47.**
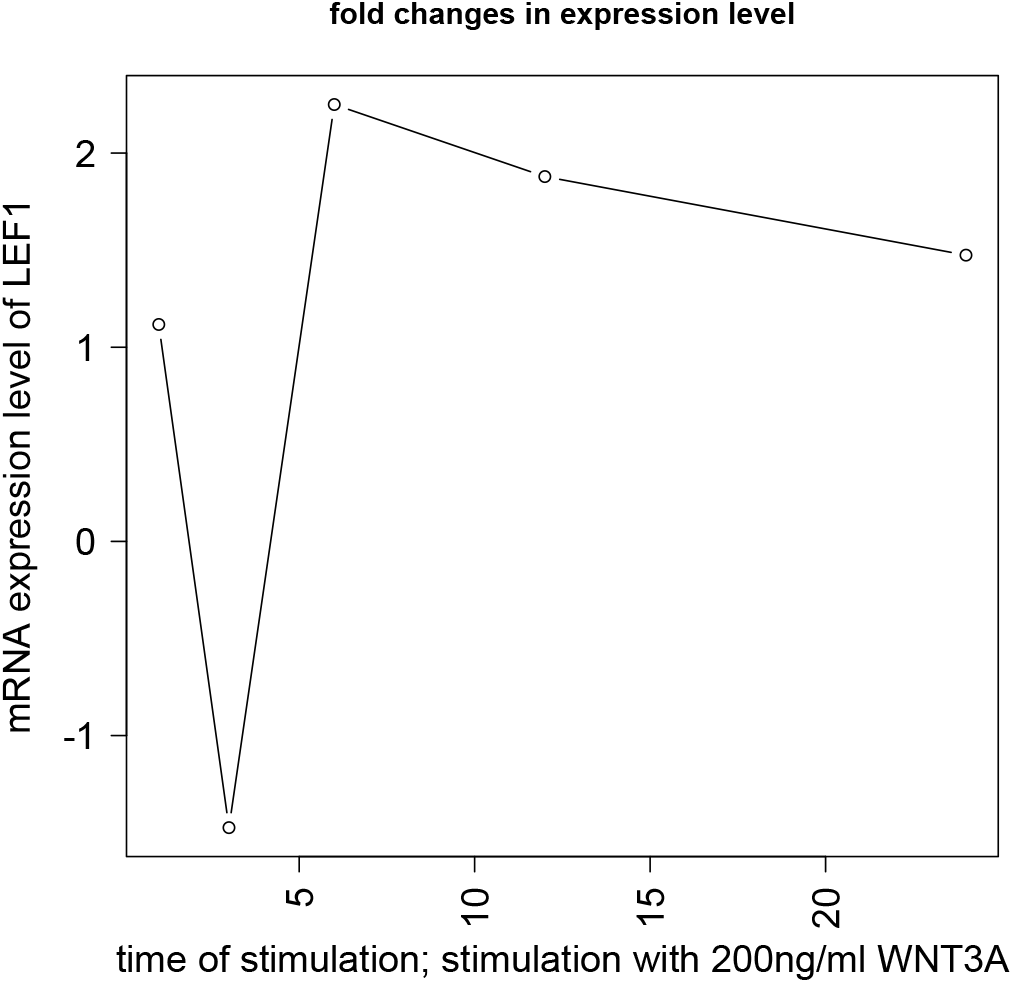
mRNA expression levels of *LEF*1 at 1^*st*^, 3^*rd*^, 6^*th*^, 12^*th*^ and 24^*th*^ hour from Gujral and MacBeath^17^.

**Fig. 48.**
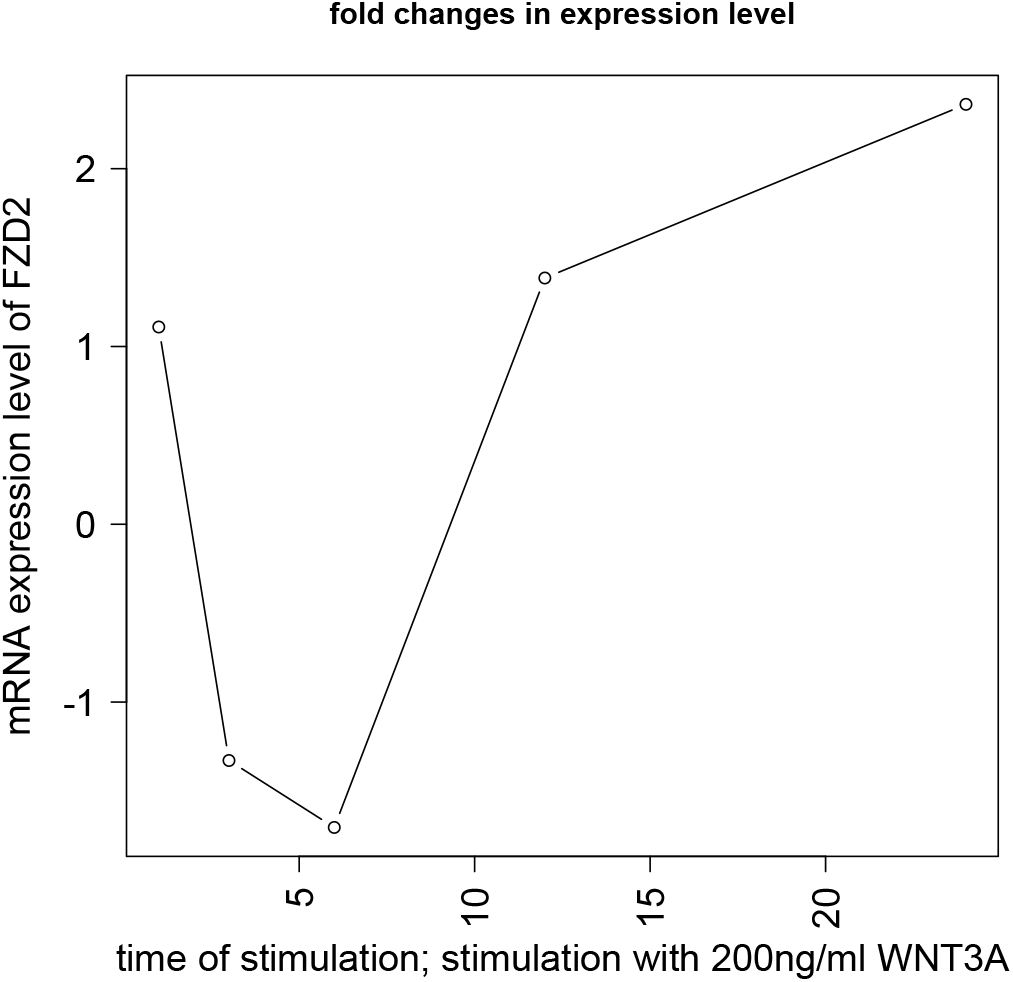
mRNA expression levels of *FZD*2 at 1^*st*^, 3^*rd*^, 6^*th*^, 12^*th*^ and 24^*th*^ hour from Gujral and MacBeath^17^.

**Fig. 49.**
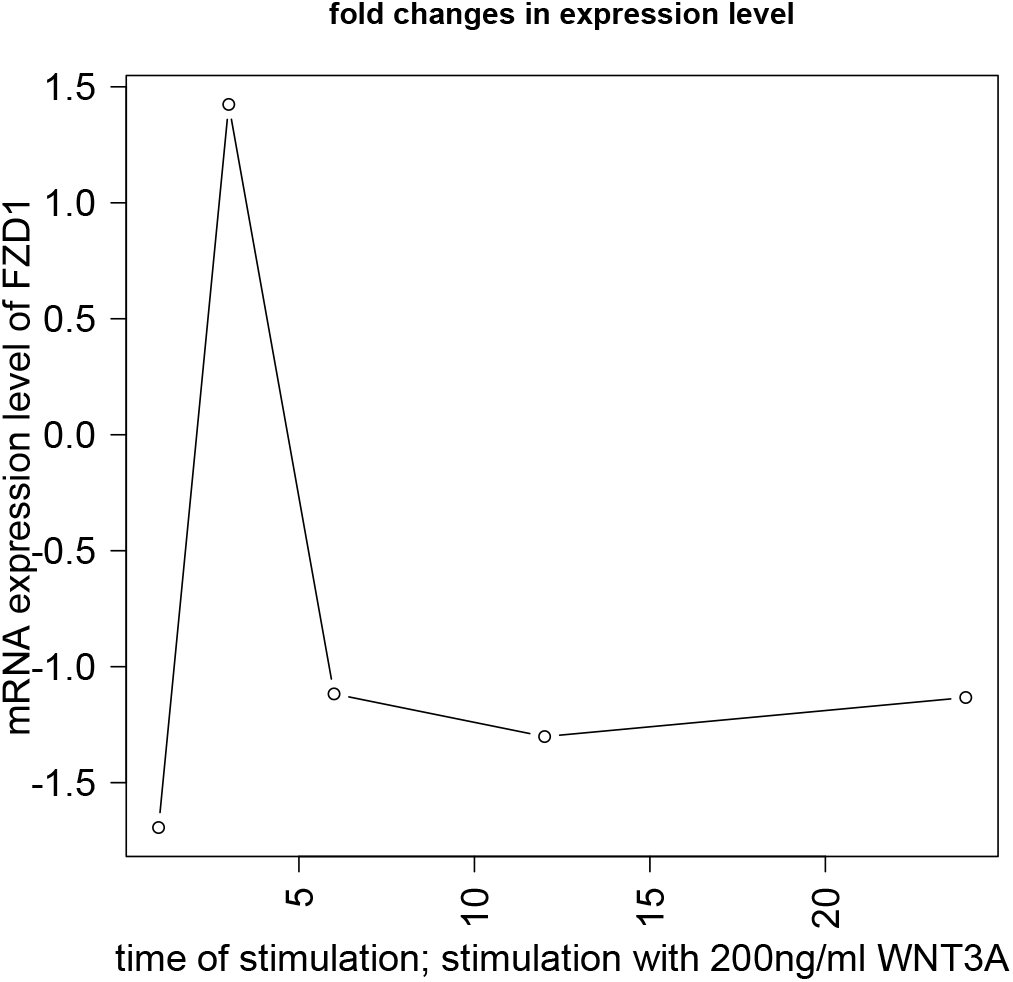
mRNA expression levels of *FZD*1 at 1^*st*^, 3^*rd*^, 6^*th*^, 12^*th*^ and 24^*th*^ hour from Gujral and MacBeath^17^.

